# The Preclinical Animal Network (PCAN): Integrative high-throughput phenotyping of standardized mouse models for Prader-Willi syndrome

**DOI:** 10.1101/2025.10.24.684371

**Authors:** Robert Wolff, Theresa V. Strong, Lisa C. Burnett, Nathalie Kayadjanian, James L. Resnick, Michelle Stewart, Sara E. Wells, Lydia Teboul, Alasdair J. Allan, Rasneer S. Bains, Alex Hardgrave, Anthony R. Isles, Françoise Muscatelli, Valter Tucci

## Abstract

The Preclinical Animal Network (PCAN) is a collaborative resource established to advance translational research and therapeutic development for Prader-Willi syndrome (PWS), a rare neurodevelopmental disorder caused by loss of paternally expressed genes on chromosome 15q11–q13. PCAN integrates standardized, high-throughput phenotyping of approximately 1,000 mice across six engineered lines carrying paternal deletions of *Ndn*, *Magel2*, *Snord116*, *Ipw*, and multigenic loci. Using validated experimental pipelines spanning metabolic, behavioral, and developmental domains, we captured comprehensive phenotypic profiles by applying a statistical variance-decomposition framework to partition genetic and environmental contributions. This analysis provides robust genotype–phenotype maps, revealing shared and model-specific phenotypic effects while accounting for key confounding factors. The PCAN resource provides a framework for evaluating PWS-relevant phenotypes and benchmarking therapeutic strategies. By providing open access to its mouse models, datasets and analytical resources, PCAN enables the research community to apply standardized, reproducible workflows and accelerate the development of targeted, mechanism-based treatments.

## Introduction

Prader-Willi syndrome (PWS) is a complex neurodevelopmental disorder caused by the loss of paternally expressed genes on chromosome 15q11.2–q13 (Fig. 1a). Clinically, PWS is characterized by neonatal hypotonia, developmental delay, intellectual disability, endocrine dysfunction, and hyperphagia, which often lead to severe obesity and related comorbidities if left unmanaged^1,2^. Individuals with PWS also commonly develop a range of behavioral and psychiatric problems^3,4^. Among the paternally expressed genes, *MAGEL2* is particularly notable: mutations in this gene cause Schaaf-Yang syndrome, a clinically overlapping neurodevelopmental disorder^5^.

**Fig. 1.**
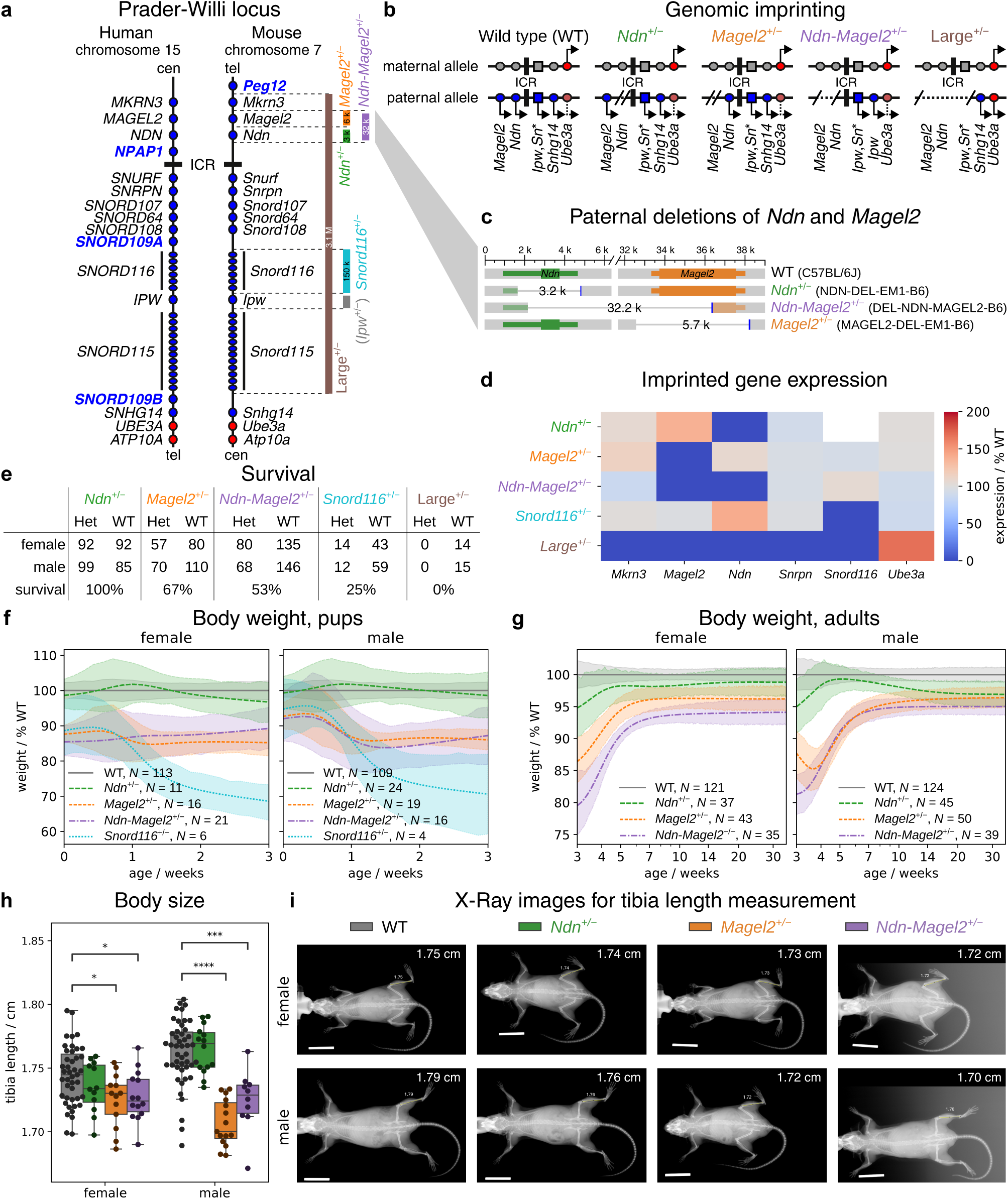
PWS mouse models in PCAN. **a**, Imprinted genes at the PW locus on human chromosome 15 and orthologues at mouse chromosome 7. The majority of genes are paternally expressed (blue dots). Here, only *UBE3A*/*Ube3a* and *ATP10A*/*Atp10a* are expressed with bias towards the maternal allele (red dots). The paternal deletions of the investigated mouse models are indicated at the right. ICR: imprinting control region. **b**, Modified allelic expression in the mouse models with paternal deletion due to genomic imprinting. **c**, Deletions in the mouse models for *Magel2*, *Ndn* and *Ndn*-*Magel2* double paternal deletion. **d**, Gene expression of imprinted genes at the PW locus for WT mice and mice with paternal deletion of single or multiple genes. **e**, Survival of animals at weaning for mice with paternal deletion (Het: heterozygous) and their WT littermates and estimated survival rates. **f**,**g**, Normalized body weight curves for pups (**f**) and adults (**g**). The line and upper and lower bounds of the shaded area correspond to the mean *±* 95% bootstrap confidence interval. **h**,**i**, Tibia length, as a proxy for body size, measured from X-ray images of animals at the age of around 26 weeks. Boxplot boxes in **h** extend from the first to the third quartile, with a line at the median and whiskers at the fartherst data points within the 1.5 *×* inter-quartile range. The bar in **i** indicates 2 cm.

Despite advances in understanding the genetic and molecular basis of PWS, therapeutic options remain scarce and primarily symptomatic. No approved treatments target the core behavioral or metabolic manifestations of the disorder. Rodent models have been instrumental in dissecting the roles of individual genes and mechanisms in PWS pathogenesis. However, their translational impact has been constrained by variability in genetic constructs, background strains, and phenotyping methods^6^. Models with large deletions encompassing the entire PWS locus often cause embryonic or perinatal lethality, whereas targeted knockouts of single genes typically produce partial phenotypes that fail to capture the full clinical spectrum^6,7^. A key remaining challenge is the limited standardization of phenotyping methodologies, which hampers consistent comparison across models and undermines reproducibility and translational relevance.

To address these challenges, the Foundation for Prader-Willi Research (FPWR) established the Preclinical Animal Network (PCAN). This collaborative initiative aims to generate, characterize, and share a robust panel of mouse models with targeted deletions of key PWS-associated genes. Here, we describe four viable mouse lines carrying paternal deletions of *Ndn*, *Magel2*, *Snord116*^8^, and a combined *Ndn*-*Magel2* deletion^9^ (Fig. 1a–c). A mouse line with a large deletion spanning multiple genes and the imprinting control region (ICR) of the PWS locus was also generated but proved non-viable (Fig. 1a,b). Attempts to generate a targeted deletion of *Ipw* were unsuccessful (Methods). All models were generated on a C57BL/6J background to ensure genetic consistency and facilitate direct comparisons.

The dataset comprises phenotyping outcomes from approximately 1,000 mice assessed across three high-throughput pipelines, representing one of the largest coordinated efforts to characterize mouse models for a rare disease. Model generation and optimization were conducted at the Medical Research Council (MRC), Mary Lyon Centre in Harwell, a key European hub for mouse genetics and functional genomics. The phenotyping pipelines build on two decades of European infrastructure development, including platforms established by the International Mouse Phenotyping Consortium (IMPC) and INFRAFRONTIER^10–13^. By implementing these standardized, validated experimental pipelines, PCAN ensures reproducibility of results and comparability of data across lines. These pipelines encompass metabolic, behavioral, and developmental domains with translational utility.

Equally critical to the success of this large-scale phenotyping effort is the implementation of rigorous analytical frameworks. Behavioral and physiological datasets are particularly susceptible to data-driven statistical artifacts, such as outcome selection bias and post-hoc hypothesis formulation, which can undermine reproducibility and inflate false discovery rates^14–16^. Moreover, subtle environmental variables, including cage density, handling protocols, and experimenter identity, can further confound genotype–phenotype associations, if not appropriately modeled^17^. Previous studies have emphasized the need to account for systematic environmental influences and individual-level variation to ensure reliable inferences in preclinical research^18^. However, these efforts have often been limited by small sample sizes, heterogeneous protocols, and insufficient control of confounding variables^19,20^.

To address these issues, we implemented a variance decomposition framework based on the ACE model, which partitions phenotypic variance into additive genetic (*A*), common environmental (*C*), and individual environmental (*E*) components^21–24^. This approach allows precise estimation of genotype effects across hundreds of phenotypic traits while accounting for non-genetic sources of variation. We further generated simulated “digital twins” using fitted ACE model parameters. These simulations disentangle the impact of genotype under controlled environmental conditions, assess robustness across sample sizes, and confirm the consistency of observed genotype–phenotype associations.

Using this integrated platform, we quantified the impact of individual and combinatorial gene deletions on a comprehensive set of morphological, developmental, metabolic, and behavioral phenotypes. The PCAN resource supports systematic cross-model comparisons, elucidates genotype-by-environment interactions, and distinguishes additive from non-additive genetic effects. Together, these data provide a high-resolution map of how discrete lesions within the PWS locus translate into specific biological and functional outcomes.

By making these models and data publicly available, PCAN provides a powerful, reproducible platform for preclinical research in PWS. The resource is designed to facilitate the identification of relevant endpoints, the development of targeted interventions, and the benchmarking of candidate therapies. As such, it serves as both a reference for mechanistic studies and a translational platform to advance therapeutic discovery. Building on this foundation, PCAN is entering a new phase focused on applying these standardized models to address specific clinical questions. Central to this effort is the systematic dissemination of phenotypic and analytical data, enabling both hypothesis-driven and exploratory investigations across academic and industry settings. Through the integration of validated experimental pipelines, rigorous statistical modeling, and large-scale phenotyping in a rare disease context, PCAN establishes a unique collaborative framework. This framework enables the PWS research community to adopt standardized, data-rich preclinical approaches that can accelerate the development of effective, mechanism-based therapies.

## Results

### Development of preclinical PWS mouse models

As part of the PCAN initiative, six mouse models were initially planned to target key paternally expressed genes within the PWS locus (Fig. 1a–c; Extended Data Fig. 1). Of these, four viable lines were successfully generated and maintained on a C57BL/6J genetic background: mice with paternal deletions of *Ndn* (*Ndn*^+/−^), *Magel2 (Magel2*^+/−^), *Snord116 (Snord116*^+/−^)^8^, and a combined *Ndn-Magel2 (Ndn-Magel2*^+/−^)^9^ deletion. Two additional models encountered major developmental constraints. A targeted knockout of *Ipw* (*Ipw*^+/−^) could not be successfully established, and a large deletion model (Large^+/−^) encompassing all the candidate PWS genes within the locus failed to survive beyond postnatal day one (P1), precluding its use in postnatal phenotyping studies. Furthermore, while the *Snord116* deletion line could be generated on the C57BL/6J background, it exhibited markedly reduced weight gain and neonatal viability, and could not be included in post-weaning phenotyping. Notably, the *Ndn*^+/−^ and *Magel2*^+/−^ mouse models have been constructed to allow for a Cre-dependent conditional knockout of *Ndn* and *Magel2*, respectively.

### Interdependence of gene expression within the PWS locus

We quantified the expression of the imprinted genes *Mkrn3*, *Magel2*, *Ndn*, *Snrpn*, *Snord116*, and *Ube3a* within the PWS locus using droplet digital PCR (ddPCR; Methods). Expression levels were normalized against the reference genes *Hprt1* and either *Gusb* or *Tbp*, and reported as percentages relative to wild-type (WT) controls (Supplementary Fig. 1; Methods). As expected, paternally expressed genes were not detectable in mouse lines carrying paternal single- or multi-gene deletions (Fig. 1b,d; Supplementary Fig. 2a–e). Specifically, *Ndn* was absent in the *Ndn*^+/−^, *Ndn-Magel2*^+/−^, and Large^+/−^ models, while *Magel2* was absent in the *Magel2*^+/−^, *Ndn-Magel2*^+/−^, and Large^+/−^ models. Notably, *Ube3a* expression was elevated in the Large^+/−^ model, which carries a deletion encompassing the imprinting control region (ICR), but remained unchanged relative to WT in the other models (Fig. 1b,d; Supplementary Fig. 2f). This is consistent with prior reports of *Ube3a* being expressed with bias towards the maternal allele^25,26^. We verified that mice with equivalent maternal deletions (e.g., *Ndn*^+/−^, Large^+/−^) or with floxed maternal or paternal alleles revealed no alterations in gene expression compared to WT (Supplementary Fig. 2). ELISA-based protein measurements of *Neu2*, *Oxt* and *Igf1*, which may be linked to PWS phenotypes^27–32^, in *Ndn*^+/−^ and *Ndn-Magel2*^+/−^ animals at end of life showed no differences relative to their WT littermates (Supplementary Fig. 3). Furthermore, we measured hematology for *Ndn-Magel2*^+/−^ animals and found no differences to their WT littermates (Supplementary Fig. 4). Together, these data confirm that gene expression within the PWS locus is tightly regulated by imprinting, with predictable loss upon paternal deletion and selective dysregulation only in models disrupting the imprinting control region.

### Paternal deletion of imprinted genes reduces survival

We quantified survival to weaning across the different paternal deletion lines relative to wild-type (WT) littermates (Fig. 1e). Single paternal deletion of *Ndn* did not affect survival, with *Ndn*^+/−^ mice reaching weaning at rates comparable to WT. In contrast, survival was reduced in *Magel2*^+/−^ mice (67%; 95% CI: 60–73%) and further decreased in *Ndn-Magel2*^+/−^ mice (53%; 95% CI: 47–59%), confronting previous contradictory findings^33–39^. Only 25% of *Snord116*^+/−^ mice (95% CI: 18–35%) survived to weaning, while Large^+/−^ mice exhibited perinatal lethality*Snord116*^+/−^. Across all models, females exhibited a non-significant (*p* > 0.05) trend towards better survival than males. Collectively, these findings demonstrate that paternal deletion of key imprinted genes within the PWS locus differentially impacts postnatal viability.

### Paternal deletion of *Ndn*, *Magel2*, and *Snord116* alters growth and body composition

We tracked pup body weight daily during the first two to three postnatal weeks and measured adult weights weekly to bi-weekly thereafter. Individual growth trajectories were modeled using a custom regression function resembling a modified inverse tangent, which captures asymptotic adult weight and the inflection point of maximal weight gain (Extended Data Fig. 2a,b). Estimated weights were used to plot growth curves for each genotype relative to wild-type (WT) littermates. At birth (P0), *Magel2*^+/−^, *Ndn-Magel2*^+/−^, and *Snord116*^+/−^ pups displayed reduced body weights of 85–95% of WT littermates, whereas *Ndn*^+/−^ pups did not (Fig. 1f, Extended Data Fig. 2c). This reduction was persistent throughout development for *Magel2*^+/−^ and *Ndn-Magel2*^+/−^ pups. In contrast, surviving *Snord116*^+/−^ pups showed a progressive reduction in WT-relative body weight after P4, reaching 70–75% of WT by P14 (Fig. 1f; Extended Data Fig. 2d), which manifests itself in reduced age at maximum gain (Extended Data Fig. 2e). The reduction in maximum gain for *Magel2*^+/−^, *Ndn-Magel2*^+/−^, and *Snord116*^+/−^ pups reflects the reduced body weights (Extended Data Fig. 2f). For ethical reasons, *Snord116*^+/−^ mice were not retained for later experiments. After weaning, *Magel2*^+/−^ and *Ndn-Magel2*^+/−^ mice exhibited delayed growth, but reached near-WT weights by 6 weeks (Fig. 1g; Extended Data Fig. 2g,h). The age of maximum gain was increased in *Magel2*^+/−^ and *Ndn-Magel2*^+/−^ mice compared to their WT littermates but the maximum gain is not anymore significantly reduced (Extended Data Fig. 2i,j), suggesting that during 2 weeks after weaning the mutant mice quickly gained weight but did not reach the weight of the controls (Fig. 1g). Body size was assessed by tibia length at 26 weeks and a reduction of length was observed in *Magel2*^+/−^ and *Ndn-Magel2*^+/−^ mice suggesting a reduction in body size, consistent with the reduced body weight (Fig. 1h,i; Extended Data Fig. 3a,b). Body composition assessed by MRI-NMR and X-ray DEXA suggested a trend toward increased fat mass with reduced lean and water mass in *Ndn*^+/−^ mice, whereas *Magel2*^+/−^ and *Ndn-Magel2*^+/−^ mice showed the opposite trend, with reduced fat mass and increased lean mass (Extended Data Fig. 3d–g). Altogether, these data showed an alteration of body weight and body composition in all mutant mice with a different effect in *Ndn*^+/−^ mice compared with the three other mouse lines.

### Phenotyping pipelines and variance decomposition

We performed phenotyping experiments on *Ndn*^+/−^, *Magel2*^+/−^, and *Ndn-Magel2*^+/−^ mice and their wild-type littermates across three independent pipelines (*I*, preweaning and nociception; *II*, metabolic; and *III*, behavioral), each comprising a separate cohort (Fig. 2a; Table 1; Methods). All experiments were performed at MLC Harwell with an effort for minimizing environmental variance. Experimental outputs were refined into standardized phenotypic variables (Fig. 2b). For example, in the passive infrared (PIR) assay, circadian rhythm parameters were derived from raw activity data binned at 10-second intervals (Extended Data Fig. 4). Power-transformed variables were then subjected to principal component analysis (PCA) separately for males and females. Each resulting principal component (“observable”) was analyzed with the ACE model to partition variance into additive genetic effects (*A*), common environmental influences (*C*), and individual-specific noise (*E*). Here, the additive genetic effects were attributed to paternal gene deletions, where the *Ndn-Magel2*^+/−^ mice have both paternal deletions of *Ndn* and *Magel2* plus an additional term used to check the non-additivity of the single deletions. We considered the metadata, collected together with the phenotypic outcomes, for various quantitative and qualitative covariates of common environmental effects, such as the age of the animals at the test, their body weight, their batch of origin, the year and month when the experiment was performed, and the identifier of the experimenter who performed the test (Extended Data Table 1; Methods). This framework enabled estimation of deletion effect sizes and provided AE-adjusted values that exclude common environmental confounds for downstream analyses (Methods).

**Fig. 2.**
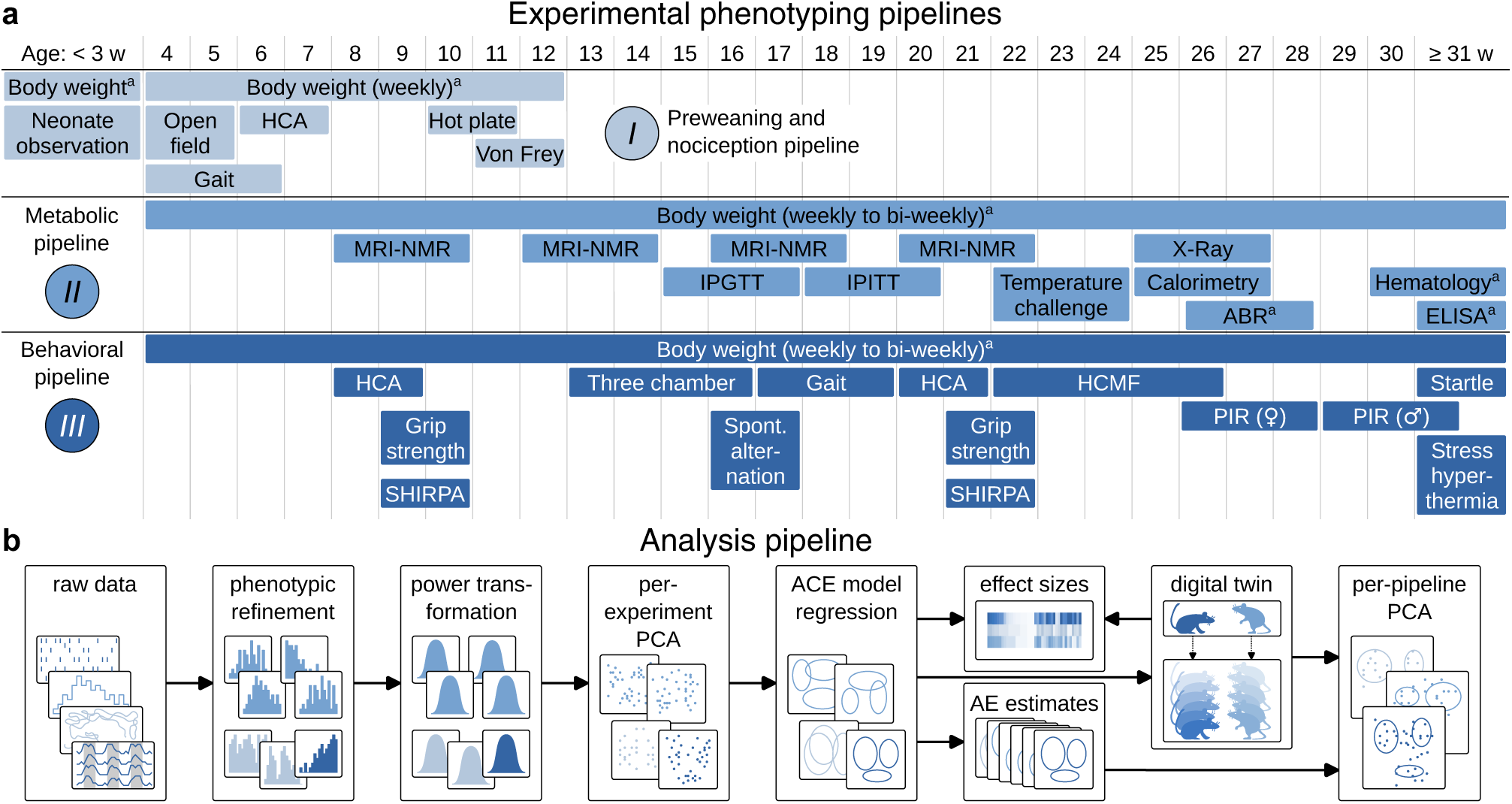
Scheme of experiments and analysis. **a**, Experimental phenotyping pipelines (*I*, *II* and *III*) with experimental tests and approximate age (in weeks) of animals when we performed these tests. HCA: Home cage analyzer; MRI-NMR: Magnetic resonance imaging by nuclear magnetic resonance; IPGTT and IPITT: Intraperitoneal tolerance test for glucose and insulin, respectively; ABR: Auditory brain stem response; ELISA: Enzyme-linked immunosorbent assay; SHIRPA: Combined Shirpa and Dysmorphology; HCMF: Home-cage motor function; PIR: Passive infrared; Startle: Acoustic startle. **b**, Analysis pipeline. PCA: principal component analysis; ACE: regression model with additive genetic effect (*A*), common environment (*C*) and unique environment (*E*); AE estimates: estimates removing the common environment. *^a^*) Body weights and ABR, hematology and ELISA experiments are not included in the final per-pipeline PCAs.

**Table 1.** Experiments in three pipelines and numbers of tested WT and heterozygous animals.

| Pipeline | Experiment | Number of animals |  |  |  | Age / d |
| --- | --- | --- | --- | --- | --- | --- |
|  |  | WT | *Ndn*<sup>+/-</sup> | *Magel2*<sup>+/-</sup> | *Ndn-Magel2*<sup>+/-</sup> |  |
| Prewaning and nociception (pipeline I) | Neonate observation* | 224 | 35 | 35 | 37 | 0–21 |
|  | Body weight (pups)<sup>†‡</sup> | 222 | 35 | 35 | 37 | 0–21 |
|  | Open field | 89 | 24 | 24 | 41 | 28 ± 3 |
|  | Gait | 85 | 22 | 23 | 35 | 29 ± 5 |
|  | Home-cage activity (HCA) | 65 | 23 | 23 | 19 | 44 ± 3 |
|  | Hot plate | 77 | 23 | 24 | 21 | 72 ± 3 |
|  | Von Frey | 77 | 23 | 24 | 21 | 78 ± 3 |
|  | Body weight (adults)<sup>†</sup> | 51 | 23 | 24 | 3 | – |
| Metabolic (pipeline II) | Body composition (MRI-NMR) | 95 | 29 | 37 | 30 | 63 ± 5 |
|  |  | 95 | 29 | 36 | 29 | 89 ± 4 |
|  |  | 95 | 29 | 32 | 29 | 117 ± 4 |
|  |  | 95 | 27 | 32 | 29 | 146 ± 4 |
|  | Intraperitoneal glucose tolerance test (IPGTT) | 95 | 29 | 34 | 28 | 111 ± 4 |
|  | Intraperitoneal insulin tolerance test (IPITT) | 83 | 20 | 30 | 29 | 127 ± 5 |
|  | Temperature challenge | 95 | 27 | 33 | 29 | 154 ± 4 |
|  | Calorimetry | 79 | 27 | 33 | 13 | 179 ± 11 |
|  | X-ray | 90 | 26 | 28 | 24 | 182 ± 4 |
|  | Body composition (X-Ray) | 91 | 27 | 33 | 26 | 182 ± 4 |
|  | Auditory brainstem response (ABR) | 15 | 8 | 4 | 3 | 184 ± 5 |
|  | Hematology | 29 | –<sup>§</sup> | –<sup>§</sup> | 25 | 215 ± 6 |
|  | ELISA | 59 | 30 | –<sup>§</sup> | 26 | – |
|  | Body weight (adults)<sup>†</sup> | 95 | 29 | 38 | 30 | – |
| Behavioral (pipeline III) | Home-cage activity (HCA) | 97 | 24 | 29 | 38 | 58 ± 2 |
|  |  | 93 | 26 | 27 | 29 | 142 ± 3 |
|  | Combined Shirpa and dysmorphology (SHIRPA) | 102 | 30 | 31 | 41 | 63 ± 2 |
|  |  | 102 | 30 | 29 | 36 | 148 ± 2 |
|  | Grip strength | 102 | 30 | 31 | 41 | 66 ± 3 |
|  |  | 101 | 30 | 29 | 34 | 150 ± 2 |
|  | Three chamber | 88 | 21 | 24 | 26 | 100 ± 7 |
|  | Spontaneous alternation | 90 | 30 | 29 | 32 | 113 ± 2 |
|  | Gait | 92 | 29 | 20 | 40 | 120 ± 3 |
|  | Home-cage motor function (HCMF) | 87 | 28 | 29 | 25 | 164 ± 10 |
|  | Passive infrared (PIR) | 70 | 24 | 15 | 26 | 196 ± 13 |
|  | Stress induced hyperthermia | 89 | 30 | 28 | 26 | 224 ± 6 |
|  | Acoustic startle | 84 | 30 | 28 | 26 | 233 ± 14 |
|  | Body weight (adults)<sup>†</sup> | 102 | 30 | 31 | 41 | – |

### Environmental and genetic contributions to phenotypic variance

Across most phenotypic outcomes, variance was primarily explained by individual-specific effects (*E*), followed by common environmental factors (*C*) captured through model covariates (Fig. 3a–d, Extended Data Fig. 6). Genetic contributions (*A*) from paternal deletions were smaller overall but detectable for selected traits. We observed similar environment effect sizes for heterozygous mice and their WT littermates (Extended Data Fig. 6c). Moreover, we observed correlations between covariates and the genetic deletion of similar size as of the genetic deletion effect size. In particular the batch, the experimenter and the date of experiment were correlated and had similar effect size as of the genetic deletion in many experiments (Supplementary Fig. 5–7). Among the deletions, *Ndn*^+/−^ had the weakest impact, whereas *Magel2*^+/−^ and *Ndn-Magel2*^+/−^ affected more outcomes. Within ACE model regression, we estimated *p*-values for non-zero paternal deletions of *Ndn* and *Magel2* (Fig. 3e). Consistent with the variance analysis, *Magel2* deletion influenced a broader range of the phenotypes than *Ndn* deletion. Together, these results indicate that while environmental factors account for the majority of phenotypic variance, paternal *Magel2* deletion drives clear functional deficits across multiple traits.

**Fig. 3.**
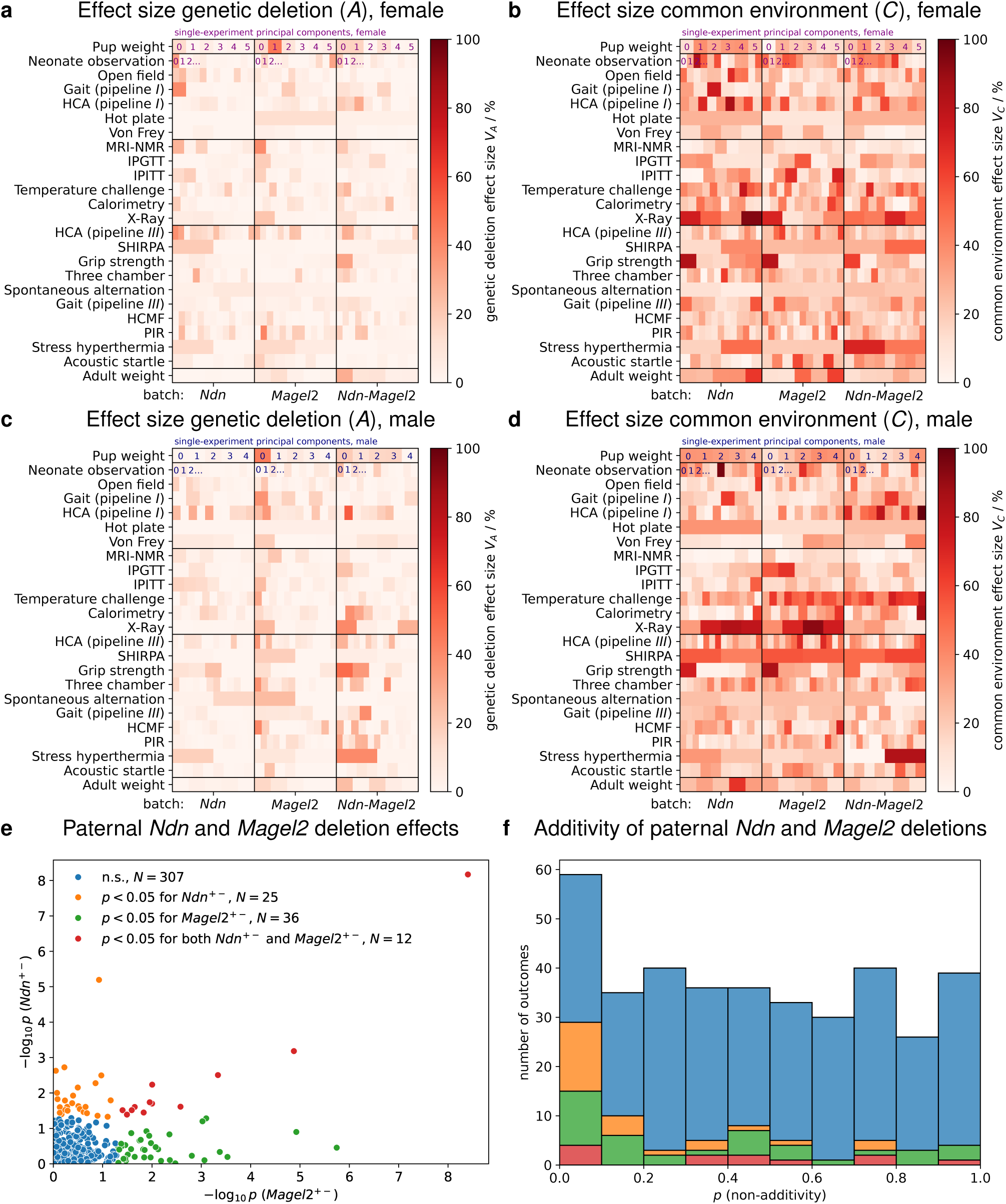
Effect sizes of paternal *Ndn* and *Magel2* deletion and environmental effects. **a**,**c**, Effect sizes by genetic deletion for females (**a**) and males (**c**). **b**,**d**, Effect sizes by common environment for females (**b**) and males (**d**). Each data point in **a**–**d** corresponds to one single-experiment principal component, sorted from left to right by highest explained variance. The female and male *Ndn* batches include heterozygous *Ndn*^+^*^/−^* animals and their WT littermates, and *Magel2* and *Ndn*-*Magel2* batches are defined analogously. **e**, Observed *p*-values for testing the null-hypothesis of no effect of paternal *Ndn* or *Magel2* deletion. *N* = 380 outcomes (PCs). **f**, Observed *p*-values testing the null-hypothesis of additivity of paternal *Ndn* and *Magel2* deletion effects.

### Additive effects of paternal *Ndn* and *Magel2* deletions and validation with digital twins

Analysis of mice carrying combined *Ndn* and *Magel2* deletions allowed us to test whether the two single deletions act additively. The distribution of *p*-values for deviation from additivity was nearly uniform, with only a few low *p*-values, indicating that for most traits the effects of paternal *Ndn* and *Magel2* deletions are additive (Fig. 3f).

We next used the ACE model estimates to generate simulated “digital twins” (Fig. 4a). In these simulations, additive genetic effects (A) were modeled with or without paternal deletion, common environmental effects (C) were sampled from observed covariate distributions, and residual variance (E) was drawn from the regression residuals. For example, in female adult weight (PC1), the simulated impact of *Magel2* deletion closely matched the observed regression effect, whereas simulations with no *Magel2* effect produced values centered around zero (Fig. 4b). Across all traits, simulated and observed *p*-values were highly correlated for both *Ndn* and *Magel2* deletions, with only a few observed low *p*-values (< 0.05) not reproduced in simulation (Fig. 4c,d).

**Fig. 4.**
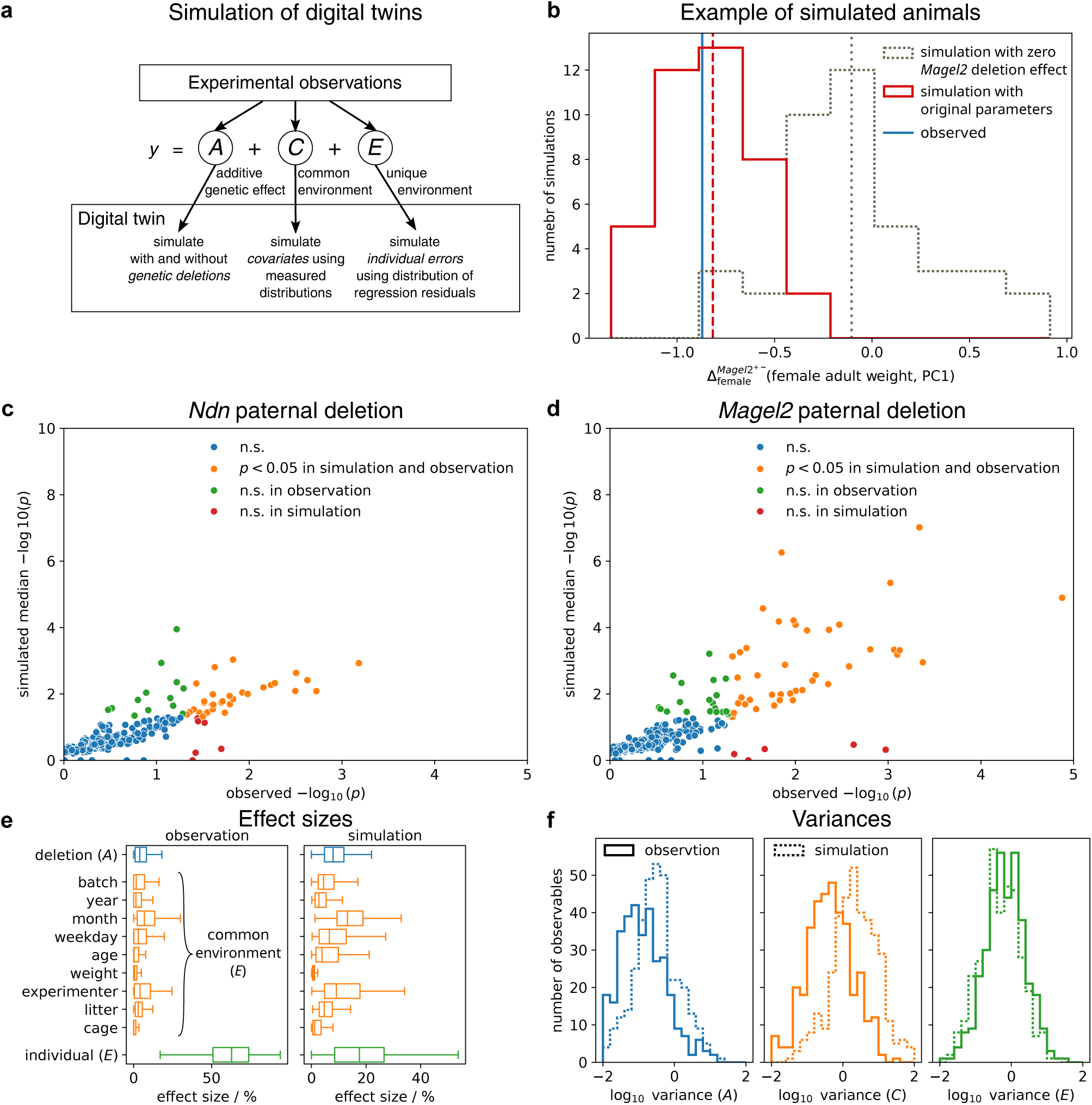
Simulation of digital twins. **a**, Schematic overview of the simulation. **b**, Example of simulations with original parameters using the measured female adult weight 2nd principal component (PC1) with and without the effect of paternal *Magel2* deletion. *N* = 40 simulations in both settings. **c**,**d**, *p*-values for observables (PCs) in observation (*x*-axis) and simulation (*y*-axis) for the effect of paternal *Ndn* (**c**) and *Magel2* (**d**) deletion. **e**, Effect sizes (relative variances) in observation (left) and simulation (right) by paternal deletion (blue), common environment (orange) and individual, unique environment (green). **f**, Absolute variances (effect sizes) in observation (solid line) and simulation (dashed line). Simulated *p*-values in **c**,**d**, simulated effect sizes in **e** and simulated variances **f** have been calculated from the median values on *N* = 40 simulations. Boxplot boxes extend from the first to the third quartile, with a line at the median and whiskers at the fartherst data points within the 1.5 *×* inter-quartile range.

Comparison of relative effect sizes between ACE model outputs and simulations showed consistent relationships among subsubcomponents of the common environment, although absolute values of both *A* and *C* were overestimated in simulations (Fig. 4e,f). This discrepancy likely reflects unmodeled correlations among covariates.

### Paternal *Ndn* and *Magel2* deletions preferentially affect behavioral outcomes

The behavioral pipeline included assays of locomotion, social interaction, gait, grip strength, circadian rhythm, and sensory responsiveness, providing an integrated assessment of neuromotor coordination, activity, and social behavior. The metabolic pipeline comprised measurements of body composition, calorimetry, glucose and insulin tolerance, thermoregulation, and hematological profiles, capturing systemic metabolic homeostasis. To assess cross-experiment effects, we performed PCAs that combined all experiments within each pipeline, using AE estimates from the single-experiment analyses as input. This revealed that mutant mice were most clearly separated from WT by the “behavioral” pipeline (Fig. 5a), indicating that paternal *Ndn* and *Magel2* deletions primarily disrupt neuromotor and behavioral domains, consistent with known functions of these genes in neuronal development and synaptic regulation. Differences were also detectable in the “metabolic” pipeline, particularly among males, whereas the “preweaning and nociception” pipeline showed no separation between groups. Effects observed in the metabolic pipeline suggest physiological consequences, such as altered energy balance and thermogenic control. Together, these findings reveal that paternal loss of *Ndn* and *Magel2* exerts its strongest impact on neurobehavioral traits, while also inducing metabolic alterations, thereby recapitulating key features of PWS pathophysiology.

**Fig. 5.**
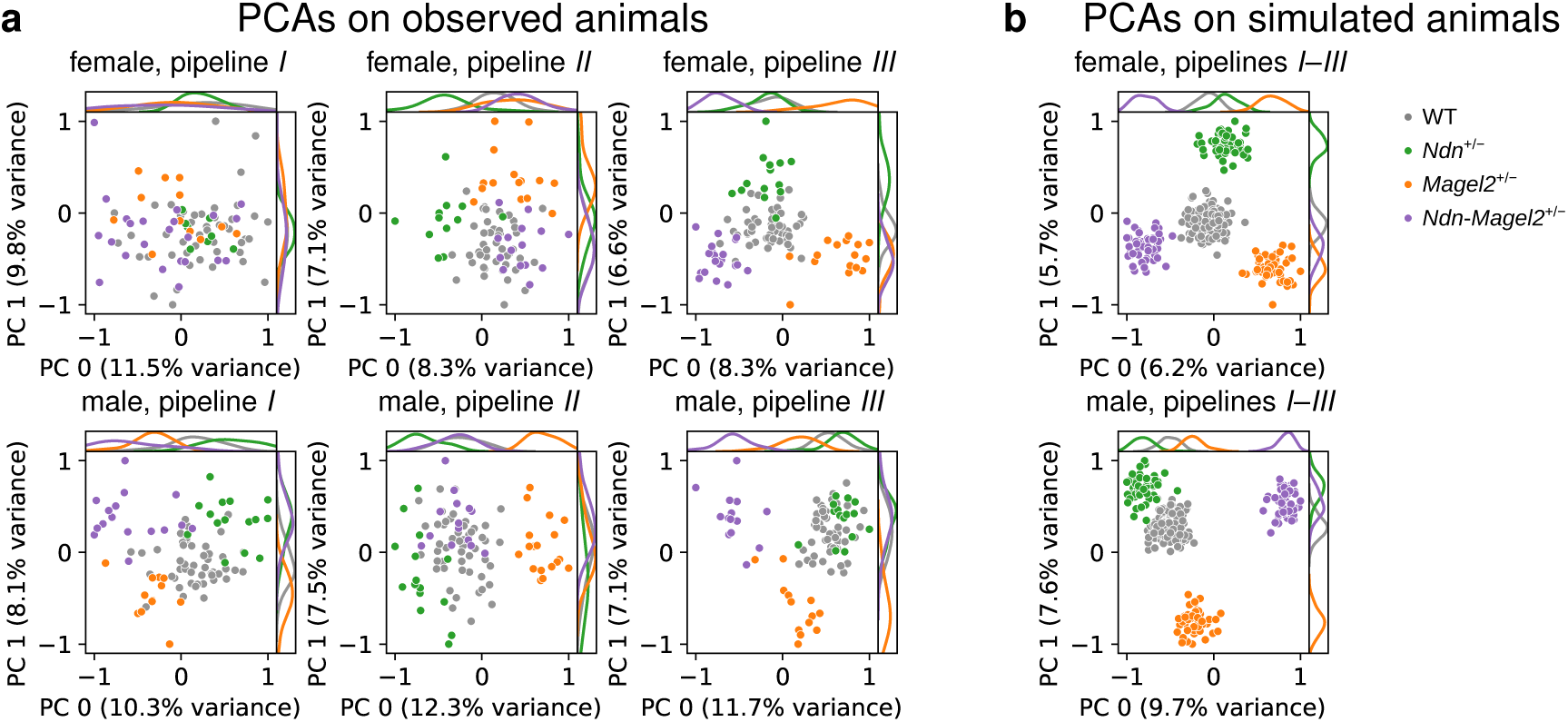
Principal component analysis (PCA) on observed and simulated animals. **a**, PCAs for the three pipelines for observations of females (top) and males (bottom) from the preweaning and nociception pipeline (pipeline *I*, left), the metabolic pipeline (pipeline *II*, middle) and the behavioral pipeline (pipeline *III*, right). **b**, PCAs on simulated females (top) and males (bottom) in one example simulation with *N* = 50 animals per group (6 groups: *Ndn*^+^*^/−^*, *Magel2*^+^*^/−^* and *Ndn*-*Magel2*^+^*^/−^*, and their respective WT-littermate groups).

Simulations with 50 digital animals per group provided further confirmation (Fig. 5b). Because simulated animals could be assigned outcomes across all experiments, we did not need to stratify by pipeline. In this setting, group separation was even more pronounced than in the observed data, consistent with the larger fraction of p-values < 0.05 obtained from simulations (see Fig. 4c,d; Supplementary Fig. 26). Combining all experiments across pipelines in the simulations further improved discrimination between genotypes.

Importantly, varying the number of simulated animals or removing common environment effects produced only minor changes in the PCA results, underscoring the robustness of the approach and reflecting the use of AE estimates that already exclude shared environmental variance. The digital twin simulations thus provide an independent and scalable validation of the experimental findings, demonstrating that the observed genotype effects are robust to environmental variation and can be reliably reproduced even under conditions not directly accessible *in vivo*.

## Discussion

Much of the inconsistency in PWS preclinical models stems from the use of diverse mouse lines and genetic backgrounds, as well as laboratory-specific protocols for generation and phenotyping. Together, these factors hinder reproducibility, interpretation, and clinical translation. The Preclinical Animal Network (PCAN) directly addresses these issues by establishing a unified and standardized platform for model generation, phenotyping, and analysis. Through systematic characterization of approximately 1,000 mice, comprising paternal-deletion mutants and their wild-type littermates, on a uniform C57BL/6J background, PCAN provides a comprehensive and openly accessible resource. Notably, the *Ndn* and *Magel2* mouse lines were engineered as conditional knockouts, allowing gene deletion in specific tissues or developmental stages. This design enables targeted investigations of spatially and temporally restricted gene functions, supporting future studies aimed at dissecting mechanistic pathways underlying PWS phenotypes. The integration of variance decomposition using the ACE model and simulation-based validation via digital twins extends this dataset beyond descriptive phenotyping, providing a scalable analytical foundation for hypothesis testing and therapeutic benchmarking. Rather than another isolated dataset, PCAN represents a methodological and organizational framework for how preclinical resources in rare genetic disorders can be designed and shared.

One of the main strengths of PCAN lies in its ability to directly compare the phenotypic consequences of single and combined gene deletions within a common experimental and analytical framework. These comparisons reveal both convergent and distinct roles for *Ndn*, *Magel2*, and *Snord116*, establishing a modular view of the PWS locus. Survival analyses illustrate this principle: while paternal *Ndn* deletion alone minimally affected early viability, *Magel2* deletion reduced survival to two-thirds of expected levels, and the combined *Ndn*-*Magel2* deletion to about half. *Snord116* deletion caused more severe early lethality, and the large multigenic deletion covering these loci and the intervening ICR proved non-viable. These results, consistent with previous reports^33–39^, now stand within a standardized and quantitative context. Importantly, the *Ndn*-*Magel2* deletion model revealed additive effects on survival, with limited evidence for non-linear interactions, demonstrating that distinct imprinted genes within the PWS region contribute modularly to the phenotype. This insight may inform therapeutic strategies targeting subsets of molecular pathways.

Growth and body composition outcomes further delineate the phenotypic manifestations of these deletions. *Magel2* and *Ndn-Magel2* mutants displayed consistent reductions in birth weight and partial catch-up growth after weaning, whereas *Snord116* mutants exhibited progressive and irreversible growth impairment, leading to early termination of the experiments for ethical reasons. *Ndn* mutants grew comparably to wild type but showed subtle alterations in body composition, including a relative increase in fat mass. These developmental trajectories reflect the clinical heterogeneity in individuals with PWS: many patients show early growth failure followed by later obesity, while others display more restricted phenotypes^40–43^. By anchoring these dynamics in gene-specific contributions, PCAN provides a roadmap for disentangling the molecular basis of metabolic and growth abnormalities in PWS.

After somatic traits, behavioral phenotypes emerged as the most sensitive and discriminative indicators of PWS-related paternal gene deletions. Across experimental pipelines, behavioral assays consistently distinguished mutant from wild-type groups, with *Magel2* deletion producing deficits in activity and sleep-like patterns. These findings resonate with clinical experience in PWS, where neurobehavioral manifestations are among the most penetrant and therapeutically challenging features^44^. While the presented data were obtained under tightly controlled laboratory conditions, additional discriminative phenotypes may emerge under environmental stress or pathogen exposure. Under such conditions, Mage genes, such as *Magel2* and *Ndn*, are known to modulate adaptive responses^45^. For instance, survival of *Ndn*-deficient mice is variable and depends on respiratory deficits that become apparent when environmental pathogens are present, emphasizing that future studies incorporating environmental challenges will refine the translational relevance of these models.

A key methodological contribution of PCAN is the implementation of the ACE variance decomposition model to partition phenotypic variance into additive genetic, common environmental, and individual-specific components. Preclinical studies often suffer from uncontrolled environmental noise, such as litter effects, experimenter differences, seasonal changes, and other factors, that obscure genuine genotype–phenotype associations. By explicitly incorporating these structured covariates, the ACE model provides refined estimates of genetic contributions while quantifying both common and individual environmental influences. Across pipelines, wild-type cohorts displayed minimal dispersion, confirming the technical robustness and reproducibility of the standardized phenotyping platform. Individual-specific environmental variance accounted for the largest share of total variance across most traits, while much of the systematic variation was captured within the modeled common-environmental component, indicating that environmental effects were both controlled and quantifiable. Despite being smaller in magnitude, consistent genetic effects across traits and pipelines demonstrated that true genetic contributions were robustly detected in mutants. *Magel2* deletion produced broad effects across growth and behavioral domains, whereas *Ndn* deletion contributed more narrowly. The combined *Ndn-Magel2* deletion showed largely additive effects, again underscoring the modular architecture of the PWS locus. By quantifying and adjusting for environmental variance, PCAN ensures that genotype–phenotype associations are not confounded by laboratory-specific conditions, a significant advance in a field often challenged by reproducibility issues^14–19^. Beyond PCAN, large-scale infrastructures such as the IMPC offer complementary platforms to validate and extend this variance-decomposition approach across diverse genetic backgrounds and experimental settings.

The introduction of digital twin simulations represents another major advance of PCAN. Using ACE-model-derived distributions, we generated synthetic cohorts of virtual animals to test hypotheses under conditions not directly accessible *in vivo* and to validate the robustness of observed effects. Simulated animals reproduced the key genetic effects seen *in vivo*, confirming their robustness against environmental variation, and showed that increasing sample sizes or combining outcomes across pipelines enhances statistical power without additional animal use. Discrepancies between observed and simulated effect sizes highlighted the impact of unmodeled correlations among covariates and between covariates and genotype on variance estimates, suggesting directions for further refinement. Beyond validation, digital twins embody the principles of the 3Rs in animal research (replacement, reduction, refinement) by extending inference without increasing animal numbers. Moreover, by providing open access to raw data and simulation tools, PCAN democratizes research: any laboratory can generate digital twins to test hypotheses, benchmark interventions, or explore gene–environment interactions. This accessibility also enables the development and testing of analytical models without *a priori* hypotheses, supporting unbiased exploration of genotype–phenotype relationships and transforming PCAN from a static dataset into a dynamic, evolving platform.

The ultimate goal of PCAN is to accelerate translational progress in PWS. Several directions emerge from the resource. First, PCAN establishes standardized phenotypic benchmarks for preclinical testing. Behavioral outcomes, validated as sensitive endpoints, can serve as primary readouts in therapeutic trials, while growth and body composition provide complementary secondary metrics. This framework ensures that interventions are evaluated against comparable standards across laboratories. Second, PCAN clarifies gene-specific contributions, informing therapeutic strategies that may need to target multiple pathways. The additive nature of *Ndn* and *Magel2* effects suggests that combination approaches involving pharmacological, genetic, or epigenetic interventions may be required to achieve meaningful clinical benefit. Third, the variance decomposition framework distinguishes true therapeutic effects from environmental noise, improving reproducibility and translational fidelity, which are both essential for advancing therapeutic candidates toward clinical trials. Finally, the digital twin platform offers a bridge to personalized and predictive medicine. By simulating large, diverse cohorts under varying environmental conditions, digital twins can forecast how interventions might perform across heterogeneous patient populations. Over time, this approach could integrate molecular and omics-level data to create multi-scale simulations that more closely reflect the complexity of human biology.

Beyond PWS, PCAN serves as a proof of concept for collaborative, standardized resource development in rare disease research. Funded by the FPWR, the initiative combined large-scale mouse colony generation and phenotyping at the MRC Harwell Institute with strategic guidance from an international panel of experts who defined the genetic backgrounds, mutations, and phenotypes to prioritize. The scientific program was coordinated by three dedicated PWS laboratories, and data analyses were performed independently to ensure objectivity. By integrating this organizational framework with standardized pipelines, robust statistical modeling, and digital twin simulations, PCAN provides a new template for generating and disseminating preclinical resources. Open accessibility to its data and tools empowers the research community to drive discovery, validation, and innovation. This model of collaborative, data-rich resource building has broad implications, with the potential to accelerate progress in other imprinting disorders, neurodevelopmental syndromes, and even complex polygenic traits.

Overall, PCAN establishes a comprehensive, validated resource for preclinical and translational research in PWS. The standardized experimental and analytical framework, combined with digital twin simulations, confirms the robustness of behavioral and somatic phenotypes across the analyzed PWS mouse models and provides a foundation for reproducible benchmarking of therapeutic interventions. The inability of certain deletions to support survival on the C57BL/6J background underscores the importance of accounting for both genetic and epigenetic context in preclinical model development. Variability in developmental, metabolic, and behavioral outcomes across strains reflects the influence of background-specific modifiers that shape the penetrance and expressivity of PWS-related phenotypes^46^. As such, these observations not only inform model selection but also provide a framework for understanding the broader regulatory landscape of the PWS locus. By coupling methodological rigor with open accessibility, PCAN advances the development of mechanism-based therapies for PWS and establishes a scalable paradigm for collaborative, data-driven discovery in rare and complex genetic disorders.

## Methods

### Mouse lines

Three new PWS mouse lines with paternal gene deletions were created using CRISPR-Cas9 technology on C57BL/6J background (Fig. 1; Extended Data Fig. 1): *Ndn* (*Ndn*^+/−^, NDN-DEL-EM1-B6), *Magel2* (*Magel2*^+/−^, MAGEL2-DEL-EM1-B6), and a large deletion (Large^+/−^, PCAN-DEL3Mbp-EM4-B6). Initially, floxed alleles of both *Ndn* and *Magel2* were generated separately by pronuclear injection of CRISPR/Cas9 reagents, including a long single stranded donor template, into single cell embryos (see Supplementary information for sequence and design details). The null alleles of *Ndn* and *Magel2* were then generated by carrying out an in vitro fertilization (IVF) using *Ndn*^m:flox/p+^ (NDN-FLOX-EM1-B6) or *Magel2*^m:flox/p+^ (MAGEL2-FLOX-EM1-B6) females, respectively. Heterozygous females underwent superovulation by injection of pregnant mares serum (5 iu per female), followed 48 hours later by injection of human chorionic gonadotrophin (5 iu per female). 16 hours later oocytes were harvested and IVF carried out using C57BL/6J sperm. Soluble cell-permeable Cre (TAT-Cre (Tat-NLS-Cre, HTNC, HTNCre), Excellegen, Rockville, MD, USA) was added to two cell embryos. The soluble Cre excised the DNA between the loxP sites, generating a null allele of each line. Following washing to remove the soluble Cre, the IVF procedure was completed as normal. Resulting embryos were genotyped and females carrying the deletions were crossed to C57BL/6J males to expand the colony and for colony maintenance. We also generated a line with a 3 Mbp large deletion of a portion of the PWS region on mouse chromosome 7, from 2,820 bp upstream to *Mkrn3* until 933 bp downstream of *Ube3a* (Large^m–/p+^, PCAN-DEL-B6, Supplementary information). The generation of a fourth mouse line with deletion of *Ipw* (*Ipw*^m−/p+^) was not successful. The region to flox with was on the large side for genome editing at the time with a donor of 2.6 kbp. We screened over 400 G0s and 60 showed evidence of guide cutting but there was no correct integration of the donor on target, although some animals were obtained with recombined segments and evidence of at least integration of one loxP. Mice with *Snord116* deletion were created using mice with floxed *Snord116* (B6.Cg-*Snord116*^tm1Uta^/J, https://jax.org/strain/008118)^8^.

### Animals

All animals were generated, housed, maintained and tested in the Mary Lyon Centre (MLC) at MRC Harwell under specific pathogen-free conditions, in individually ventilated cages (IVCs) adhering to environmental conditions as outlined in the Home Office Code of Practice. All animal studies were licensed by the Home Office under the Animals (Scientific Procedures) Act 1986 Amendment Regulations 2012 (SI 4 2012/3039), UK, and additionally approved by the Institutional Ethical Review Committees. Mice were randomized, blocked by genotype and sex at the time of weaning (at the age of 20 to 22 days), into cages of three mice. All mice used in the study were bred in the MLC and were housed in IVCs (Tecniplast BlueLine 1284), on grade 4 aspen wood chips (Datesand, UK), with shredded paper shaving nesting material and small cardboard play tunnels for enrichment. The mice were kept under controlled light (light, 07:00–19:00; dark, 19:00–07:00), temperature (22 ± 2 °C) and humidity (55 ± 10%) conditions. They had free access to water (25 ppm chlorine) and were fed *ad libitum* on a commercial diet (SDS Rat and Mouse No. 3 Breeding diet, RM3). All procedures and animal studies were carried out in accordance with the Animals (Scientific Procedures) Act 1986, UK, Amendment Regulations 2012 (SI 4 2012/3039). From crossings of wild-type (WT, C57BL/6J) mothers and heterozygous fathers with maternal deletion (m–/p+) of *Ndn*, *Magel2*, *Snord116,* double deletion of *Ndn* and *Magel2* (*Ndn*-*Magel2*^m−/p+^, DEL-NDN-MAGEL2-B6)^9^, and the large deletion (Large^m−/p+^), mixed litters of WT and heterozygous animals with paternal deletion (m+/p−) were produced and underwent different experiments in one of three experimental pipelines.

### Phenotyping

The experiments of the three pipelines (Fig. 2a) are summarized in Table 1 with the numbers and the age of the included animals. The first pipeline is the so called preweaning and nociception pipeline (pipeline I). It includes daily neonate observation and weight measures from 0 to 3 weeks of age and various experiments until the age of 12 weeks. The second pipeline consists of several experiments related to the metabolism (pipeline *II*). The third pipeline is dedicated to experiments mainly measuring motor and behavioral functions (pipeline III). Pipelines *II* and *III* have been performed on adult animals from 3 to 9 months of age.

#### Gene expression from droplet digital polymerase chain reaction (ddPCR)

Brains were extracted from selected mouse lines at either E18.5 or P4 and stored at −80 °C in RNALater (ThermoFisher) until genotyping via yolk sac or tail clip was confirmed. RNA from the selected animals was extracted using RNeasy Mini kit (Qiagen). Quantity and quality scores of extracted RNA were then checked using a nanodrop spectrophotometer (ThermoFisher) for the initial batches and latterly using a Qubit 4 (Invitrogen). 0.5 µg RNA was converted into cDNA using a High-capacity RNA-to-cDNA conversion kit (Applied Biosystems) following the manufacturer’s protocol. For each line, 1–2 heterozygous and 1–2 WT individual were selected and along with 2 male and female unrelated WT controls (cDNA) were assessed for the expression of 6 different target genes of interest in the PW region: *Mkrn3*, *Magel2*, *Ndn*, *Snrpn*, *Snord116* and *Ube3a*. (Note: *Snord116* expression in the *Ndn*^+/−^ line was not determined.) Each of these genes was run in technical duplicates with 2 reference genes (*Hprt*, and either *Gusb* or *Tbp*) using ddPCR (Bio-Rad)^47^. Raw ddPCR data files for gene expression analysis were run through a custom-made R script for automated data processing and data analysis performed using GraphPad Prism (10.3.1). The data is presented in the raw copies/µl format typical for absolute quantification using ddPCR, where the concentration of starting cDNA is 5 ng/RXN. In the subsequent analysis, we normalized gene expression levels such that for each experiment 1) the expression levels of the reference genes are normalized to match non-WT with WT levels, and 2) they are normalized to WT levels and expressed as percentages to be compared between genes (Fig. 1d; Supplementary Fig. 1; Supplementary Fig. 2).

#### Body weights

We measured body weights regularly during the life of the animals from all three experimental phenotyping pipelines: daily during the neonate observation, and weekly to bi-weekly during adulthood. The mice were weighed using a dynamic weighing balance (Ohaus) and were always weighed in the morning to limit the effect of the circadian rhythm on body weight. To reduce systematic uncertainties from measurement variability and individual day-to-day differences, we applied for each animal a linear regression of the body weight *w* in dependency on the age *a* (in days) to the model

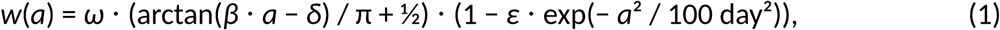

with four free parameters *ω* (the overall scale), *β*, *δ* and *ε*. This model performed well both for adults at the age starting from 2 weeks and pups at the age up to 3 weeks (Fig. 1f,g; Extended Data Fig. 2).

#### Neonate observation

We observed the pups from pipeline *I* daily during the first three weeks of their life^48^. Additionally to the daily weight measurement and general health observation, the milk spot^39^ was checked daily on days 0 to 4, the righting reflex was measured on days 2, 4, 6 and 8, the opening of left and right eyes was checked daily on days 13 to 16, and gait issues were examined daily on days 17 to 20. We encoded the milk spot ranging from 0 for no milk spot to 1 for visible milk spot. Similarly, the opening of left and right eyes is 0 for closed and 1 for opened eyes with values between 0 and 1 for partially opened eyes, and gait issues are 0 for no issues and 1 for observed issues, while 0.5 was assigned to animals with minor issues. The righting reflex was measured as the time needed for pups to turn themselves from their backs to the prone position (Supplementary Fig. 8).

#### Open field

We examined mice from pipeline *I* in the open field arena at the age of 4 to 5 weeks. Mice were placed in well-lit arenas (150–200 lux, 44 x 44 cm, equipment manufacturer: Noldus) for 20 minutes. The activity of the mice in the central zone (40% of the total area), periphery (area 8 cm towards the center from the wall) and whole arena was captured in 5 minute bins. The observed parameters include distance traveled, in-zone duration, latency and frequency of entering the zones (Ethovision XT v8.5; Supplementary Fig. 9).

#### Gait

We measured the gait of mice from pipeline *I* at the age of 4 to 6 weeks, and of mice from pipeline *III* at the age of 17 to 19 weeks. After acclimatization to the dark experimental room for at least 30 minutes, we put the animals at the start of a 10 cm wide and 1 m long corridor and let them walk three times or for up to 1 minute along the length of the corridor. Their illuminated footsteps were recorded at 81 fps with a Basler acA1300-75gc camera and StreemPix 7 software. The videos were analyzed with a custom trained DeepLabCut model, which resulted in variables such as walking speed, step size, stride length and left/right differences (Supplementary Fig. 10).

#### Microchipping

RFID microchips were injected subcutaneously into the lower left or right quadrant of the abdomen of mice from pipelines *I* and *III* at the age of 6 weeks^49^. These microchips were contained in standard ISO biocompatible glass capsule (11.5 mm x 2 mm, PeddyMark Ltd. UK). The procedure was performed on sedated mice (Isoflo, Abbott, UK) after topical application of local anaesthetic cream on the injection site prior to the procedure (EMLA Cream 5%, AstraZeneca, UK). The entry wound was sealed using a topical tissue adhesive, GLUture (Zoetis).

#### Home cage analysis (HCA)

The HCA was performed during 50 hours including three night and two day periods on the mice from pipeline *I* at the age of 6 to 7 weeks and twice on the mice of the behavioral pipeline at the age of 8 to 9 weeks and at the age of 20 to 21 weeks^49,50^. Mouse home cages were placed individually into one of the Home Cage Analyzer rigs (HCA; Actual Analytics, Edinburgh) on day 0 of the study time point and left undisturbed for 72 hours. The HCA rigs comprising of high-definition infra-red camera and a baseplate with RFID antennae on known locations, occupied two spaces on a standard IVC rack, where one half of the rig housed the camera and the electronics, while the other half housed the mouse home cage with the RFID baseplate. In bins of 6 minutes, observed parameters include the distance traveled, the time tracked in one of the 18 fields of the 3 × 6 fields area, and the time climbing. From the single-field time data, we calculated the times spent in the front (first 2 rows), middle (middle two rows) and back (last two rows) of the cage, as well as in the center (the 4 fields not at the border) and in the very center (the 2 most-centered fields). Further, we analyzed these parameters during night and day periods (Extended Data Fig. 5).

#### Hot plate

Mice from pipeline *I* underwent the hot plate experiment at the age of 10 to 11 weeks. Mice were put on the hot plate at 52 °C (BIO-HP) and as soon as they react on the heat, they were immediately removed or after 30 seconds if there was no reaction. The test was performed by two people watching the mice and mirrors. The observed parameters are reaction type and time (Supplementary Fig. 11a).

#### Von Frey

We performed the Von Frey nociception test on mice from pipeline *I* at the age of 11 to 12 weeks. Mice were placed on a mesh stand with square perforations measuring 5 × 5 mm in opaque acrylic insets of 8 cm × 6 cm × 10 cm and acclimatized for 60 minutes before testing. The range of von Frey filaments (Stoelting; Touch-Test Sensory Evaluator Kit of 12, Product number: 58011) used was 0.04 to 4.0 target force in grams (equivalent to filament number 3 to 11 and handle value 2.44 to 4.56), starting the testing always with filament number 7 with target force of 0.6 g (handle value 3.84). When all 4 feet were touching the platform, the von Frey filament was pressed against the plantar surface of the hind paw with enough force to cause the filament to bend, then held in place for up to 3 seconds. A positive response was recorded if the animal withdrew from the stimulus, including sharp withdrawal, licking, flicking, or flinching responses. After a positive response, the subsequent trial was conducted with the next smaller size of filament in the series. In the absence of a paw withdrawal response, the subsequent trial was conducted with the next larger size of filament in the series. Five consecutive trials, with an intertrial interval of at least 2 minutes, were conducted to complete 1 run. In the absence of administration of a sensitizing substance, both hind paws were tested at each time point, left hind paw first, followed by the right hind paw in a separate run. The observed parameters are the sensitivity expressed in PWT gram on the left and the right foot on the first and second day, calculated using the SUDO method^51^ approach with parameters *x* = 0.24 and *B =* −2.03. The test was repeated on three consecutive days and the data of the second and third day was averaged to get the parameters of interest (Supplementary Fig. 11b,c).

#### Magnetic resonance imaging by nuclear magnetic resonance (MRI-NMR)

We performed MRI-NMR measurement on the animals from pipeline *II* repeatedly at the age of 8 to 10, 12 to 14, 16 to 18, and 20 to 22 weeks for body composition analysis (EchoMRI 2014 Body Composition Analyser EMR-136). The observed parameters are fat, lean, free water and total water mass. We used linear regressions of the measurements to have more robust estimates of the parameters at 9 weeks and 21 weeks of age (Extended Data Fig. 3a,b,d–f).

#### X-Ray

We performed full body X-Ray scans including Dual X-ray Absorptiometry (DEXA) on mice from pipeline *II* at the age of 25 to 27 weeks. X-ray images of the mice were collected using the Faxitron X-Ray machine MX-20 whilst the mice were anesthetized (isoflurane). A lateral view, a dorsal-ventral view and a skull image were all taken to enable a full qualitative assessment of the integrity of the skeleton. A 2 cm lead bar was positioned to provide a calibration scale for the measurement of the tibia length, achieved using the ImageJ program. Apart from number of ribs, and digits and binary parameters for skeleton abnormalities, the only quantitative observed parameter from the skeleton analysis is the length of the tibia (Fig. 1h,i). The DEXA scans provided body composition observables for fat mass, lean mass, bone mineral density, and bone mineral content (Extended Data Fig. 3c).

#### Intraperitoneal glucose tolerance test (IPGTT)

We examined the glucose tolerance of mice from pipeline *II* at the age of 15 to 17 weeks with the IPGTT using Abbott Alphatrek 2 with whole blood strip on restrained animals. The mice were fasted overnight for a period not exceeding 18 hours. Local anesthetic (EMLA cream) was applied to the tail and a sample of blood was taken to determine the fasted blood glucose concentration (Accu-Chek glucose meter, Abbott, UK). The mice were injected intraperitoneally with 20% glucose solution at a concentration of 2 g/kg of body weight. Further blood glucose measurements were taken at 15, 30, 60 and 120 minutes after the injection of the glucose. Using the glucose concentration levels, we calculated the area under the curve (AUC) and the time of maximum glucose concentration (Supplementary Fig. 12a–c). More details are available at IMPRESS: https://mousephenotype.org/impress/ProcedureInfo?procID=532.

#### Intraperitoneal insulin tolerance test (IPITT)

We tested the insulin tolerance of mice from pipeline *II* at the age of 18 to 20 weeks with the IPITT using a similar procedure as for IPGTT. Mice were fasted for 4 hours before the test. We measured the glucose concentration before the insulin injection (1.25 iu for females and 1.5 iu for males at 0 minutes) and at 15, 30, 60, 45, and 90 minutes after the injection. Using the glucose concentration levels, we calculated the negative area under the curve (AUC) and the time of minimum glucose concentration (Supplementary Fig. 12d–f).

#### Temperature challenge

Mice from pipeline *II* at the age of 22 to 24 weeks underwent the temperature challenge test, where they were put in a fridge for 5 hours and body and fridge temperatures were measured every 30 minutes (Body temperature: Physitemp Rectal Thermometer, Fridge: BIO COLD Model BIO170FRSS for *Ndn*^+/−^ and *Magel2*^+/−^ and WT littermates, and Thermo Fisher T5X1205GX for *Ndn*-*Magel2*^+/−^ and WT littermates; Supplementary Fig. 13).

#### Calorimetry

The metabolic rate of mice from pipeline *II* at the age of 25 to 27 weeks was assessed using indirect calorimetry. Mice were individually housed overnight for a period of 21 hours in Phenomaster cages (TSE Systems, Germany) with standard bedding and igloos. The air content in each cage was sampled for 1 minute, in turn, and a reference cage with no mice in it was also sampled for comparison. The oxygen consumption and carbon dioxide production for each mouse was then determined as the difference between the reference amount of gas and the level in the mouse cage. Observed parameters include total and ambulatory activity, heat production, carbon dioxide production, and oxygen consumption. Additionally, the respiratory exchange ratio is defined as the ratio of the latter two (Supplementary Fig. 14).

#### Auditory brain stem response (ABR)

We tested the auditory brain stem response on some mice from pipeline *II* at the age of 26 to 28 weeks^52^. Mice were anesthetized with subcutaneous injections of ketamine and xylazine and placed on a warm plate. Three electroencephalogram (EEG) electrodes were placed on the mouse head and a speaker was placed directed towards one ear. Different stimuli of click and various frequency tones (6, 12, 18, 24, 30 kHz) at different volumes from 0 to 85 db (Medusa4Z Preamp, TDT hardware hub) were used to measure the EEG response. The observables are the hearing response onset volumes for each stimulus type that were annotated by hand (Supplementary Fig. 22).

#### Hematology

The blood of the *Ndn*-*Magel2*^+/−^ mice and their WT littermates from pipeline *II* was collected by a small incision on the tail after applying local anesthetic EMLA cream, and analyzed using Siemens Medical Solutions Diagnostics Hematology Analyzer Advia 2120i (Supplementary Fig. 4).

#### Enzyme-linked immunosorbent assay (ELISA)

We performed ELISA to measure protein contents of blood collected at the end-of-life of *Ndn*^+/−^ and *Ndn*-*Magel2*^+/−^ mice and their WT littermates from pipeline *II*, according to the manufacturer’s instructions using the following kits: Neurophysin-2 RayBiotech (cat no. EIA-NEU2) for *Neu2*, and Oxytocin RayBiotech (cat no. EIA-OXT) for *Oxt* in dilutions of 1 in 4, and IGF-1 Bio-Techne (cat no. MG100) for *Igf1* in dilutions of 1 in 500 (Supplementary Fig. 3).

#### Grip strength

We measured the grip strength of the mice from pipeline *III* at the age of 9 to 10 weeks, and again at the age of 21 to 22 weeks using the Bioseb HMGU plate Bio-GT3+MR. The observed variables are three trials for the front limb and three trials for the combined front and hind limb strength. In the further analysis, we calculated mean values and normalized the measured strength values by the body weight (Supplementary Fig. 16).

#### Combined SHIRPA and dysmorphology (SHIRPA)

SHIRPA was performed on the mice from pipeline *III* at the age of 9 to 10 weeks, and again at the age of 21 to 22 weeks^53^. The combined SHIRPA and dysmorphology test identifies physical and behavioral abnormalities through observation. Mice were individually observed in a series of environments to test a range of attributes including hearing, visual placement, activity, motor coordination, righting ability as well as morphological features. Locomotor activity was assessed in an arena of 121 cm² (Supplementary Fig. 17).

#### Three chamber

At the age of 13 to 16 weeks, the three chamber test was performed on the mice of the behavioral pipeline in the Noldus 3-chamber maze 1 (25 cm height, 39.1 cm width, 58.5 cm length). At the beginning of the habituation phase (10 minutes), the mouse was put in the central chamber and allowed to freely explore the three chambers, which were connected via open gates. At the start of the testing phase (10 minutes) a stranger mouse was placed in one lateral chamber and an object in the other lateral chamber in weighted wire cages, and the mouse was again put into the central chamber. The video tracking of the mice was performed using Noldus Ethovision XT (versions 8.5 and 15). Observable quantities include total distance traveled, the duration of stay, the latency of first entry, and the frequency of entries for each of the chambers, as well as for interactions with the mouse and the object (Supplementary Fig. 18).

#### Spontaneous alternation

We measured spontaneous alternation of the mice from pipeline *III* at the age of 16 to 17 weeks. The Opaque Grey Perspex® Y-maze with arms at 120° (arm dimensions: 20 cm height, 8 cm width, and 30 cm length) was made by the estates and engineering department of MLC Harwell. The mice were put at the end of one arm of a y-maze and their subsequent entries to the different arms were recorded. The observed quantities are the total number of alternations and the number of spontaneous alternations (Supplementary Fig. 19).

#### Home-cage motor function (HCMF)

The mice from pipeline *III* were introduced to the HCMF experiment at the age of 22 to 26 weeks. Mice were single-housed and learned wheel running during three weeks. After the second week the wheel (11.5 cm diameter) was replaced by a complex one with irregularly missing steps. The raw data (TSE PhenoMaster pre-version, 2009-8998_223652, and version 7.6.7, 2021_6110) includes hourly values for distance ran, total time running and maximum speed. We combined these measurements into variables for day and night and for each week of experiment. Ratios of 2nd to 1st and 3rd to 2nd week indicate learning behavior (Supplementary Fig. 20).

#### Passive infrared (PIR)

We examined the female and male mice from pipeline *III* in the PIR cages at the age of 26 to 28 weeks and at 29 to 31 weeks, respectively. They were singly housed and observed during 2 weeks for circadian rhythm and sleep measurement using the COMPASS passive infrared system^54^. During 4 to 5 days, they were kept in the usual light-dark (LD, light periods 7:00 to 19:00) condition, followed by dark-dark (DD) condition during 8 to 10 days. The switch from LD to DD condition was applied at 7:00 after end of LD and thus, LD usually consists of 4 light day and 5 dark night periods. To investigate changes during the DD condition we split it further into DD1, which is the 1st to 4th day after the last day of LD (4 dark night and 4 dark day periods), and DD2, which consists of the fifth to last day of DD. We applied an automatic on-/offset detection algorithm on the activity traces of each animal, followed by manual validation (Extended Data Fig. 4).

#### Stress induced hyperthermia

Stress induced hyperthermia was measured on the mice from pipeline *III* at the age of 30 weeks or older. Mice were singly housed for at least 24 hours before the test. On the day of test, mice were acclimatized to the testing room (19–23 °C). First, rectal temperature was measured using Physitemp Rectal Thermometer, the mice were returned to the singly housed home cage, 10 minutes later, a second reading was taken. The difference between these two readings was taken as an indicator of stress induced hyperthermia (Supplementary Fig. 21).

#### Acoustic startle

We measured the reaction movement amplitudes on acoustic pulses^55^ of the mice from pipeline *III* at the age of 30 weeks or older in the Med Associates Acoustic Startle Cubicle ENV 022S with sound generator Med Associates PHM 255A, analyzed with Med Associates Startle Reflex (version 6.0). Mice were placed in an acoustic startle chamber (Med Associates Inc, USA) and acclimatized to a background noise level of 53 dB for 5 min, followed by exposure to five 120 dB startle tones. Each animals was tested on 90 trials (10 trials for each amplitude) with random inter-trial intervals of 20 to 30 seconds. Responses to the startle tone were measured for 100 ms following the start of the startle tone using a piezoelectric transducer in the floor of the chamber which detected movement of the animal. Eight amplitudes were measured on different sounds: background noise at 53 db, pre-pulses at 56 db, 58 db and 65 db, startling pulse at 120 db, and pre-pulse-startling-pulse events with the three different pre-pulse volumes (80 ms interval between pre-pulse and startling pulse; Supplementary Fig. 22).

### Statistical analysis

#### Variable transformation

For each line the outcome of a phenotypic variable is transformed using the Yeo-Johnson power transformation^56^ fitted on the outcomes of WT animals of that line. Thus, the transformed outcome of WT animals has mean of 0 and variance of 1.

#### Single-experiment principal component analysis (ePCA)

For each single experiment the power transformed variables were used as input to principal component analysis (PCA). The principal components of these ePCAs are selected as “outcomes” in the further analysis, if their explained variance is greater than 1%. The ePCAs and the following analyses were all performed separately for female and male mice (Supplementary Fig. 23 and Supplementary Fig. 24).

#### ACE model: partitioning genetic and environmental variance

We applied the ACE model to phenotyping outcomes (from ePCAs), describing each outcome as the sum of additive genetic effects (*A*), common environmental effects (*C*), and individual-specific residuals (*E*). For an individual *i* with genotype *g* ∊ {WT, *Ndn*^+/−^, *Magel2*^+/−^, *Ndn-Magel2*^+/−^}, the outcome *y* is expressed as:

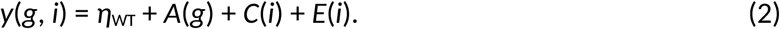

The additive genetic effect *A*(*g*) is zero for WT animals, *A*(WT) = 0, and a line- and sex-dependent value for heterozygous animals. The impact of single paternal gene deletions on the phenotypic outcome are *A*(*Ndn*^+/−^) = Δ*_Ndn_*^+/−^ and *A*(*Magel2*^+/−^) = Δ*_Magel2_*^+/−^. The additive genetic effect of the *Ndn-Magel2* deletion is written as a sum of single deletions and an additional term that accounts for possible non-additive contributions, *A*(*Ndn*-*Magel2*^+/−^) = *ΔNdn*^+/−^ + *Δ_Magel2_*^+/−^ + *δ_Ndn-Magel2_*^+/−^. The common environmental errors describe fixed effects from quantitative covariates, such as litter size and age at experiment, and from qualitative covariates, such as the batch or line, the experimenter or the month when the experiment was performed. They can be written as

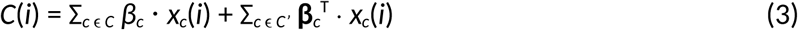

with the fixed effects (parameters) *β_c_* and **β***_c_*, and the values *x_c_*(*i*) of the covariates *c* of the sets of quantitative (discrete or continuous) covariates *C* and qualitative (nominal or ordinal) covariates *C’*. For qualitative covariates, one-hot encoding is used, and thus the fixed effects *β_c_* and the covariate values *x_c_*(*i*) are vectors with the length of the number of classes or categories minus one. Extended Data Table 1 summarizes the covariates that we considered in the analysis.

The term *E*(*i*) corresponds to the random effect of uncorrelated environmental noise. The constant parameter *η*_WT_ is the estimated value of the phenotypic outcome for WT animals. Thus, group estimates of phenotypic outcomes can be calculated as

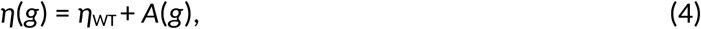

removing the effects of common environment and unique environmental noise. The total variance of the phenotypic variable *σ*_y_ ^2^ is the sum of variances of the additive effects as

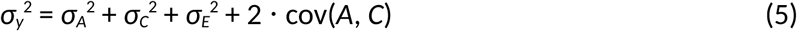

with the covariance term cov(*A*, *C*). The covariances with the unique environmental errors *E* are by construction equal to zero. Relative variances, called “effect sizes”, are defined as

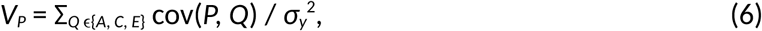

e.g. the effect size of the additive genetic effect is

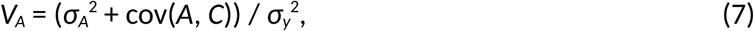

such that their sum is by construction equal to 1,

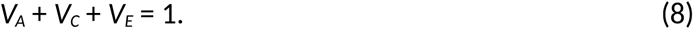

We calculated the covariances and effect sizes for every outcome of the single experiments (Supplementary Fig. 5–7). The AE estimates are defined as estimates of the outcomes removing the common environmental effect with

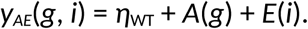

#### Single-pipeline principal component analysis (pPCA)

Because each animal belongs to one of the three experimental pipelines, the final PCAs cannot be done using all animals together and, we performed single-pipeline PCAs (pPCAs) instead. As input to the pPCAs we used the AE estimates of the outcomes, *y_AE_*, removing the common environmental effects. This was necessary because of the big effect of common environment, in particular the batch effect could be removed by this choice. This is well observable comparing the distributions of PCs of the pPCA for WT littermates using the original outcomes *y* (Supplementary Fig. 25a) with the ones using the AE estimates *y_AE_* (Supplementary Fig. 25b). We excluded the body weight measures from the pPCAs. Because also in a single pipeline not all experiments were performed on all animals, we applied a selection process. Firstly, only the outcomes of experiments with testing on more than 50% of the mice of a pipeline were considered in the pPCAs. Thus, ABR, hematology and ELISA were excluded from the pPCA of pipeline *III*. Subsequently, only animals that were tested in 50% of the experiments of a pipeline were kept for the pPCA. After the selection process, still not all experiments were performed on all remaining animals, and we used iterative imputation to fill the empty values^57^.

#### Simulation of digital twins

We performed simulation of digital twins by random choice of experimental outcomes and covariates using the probability density functions and the estimates of the parameters in the ACE model. Finally, simulated animals did not have the restriction of not being tested in some of the experiments and thus, we performed only one final PCA combining all experiments from the pPCAs. For comparison, pPCAs on simulations have been performed, too, varying the number of included animals per group (Supplementary Fig. 26).

## Data availability

The generated mouse models are available at MRC Harwell through request at FPWR. The raw experimental phenotyping data and the processed data tables will be made available at G-Node at https://gin.g-node.org/pcan/Wolff_et_al_2025_PCAN_raw_data (DOI: 10.12751/g-node.xxxxxx) and https://gin.g-node.org/pcan/Wolff_et_al_2025_PCAN_processed_data (DOI: 10.12751/g-node.yyyyyy), respectively.

## Code availability

The analysis source code will be made available at Codeberg at https://codeberg.org/pcan/pcan (DOI: https://doi.org/10.12751/g-node.zzzzzz).

## Acknowledgments

This project was funded by the Foundation for Prader-Willi Research (FPWR). We thank Prof. Steve D. M. Brown for critical reading of the manuscript.

## Author contributions

RW - Formal analysis, Methodology, Software, Visualization, Writing - Original draft preparation; TVS - Conceptualization, Funding Acquisition, Project Administration; LCB - Project administration; NK - Conceptualization, Project administration; JLR - Conceptualization, Methodology; MS - Data curation, Investigation, Methodology, Supervision; SEW - Conceptualization, Project administration, Supervision; LT - Methodology; AJA - Investigation; RSB - Data curation, Investigation, Methodology, Supervision; AH - Investigation; ARI - Conceptualization, Funding acquisition, Methodology, Supervision, Writing - Review & Editing; FM - Conceptualization, Funding acquisition, Methodology, Supervision, Writing - Review & Editing; VT - Conceptualization, Funding acquisition, Methodology, Project administration, Supervision, Writing - Review & Editing

## Extended Data Tables

**Extended Data Table 1.**
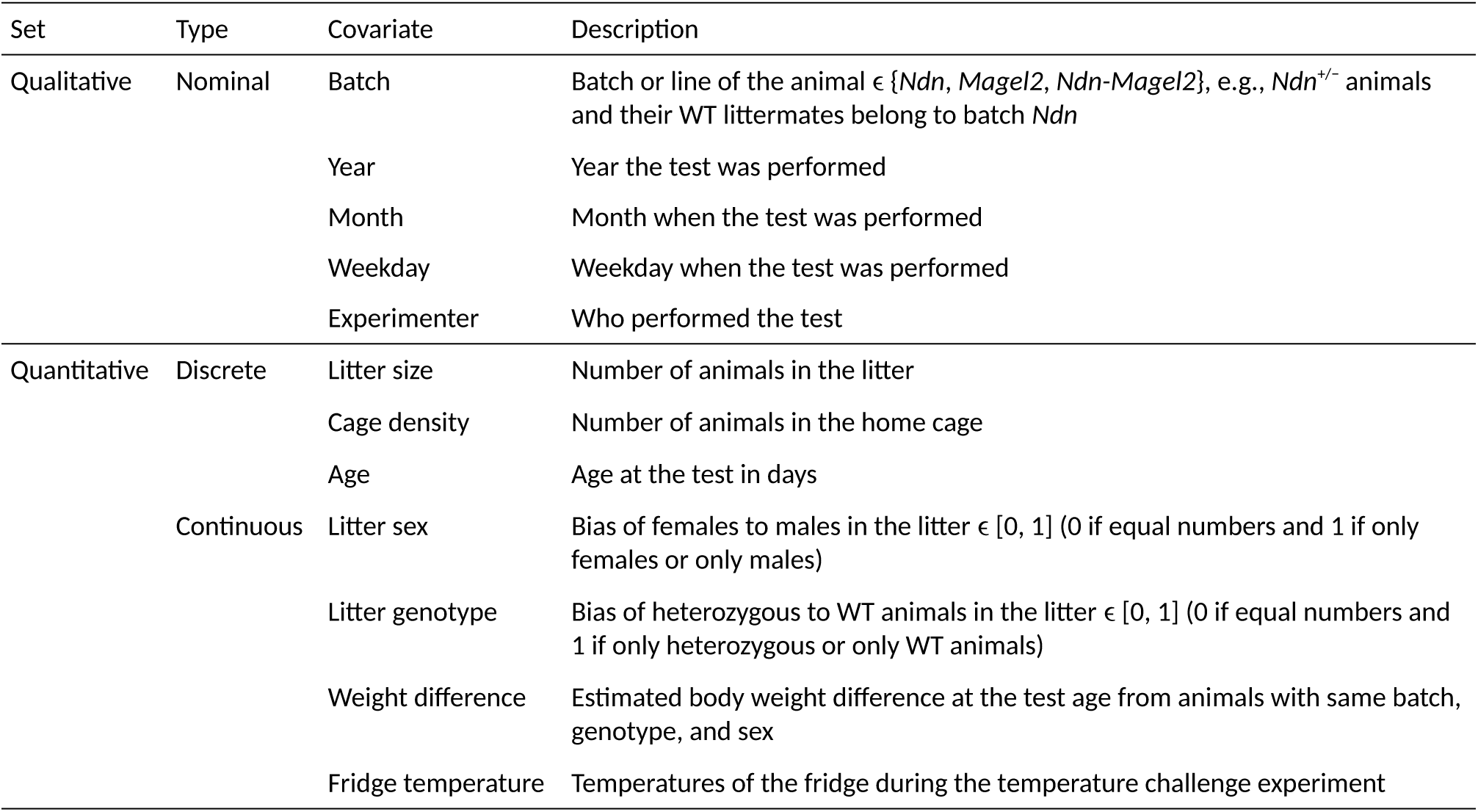
Covariates considered in the ACE model.

| Set | Type | Covariate | Description |
| --- | --- | --- | --- |
| Qualitative | Nominal | Batch | Batch or line of the animal $\in \{Ndn, Magel2, Ndn-Magel2\}$, e.g., *Ndn*<sup>+/-</sup> animals and their WT littermates belong to batch *Ndn* |
|  |  | Year | Year the test was performed |
|  |  | Month | Month when the test was performed |
|  |  | Weekday | Weekday when the test was performed |
|  |  | Experimenter | Who performed the test |
| Quantitative | Discrete | Litter size | Number of animals in the litter |
|  |  | Cage density | Number of animals in the home cage |
|  |  | Age | Age at the test in days |
| | Continuous | Litter sex | Bias of females to males in the litter $\in [0, 1]$ (0 if equal numbers and 1 if only females or only males) |
| | | Litter genotype | Bias of heterozygous to WT animals in the litter $\in [0, 1]$ (0 if equal numbers and 1 if only heterozygous or only WT animals) |
|  |  | Weight difference | Estimated body weight difference at the test age from animals with same batch, genotype, and sex |
|  |  | Fridge temperature | Temperatures of the fridge during the temperature challenge experiment |

## Extended Data Figures

**Extended Data Fig. 1.**
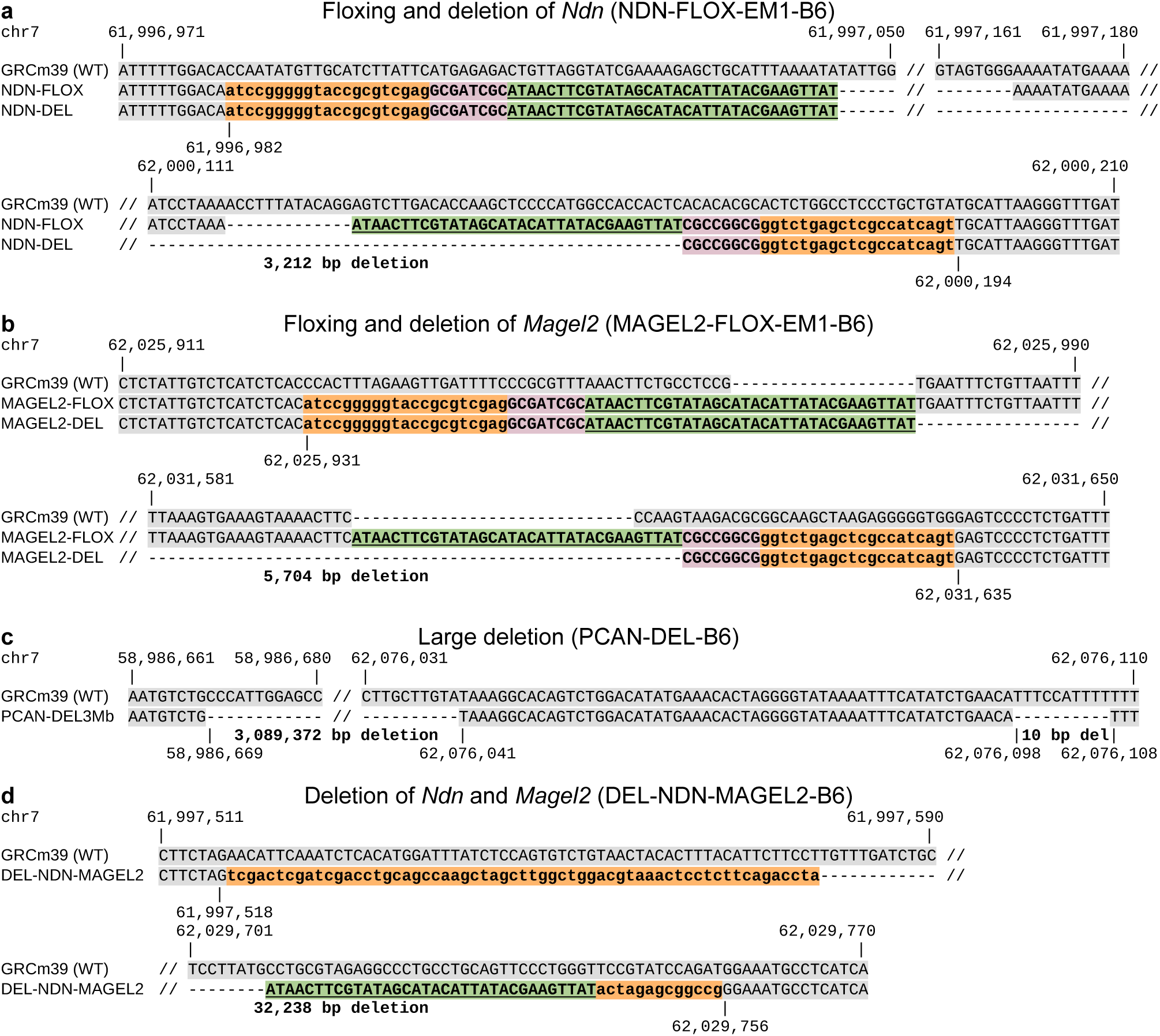
Sequences and size of deletions and floxed alleles. Wildtype (WT, C57BL/6J, GRCm39) sequences are highlighted with gray color and inserted sequences with orange, pink and green color, where the latter color highlights the loxP recognition sites (34 base pairs, underlined sequences). **a**, Floxed and deleted alleles for *Ndn*^+^*^/−^* with 3,212 deleted base pairs. **b**, Floxed and deleted alleles for *Magel2*^+^*^/−^* with 5,704 deleted base pairs. **c**, Large deletion for Large^+^*^/−^* with 3,089,372 plus 10 deleted base pairs. **d**, Double deletion of *Ndn* and *Magel2* for *Ndn*-*Magel2*^+^*^/−^* with 32,238 deleted base pairs (DOI: 10.1172/jci.insight.185159).

**Extended Data Fig. 2.**
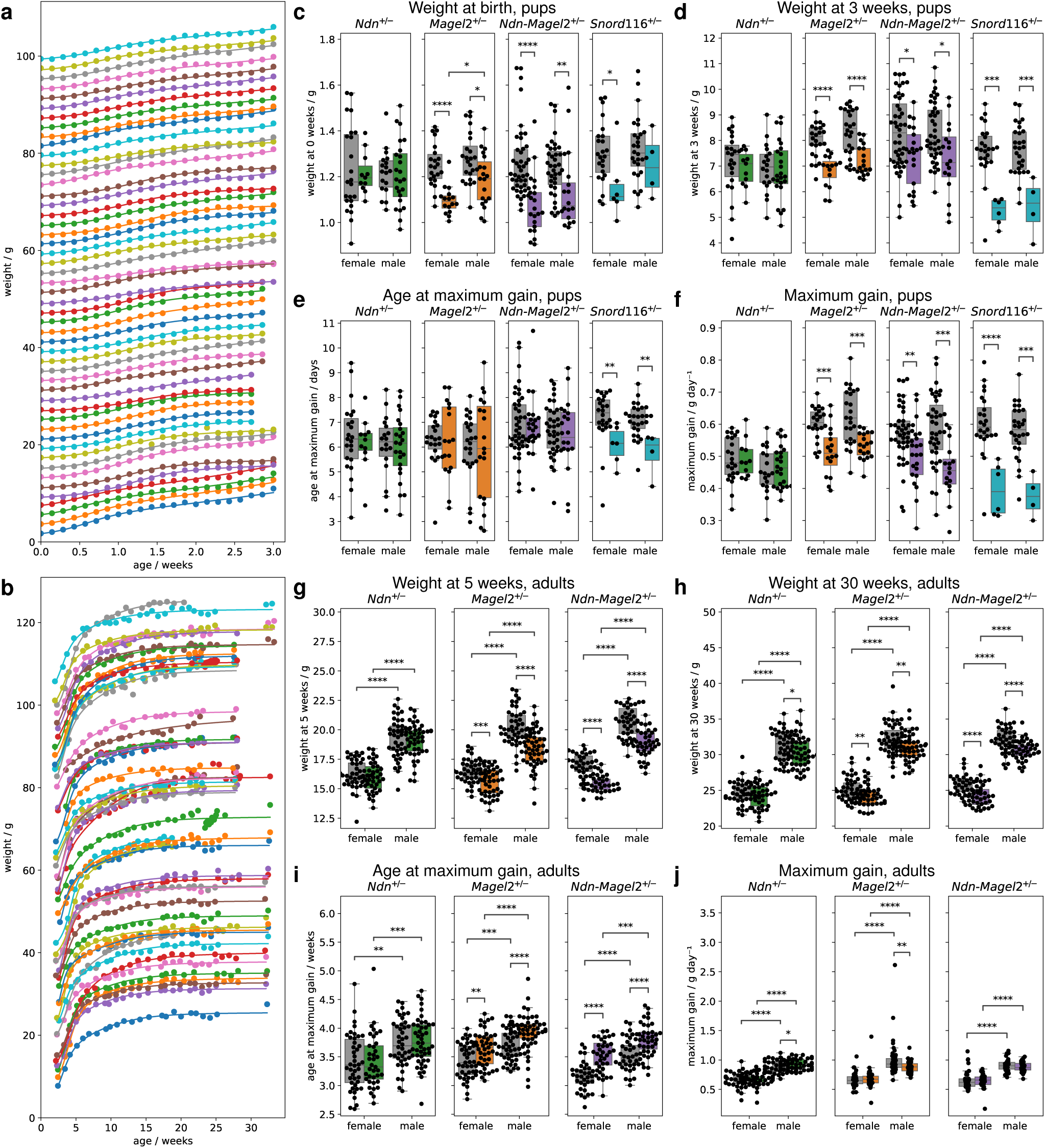
Body weights. **a**,**b**, Body weight curves for 50 example pups (**a**) and 50 example adult animals (**b**), stacked on top of each other with additional 2 g per animal (thus, only the bottom, blue animals are shown with weights corresponding to the value on the *y*-axis). The single points show the single measurements and the solid lines depict the fitted curves obtained by linear regression. **c**–**f**, Estimated parameters from linear regressions of pup body weight measurements from animals from pipeline *I* : weight at birth (**c**), weight at age of three weeks (**d**), age at maximum gain (**e**), and maximum gain (**f**). **g**–**j**, Estimated parameters from linear regressions of adult body weight measurements from animals from pipelines *I*, *II* and *III* : weight at age of 5 weeks (**g**), weight at age of 30 weeks (**h**), age at maximum gain (**i**), and maximum gain (**j**). Boxplot boxes in **c**–**f**,**g**–**j** extend from the first to the third quartile, with a line at the median and whiskers at the fartherst data points within the 1.5 *×* inter-quartile range. Reported *p*-values are calculated using the Mann-Whitney-Wilcoxon test. Right, colored boxes: heterozygous animals; left, grey boxes: their WT littermates; *: *p <* 0.05; **: *p <* 0.01; ***: *p <* 0.001; ****: *p <* 0.0001.

**Extended Data Fig. 3.**
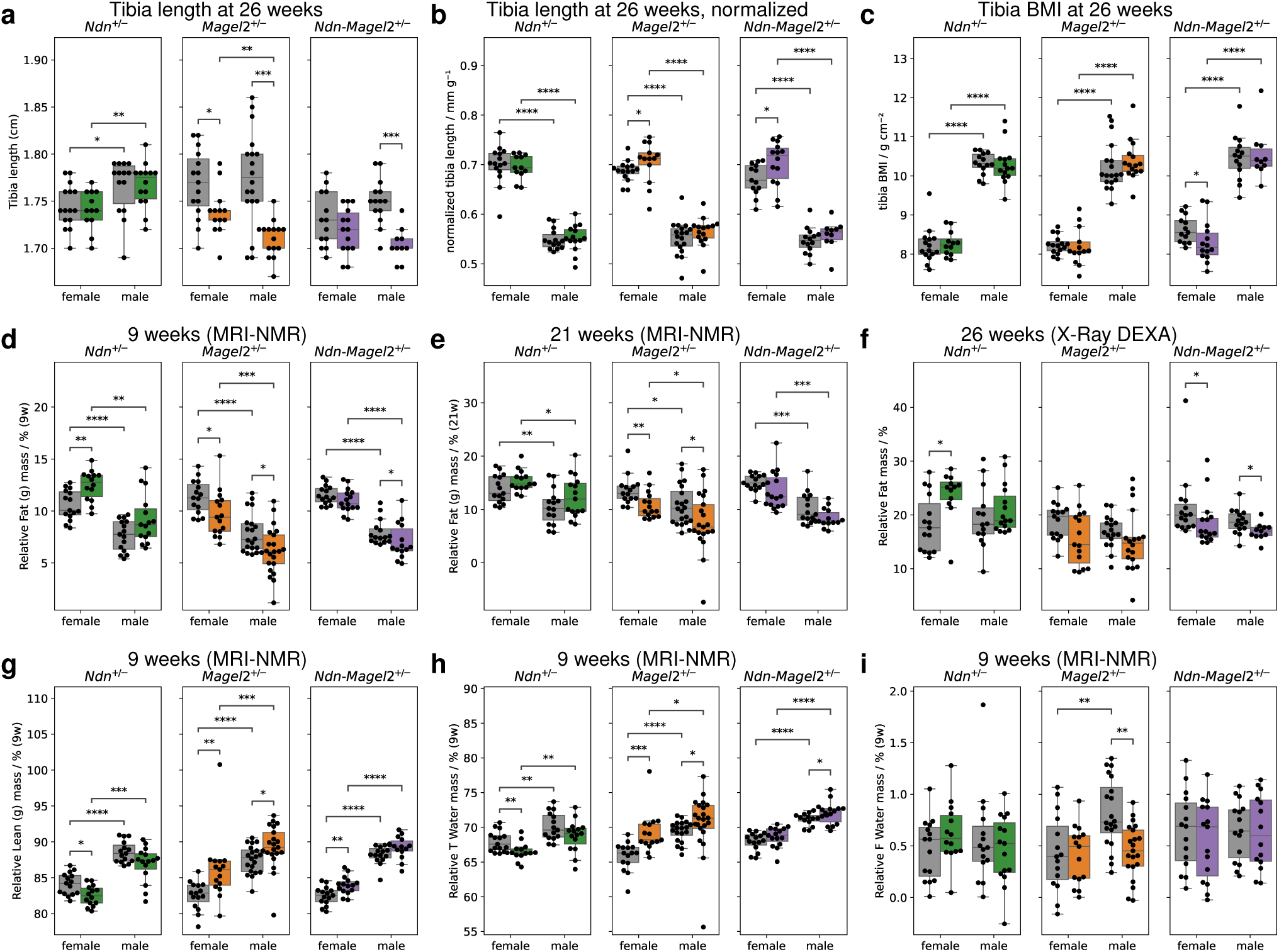
Body size and composition from MRI-NMR and X-Ray DEXA scans. The experiments were performed on animals from pipeline *II* at the age of around 9, 13, 17 and 21 (MRI-NMR), and 26 (X-Ray) weeks. **a**, Tibia length at mean age of 26 weeks from X-Ray scans. **b**, Tibia length, normalized to body weight, for the same animals as in **a**. **c**, A body-mass-index-(BMI)-like measure using the tibia length instead of the true body size for the same animals as in **a**. **d**–**f**, Relative body fat mass at age of 9 weeks (**d**) and at age of 21 weeks (**e**) from estimates using linear regression of multiple MRI-NMR experiments, and at mean age of 26 weeks (**f**) from X-Ray DEXA scans. **g**–**i**, Relative body lean mass (**g**), relative total water mass (**h**), and relative free water mass (**i**) at age of 9 weeks from estimates using linear regression of multiple MRI-NMR experiments. Boxplot boxes extend from the first to the third quartile, with a line at the median and whiskers at the fartherst data points within the 1.5 *×* inter-quartile range. Reported *p*-values are calculated using the Mann-Whitney-Wilcoxon test. Right, colored boxes: heterozygous animals; left, grey boxes: their WT littermates; *: *p <* 0.05; **: *p <* 0.01; ***: *p <* 0.001; ****: *p <* 0.0001.

**Extended Data Fig. 4.**
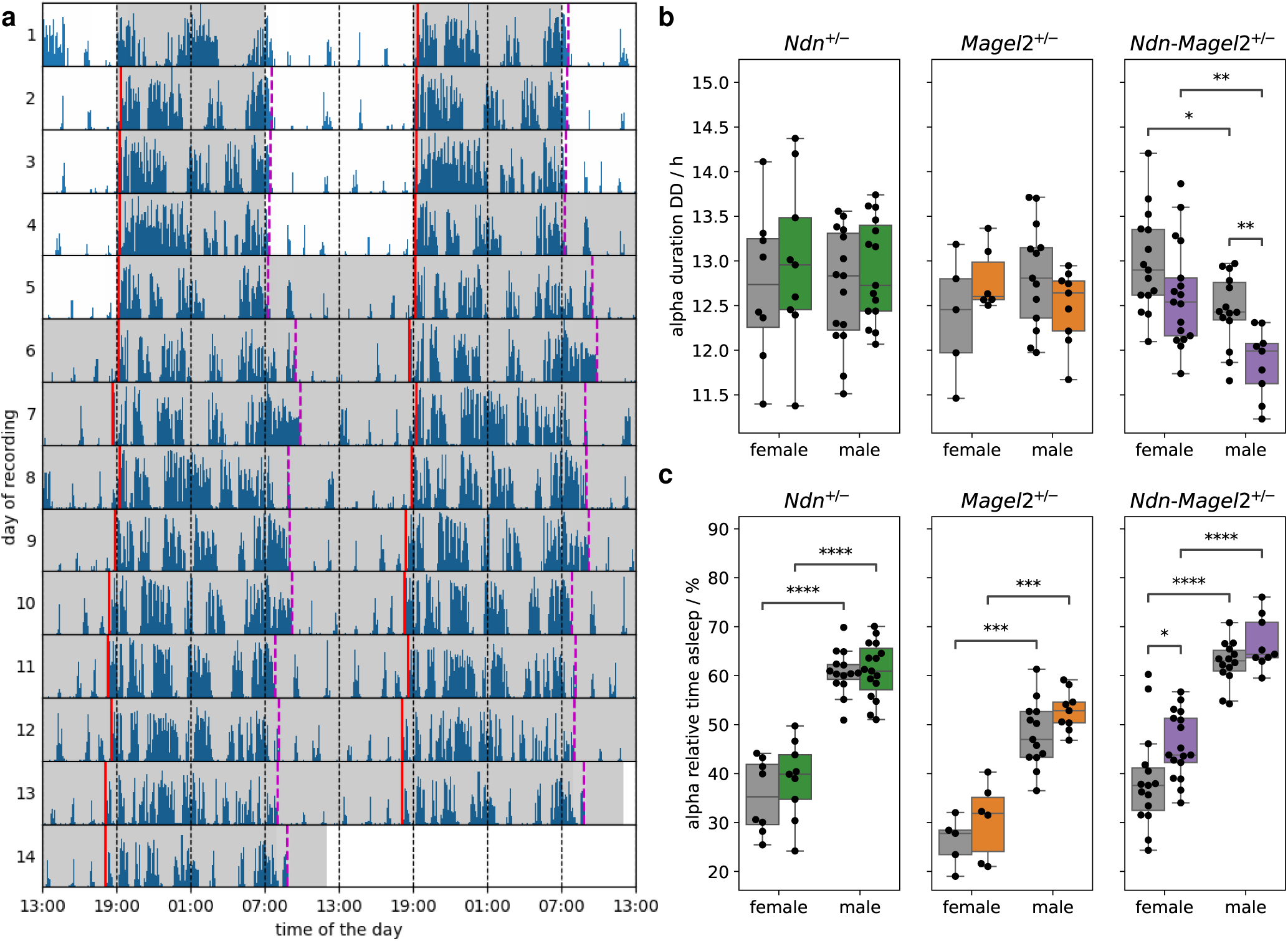
Sleep and circadian rhythm from passive infrared (PIR). The experiments were performed on animals from pipeline *III* at the age of around 28 weeks. **a**, Actogram for an example female WT mouse. The activity during each day is shown by the blue bars. Automatically estimated and manually validated activity onsets and offsets are indicated by vertical solid, red and dashed, purple lines, respectively. The shaded area indicates the dark phase, which was during 4 days from 19:00 to 07:00 during light-dark (LD) exposure. During dark-dark (DD) exposure the lights were kept off 24 hours. **b**, Duration of activity from estimated onsets to offsets during DD exposure. **c**, Relative time asleep during the alpha phase, which is the phase between estimated onsets and offsets for heterozygous animals and their WT littermates. Boxplot boxes in **b**,**c** extend from the first to the third quartile, with a line at the median and whiskers at the fartherst data points within the 1.5 *×* inter-quartile range. Reported *p*-values are calculated using the Mann-Whitney-Wilcoxon test. Right, colored boxes: heterozygous animals; left, grey boxes: their WT littermates; *: *p <* 0.05; **: *p <* 0.01; ***: *p <* 0.001; ****: *p <* 0.0001.

**Extended Data Fig. 5.**
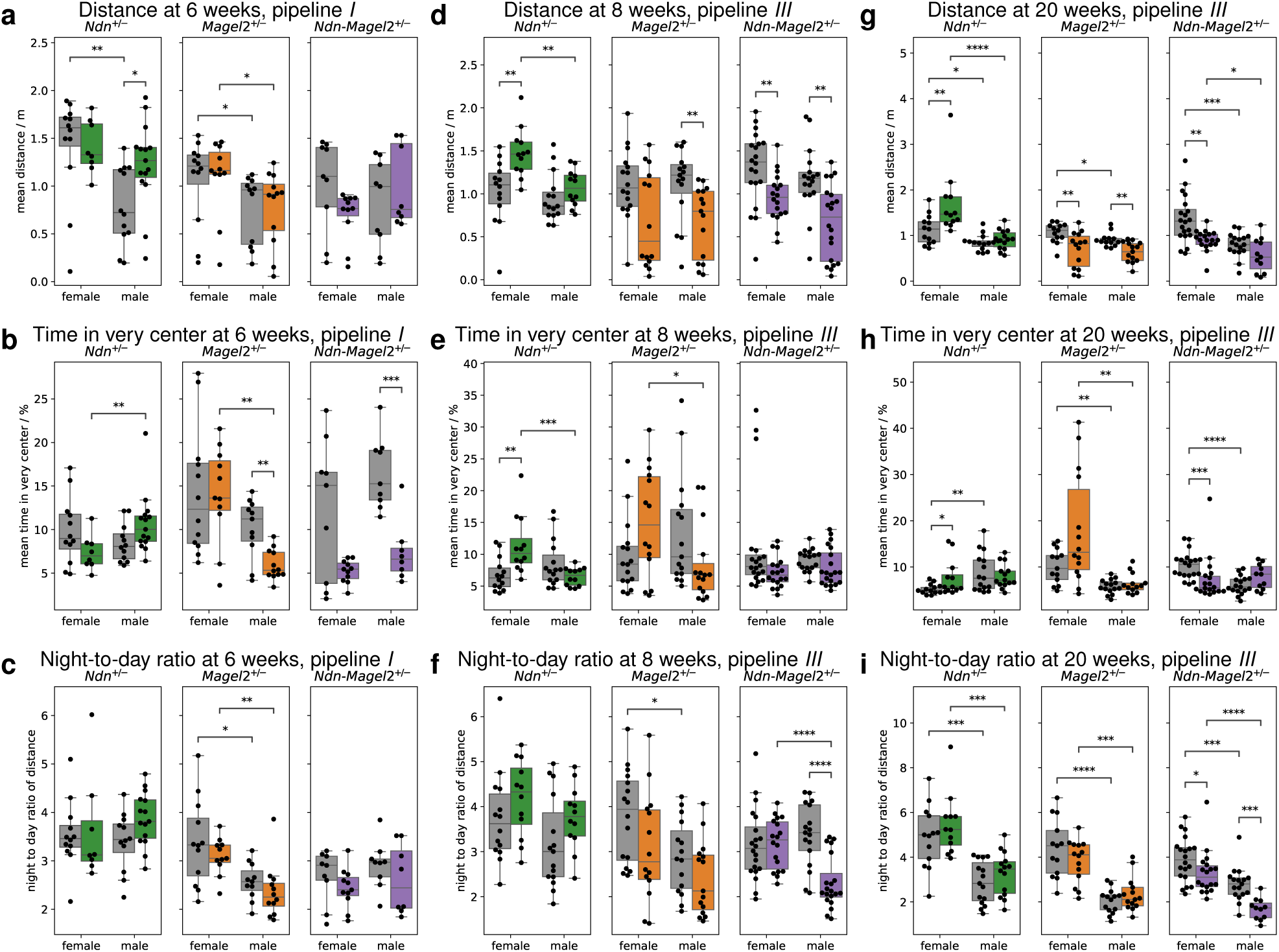
Home-cage analyzer (HCA) experiment. **a**–**c**, Estimated parameters for animals at the age of around 6 weeks from pipeline *I*. **d**–**i**, Estimated parameters for animals at the age of around 8 weeks (**d**–**i**) and 20 weeks (**g**–**i**) from pipeline *III*. **a**,**d**,**g**, Mean distance traveled. **b**,**e**,**h**, Time in the very center, which are the two central fields in the area of 3 *×* 6 fields. **c**,**f**,**i**, Ratio of the distance traveled at night to the distance traveled at day. Boxplot boxes extend from the first to the third quartile, with a line at the median and whiskers at the fartherst data points within the 1.5 *×* inter-quartile range. Reported *p*-values are calculated using the Mann-Whitney-Wilcoxon test. Right, colored boxes: heterozygous animals; left, grey boxes: their WT littermates; *: *p <* 0.05; **: *p <* 0.01; ***: *p <* 0.001; ****: *p <* 0.0001.

**Extended Data Fig. 6.**
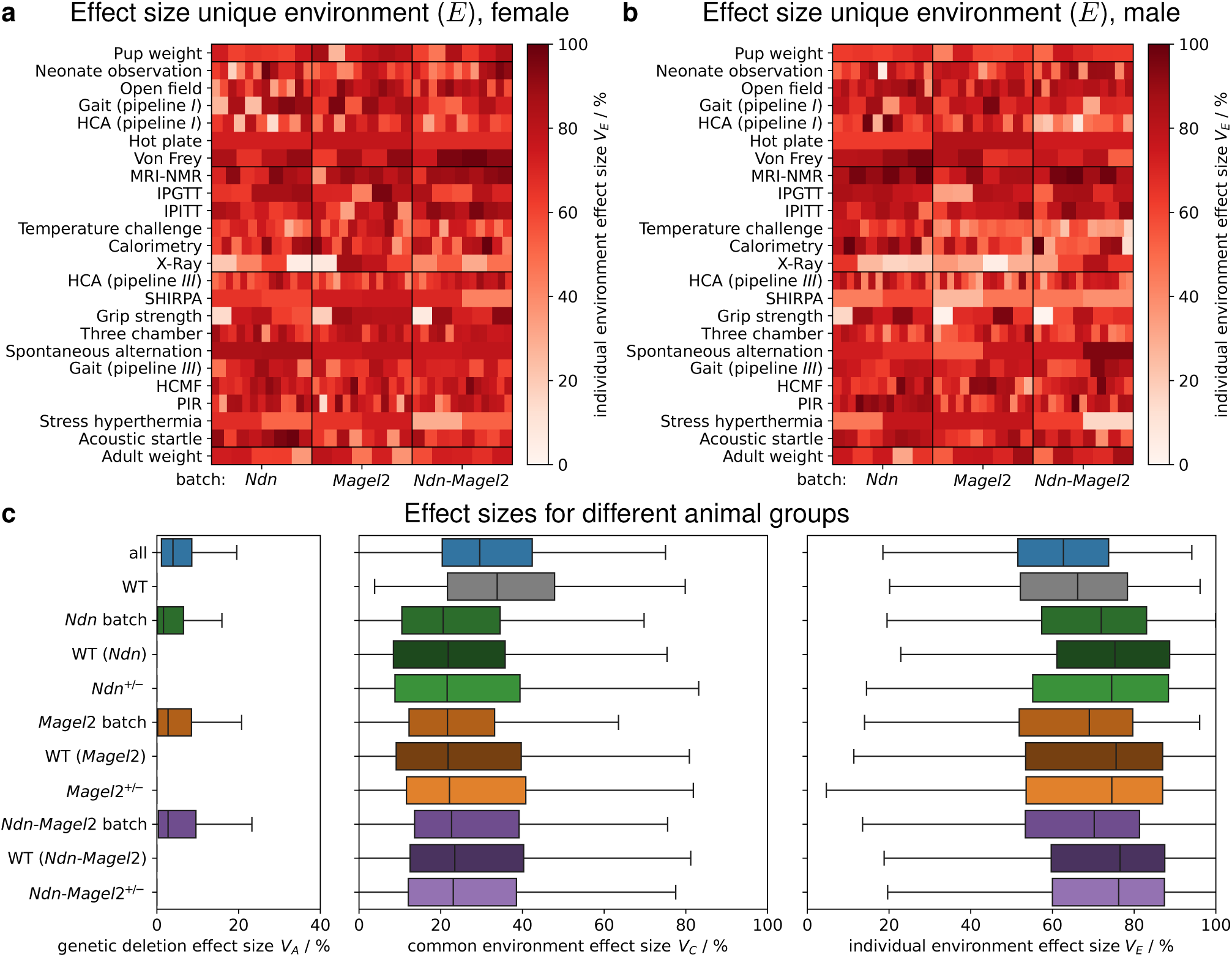
Effect sizes of unique environment and overall group comparison. **a**,**b**, Effect sizes by unique environment for females (**a**) and males (**a**). Each data point in **a**,**b** corresponds to one single-experiment principal component, sorted from left to right by highest explained variance. **c**, Effect sizes of genetic deletion (left) common environment (middle), and individual environment (right), comparing different animal groups, summarizing all PCs from all experiments, and female and male animals. Only mixed groups of WT and heterozygous animals have non-zero genetic deletion effect size. The groups of all animals and all WT animals have the highest common environment effect size because the other groups are from a single batch and thus, do not have the batch effect. Heterozygous animals from a single group have comparable common environment effect size to their WT littermates. Boxplot boxes extend from the first to the third quartile, with a line at the median and whiskers at the fartherst data points within the 1.5 *×* inter-quartile range.

---

**Supplementary Fig. 1.**
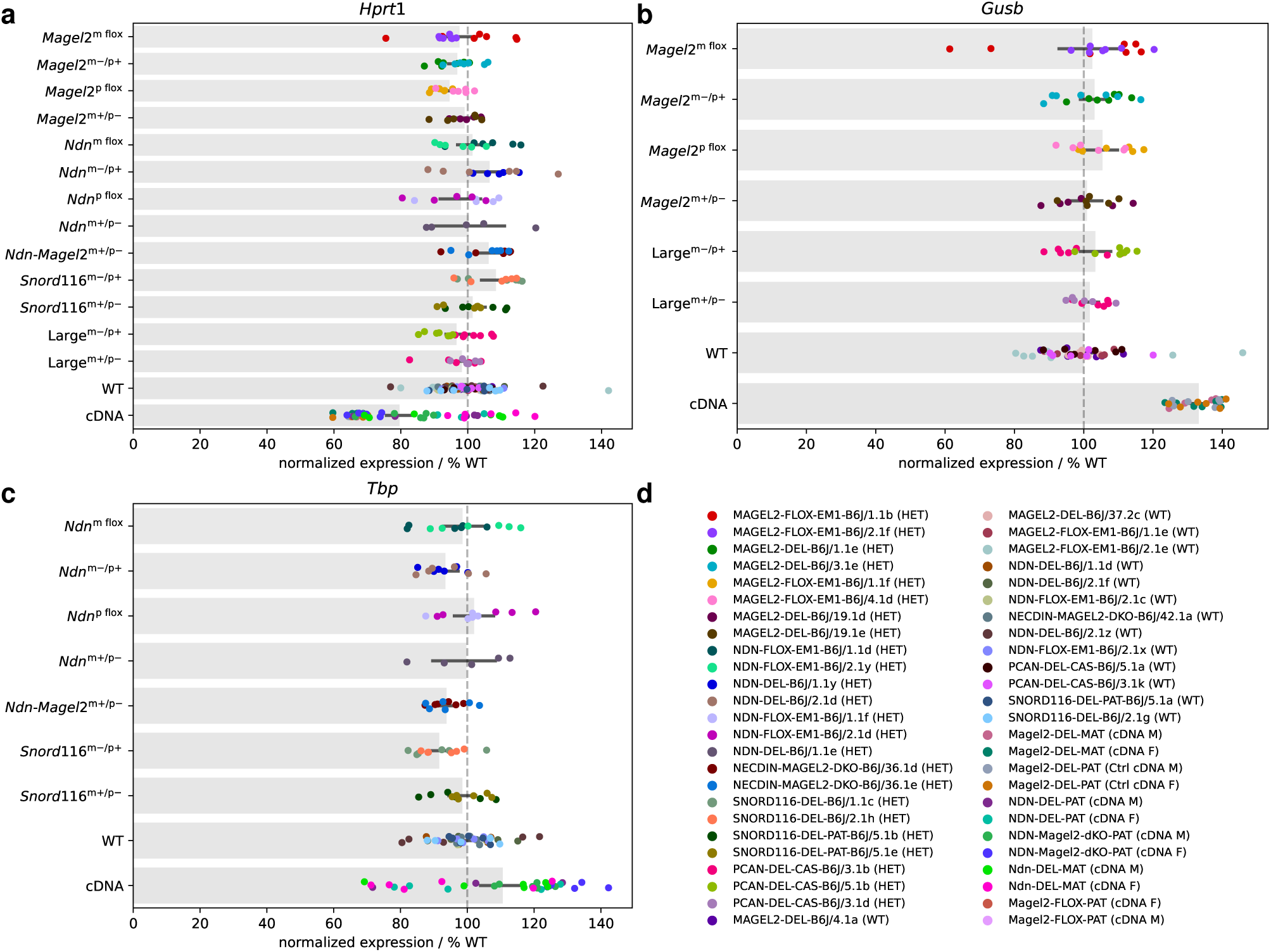
Expression levels of reference genes from ddPCR. **a**–**c**, Normalized gene expression levels for different mouse lines with maternal deletion (m*−/*p+), paternal deletion (p+/m*−*), floxed maternal allele (m flox), floxed paternal allele (p flox), and no deletion (WT), and cDNA controls, normalized to WT level (100%, dashed, grey line). Single points indicate technical replicates of with biological replicates shown with same color. **d**, Legend of biological replicates. HET: heterozygous animal.

**Supplementary Fig. 2.**
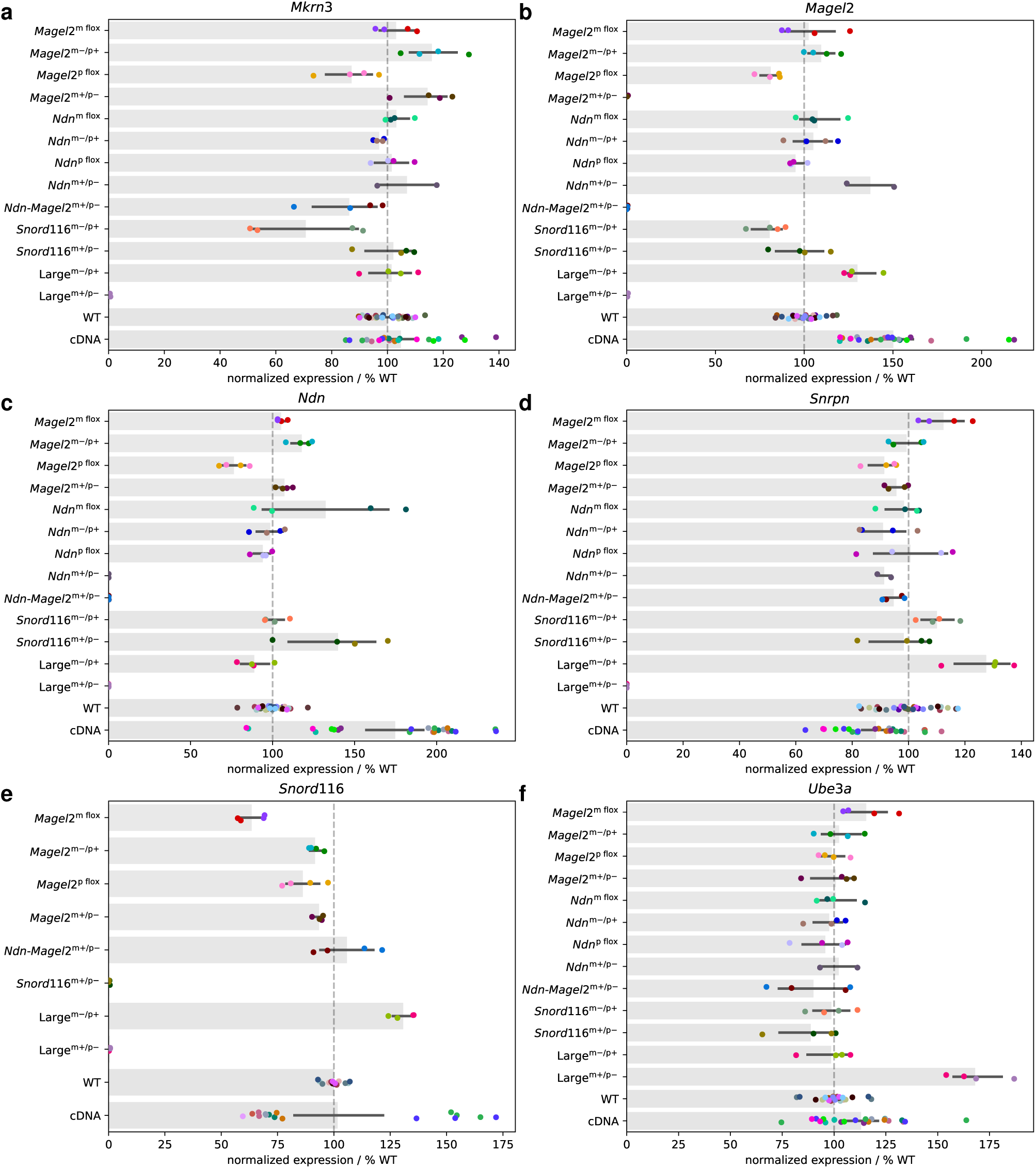
Expression levels of imprinted genes in the PW region from ddPCR experiments. **a**–**f**, Normalized gene expression levels for different mouse lines with maternal deletion (m*−/*p+), paternal deletion (p+/m*−*), floxed maternal allele (m flox), floxed paternal allele (p flox), and no deletion (WT), and cDNA controls, normalized to WT level (100%, dashed, grey line). Single points indicate technical replicates of with biological replicates shown with same color (see legend in Supplementary Fig. 1d).

**Supplementary Fig. 3.**
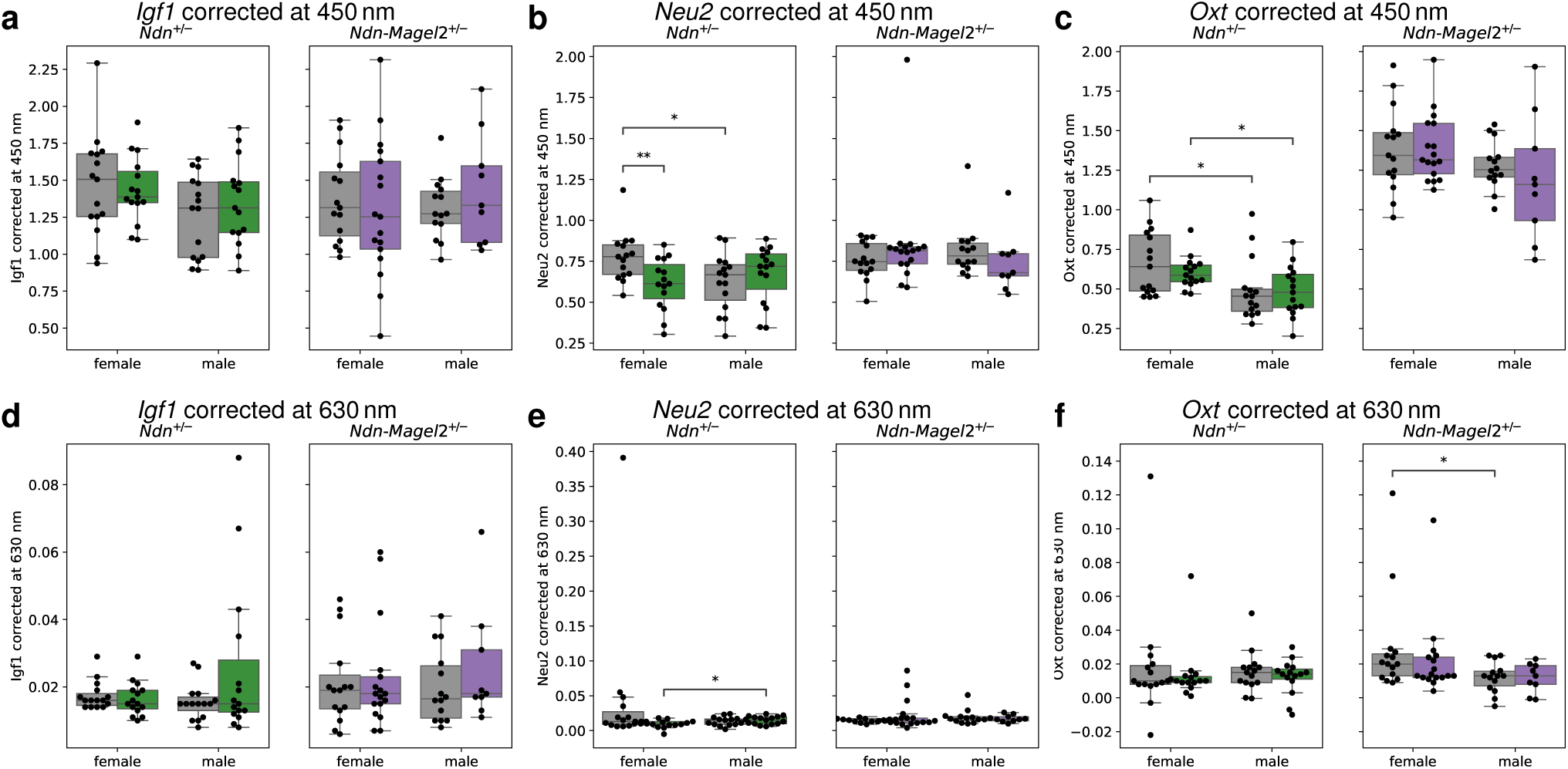
Protein levels from ELISA. The test was performed on *Ndn*^+^*^/−^* and *Ndn*-*Magel2*^+^*^/−^* animals and their WT littermates from pipeline *II*. Corrected protein levels at 450 nm (**a**–**c**) and 630 nm (**d**–**f**) of *Igf1* (**a**,**d**), *Neu2* (**b**,**e**), and *Oxt* (**c**,**f**). Boxplot boxes extend from the first to the third quartile, with a line at the median and whiskers at the fartherst data points within the 1.5 *×* inter-quartile range. Reported *p*-values are calculated using the Mann-Whitney-Wilcoxon test. Right, colored boxes: heterozygous animals; left, grey boxes: their WT littermates; *: *p <* 0.05.

**Supplementary Fig. 4.**
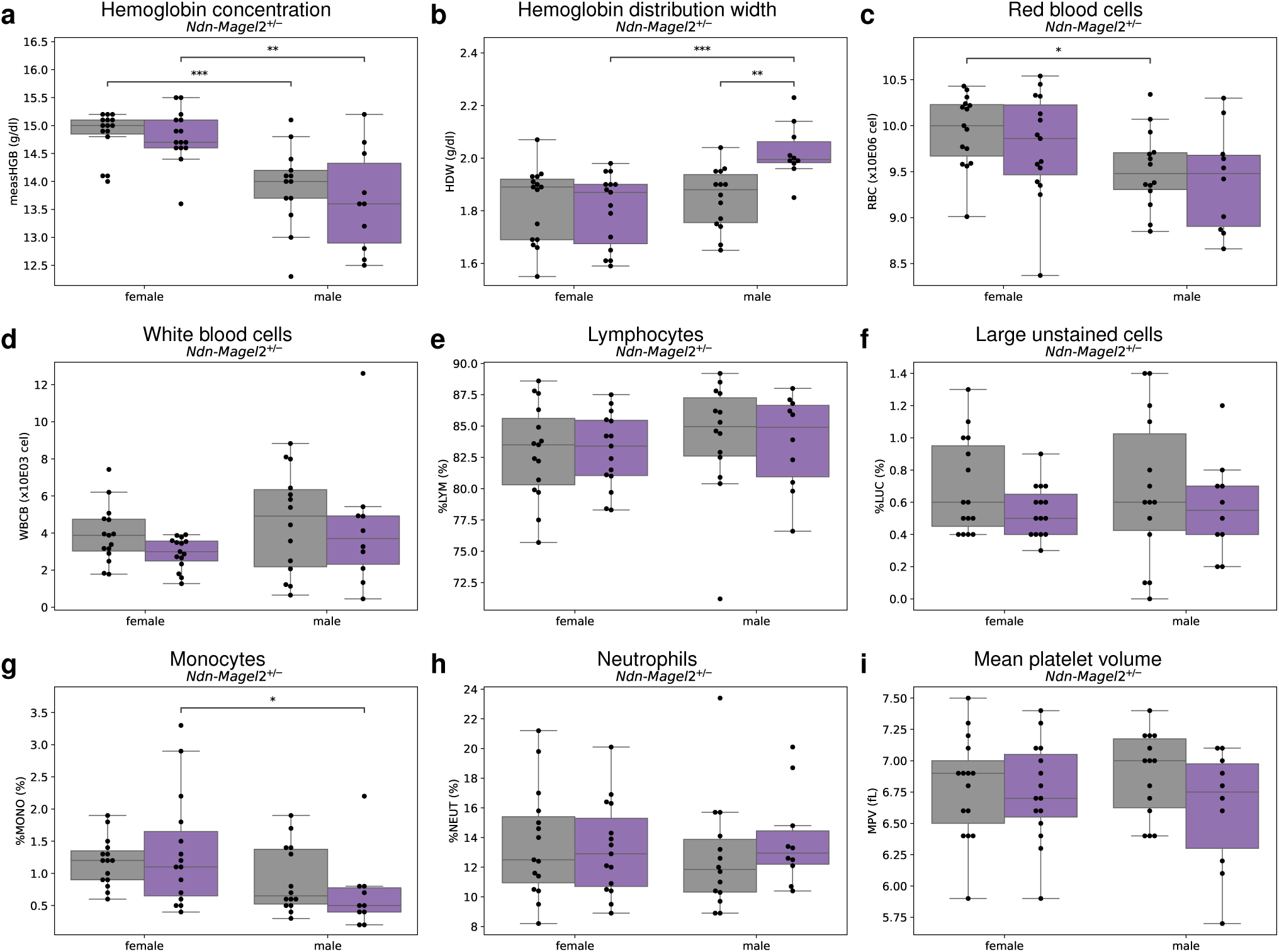
Hematology. The test was performed on *Ndn*-*Magel2* ^+^*^/−^* animals and their WT littermates from pipeline *II*. **a**, Hemoglobin concentration. **b**, Hemoglobin distribution width. **c**, Red blood cell count. **d**, Basophil white blood cell count. **e**, Lymphocytes. **f**, Large unstained cells. **g**, Monocytes. **h**, Neutrophils. **i**, Mean platelet volume. Boxplot boxes extend from the first to the third quartile, with a line at the median and whiskers at the fartherst data points within the 1.5 *×* inter-quartile range. Reported *p*-values are calculated using the Mann-Whitney-Wilcoxon test. Right, colored boxes: heterozygous animals; left, grey boxes: their WT littermates; *: *p <* 0.05; **: *p <* 0.01; ***: *p <* 0.001.

**Supplementary Fig. 5.**
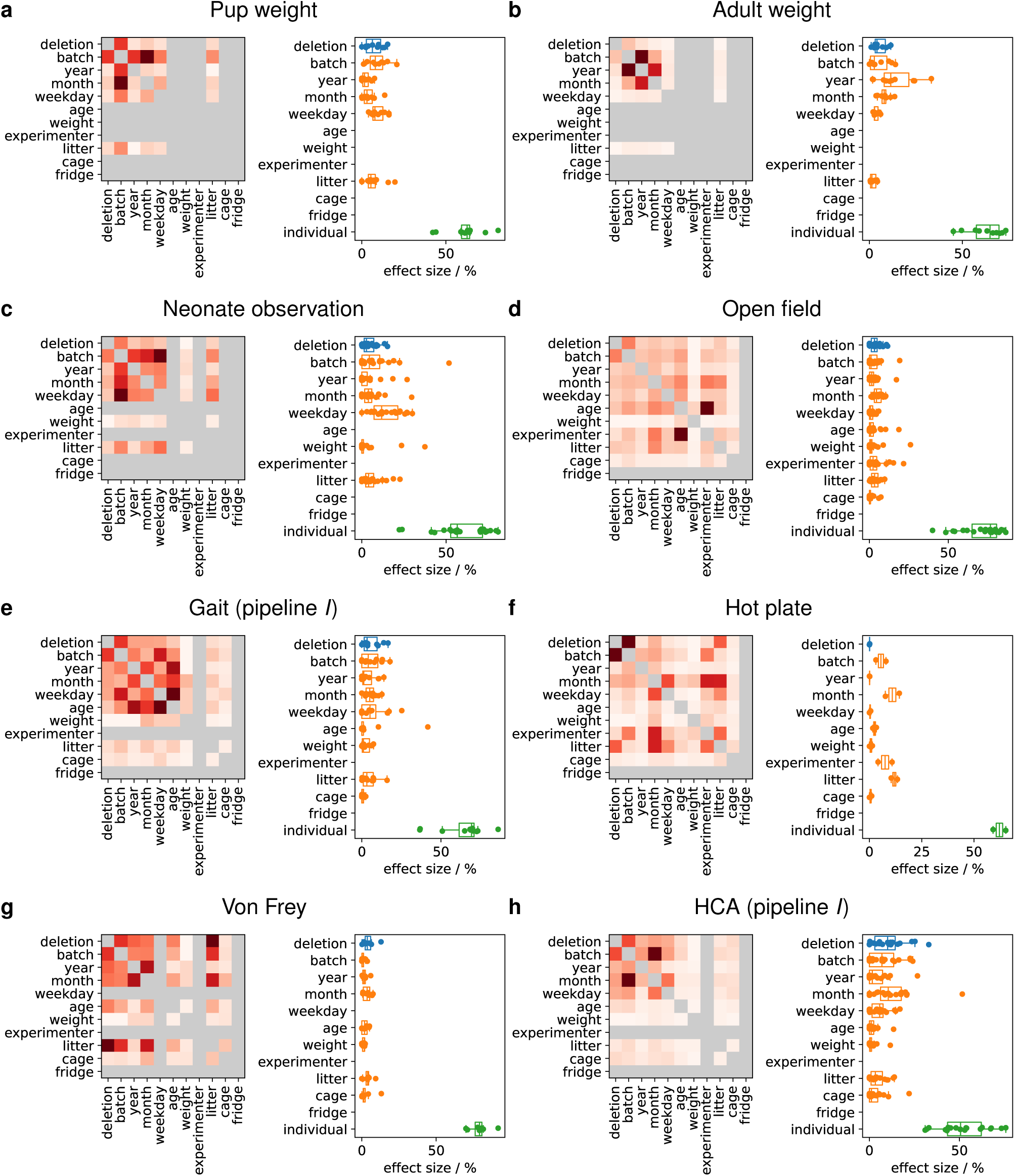
Covariate correlations and effect sizes for body weight measurements and experiments of pipeline *I*. **a**, Pup weight. **b**, Adult weight. **c–h**, Experiments of pipeline *I*.

**Supplementary Fig. 6.**
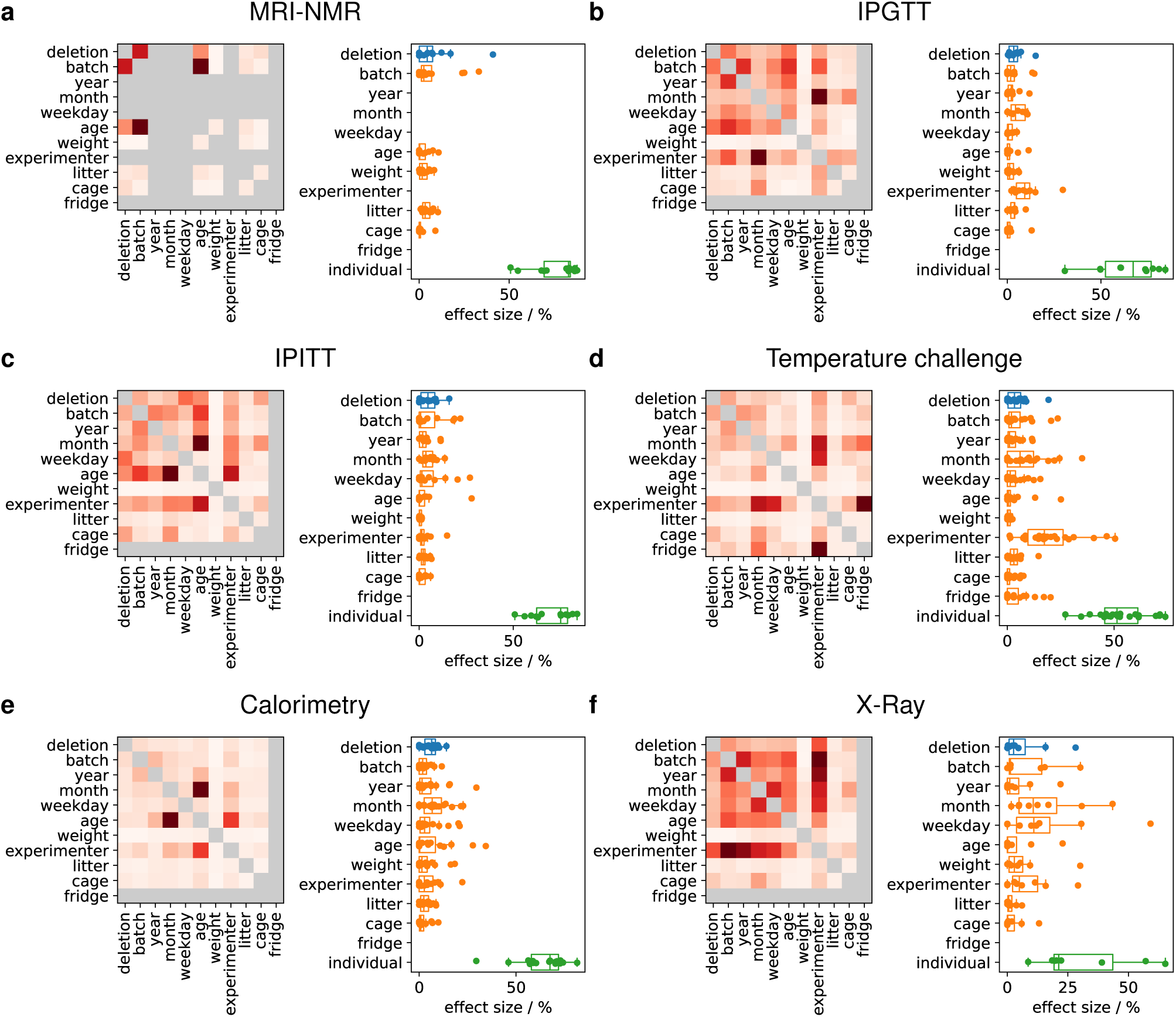
Covariate correlations and effect sizes for experiments of pipeline *II*.

**Supplementary Fig. 7.**
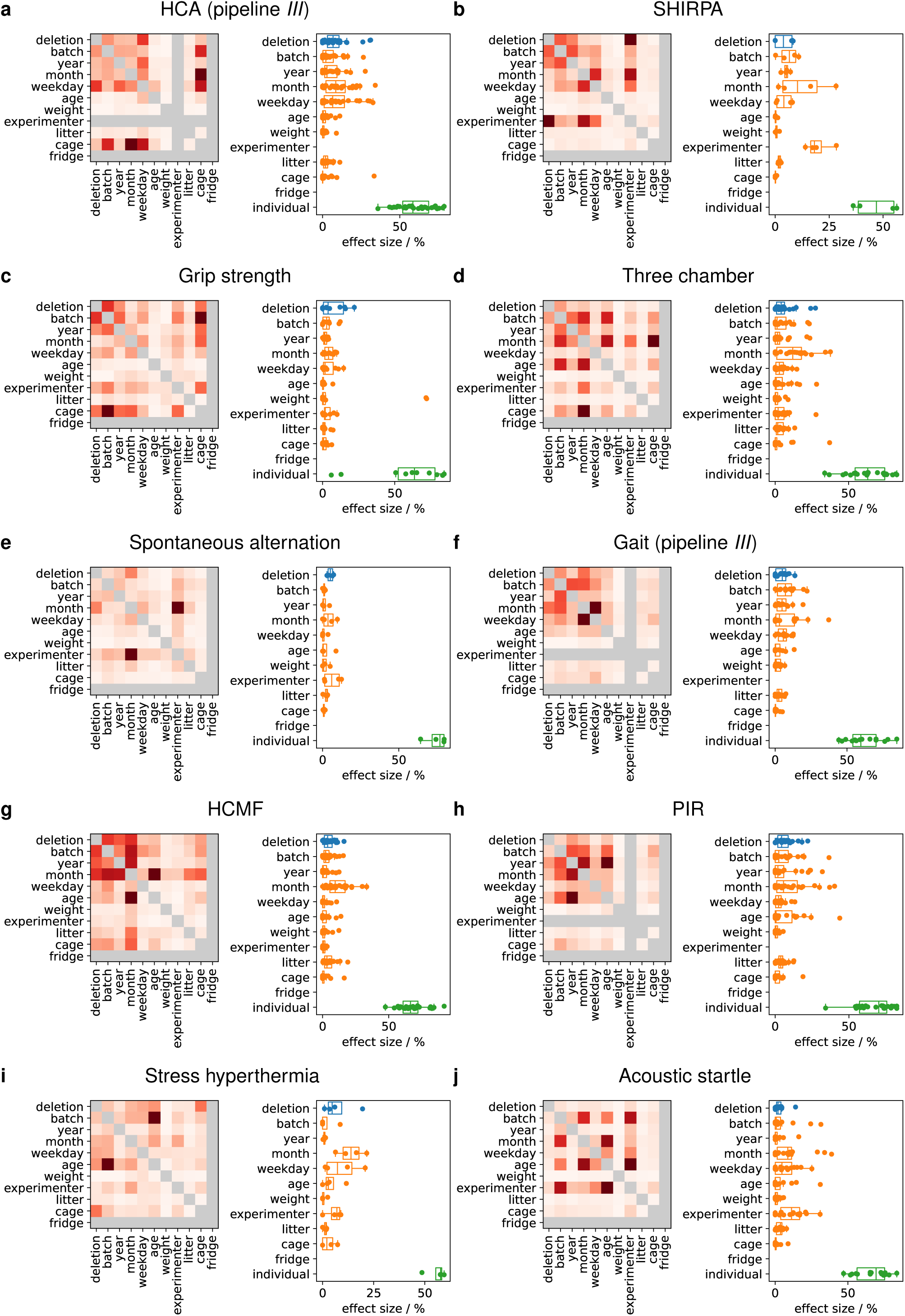
Covariate correlations and effect sizes for experiments of pipeline *III*.

**Supplementary Fig. 8.**
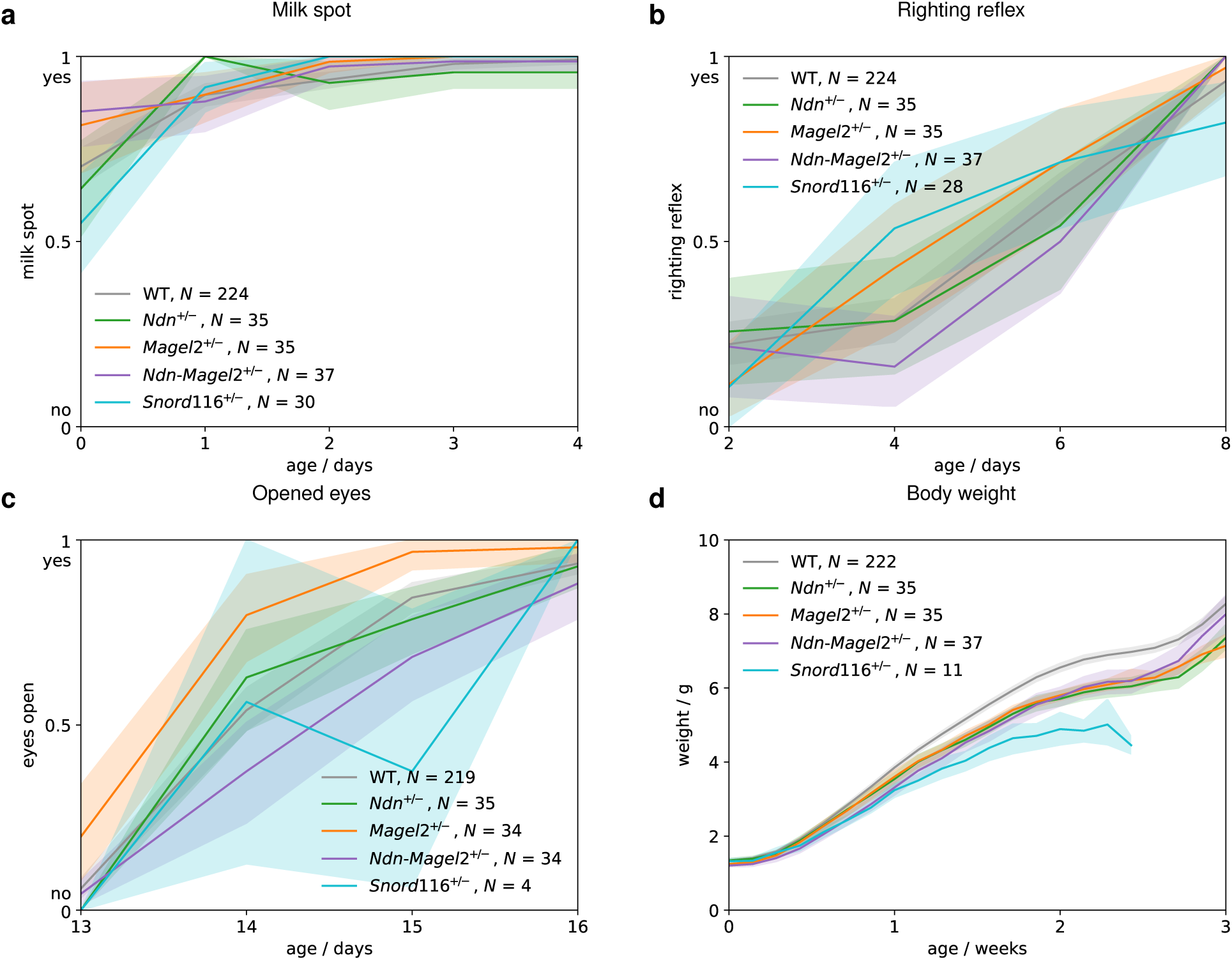
Neonate observation. The experiments were performed on animals from pipeline *I*. **a** Milk spot visibility from birth to postnatal day P4. Animals are included if they survive at least until P4. **b** Righting reflex observed at P2, P4, P6 and P8. Animals are included if they survive at least until P8. **c** Eyes opening from P13 to P16. Animals are included if they survive at least until P16. **d** Measured body weight from birth to P21 (3 weeks of age). Animals are included if they survive at least until P14. The curve for *Snord116*^+^*^/−^* animals ends at P17 because either animals did not survive or their weight was not measured after P17. The line and upper and lower bounds of the shaded area correspond to the mean *±* 95% bootstrap confidence interval.

**Supplementary Fig. 9.**
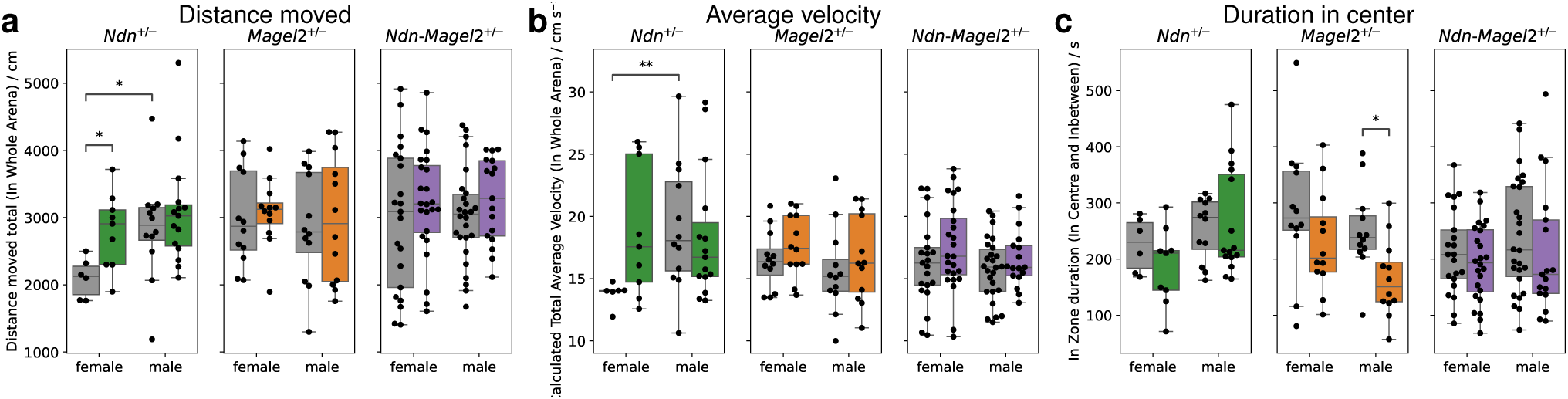
Open field. The experiments were performed on animals from pipeline *I* at the age of around 4 weeks. **a** Total distance moved. **a** Average velocity. **a** Duration in central area. Boxplot boxes extend from the first to the third quartile, with a line at the median and whiskers at the fartherst data points within the 1.5 *×* inter-quartile range. Reported *p*-values are calculated using the Mann-Whitney-Wilcoxon test. Right, colored boxes: heterozygous animals; left, grey boxes: their WT littermates; *: *p <* 0.05; **: *p <* 0.01.

**Supplementary Fig. 10.**
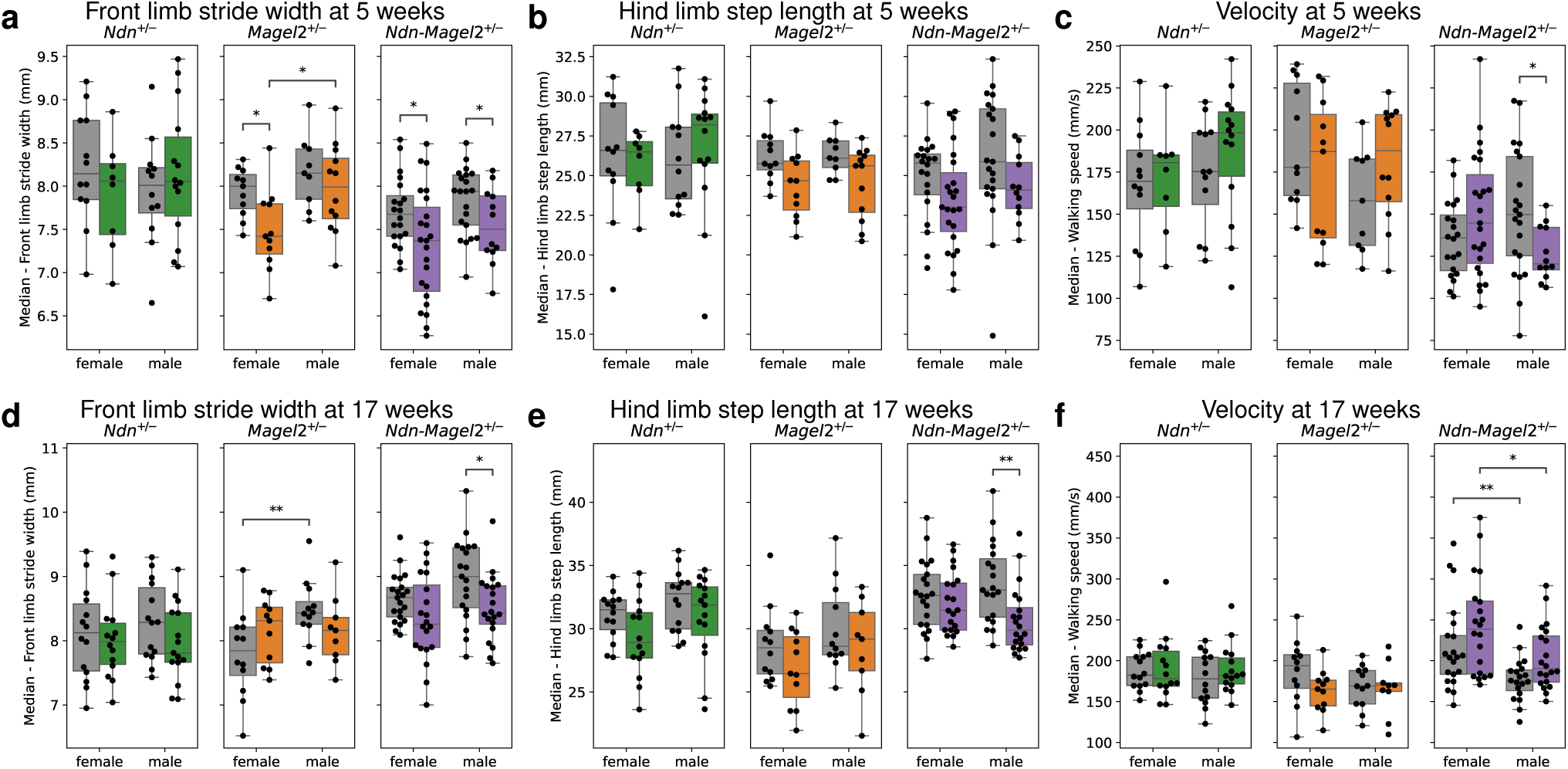
Gait. **a**–**c**, Gait parameters for animals at the age of around 5 weeks from pipeline *I*. **d**–**f**, Gait parameters for animals at the age of around 17 weeks from pipeline *III*. **a**,**d**, Front limb stride width. **b**,**e**, Hind limb step length. **c**,**f**, Velocity. Boxplot boxes extend from the first to the third quartile, with a line at the median and whiskers at the fartherst data points within the 1.5 *×* inter-quartile range. Reported *p*-values are calculated using the Mann-Whitney-Wilcoxon test. Right, colored boxes: heterozygous animals; left, grey boxes: their WT littermates; *: *p <* 0.05; **: *p <* 0.01.

**Supplementary Fig. 11.**
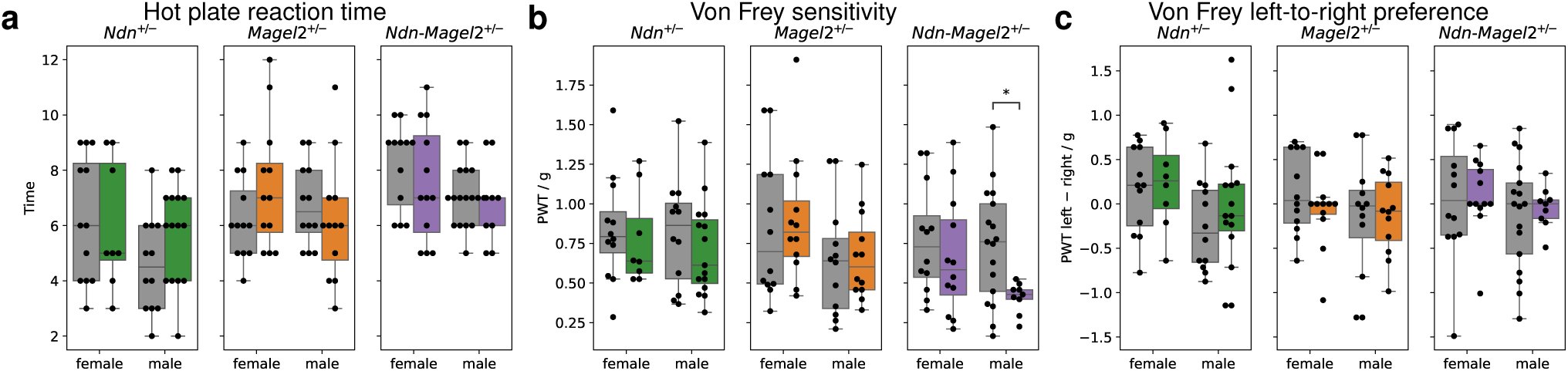
Hot plate and Von Frey sensitivity. **a**, Time until reaction on hot plate for animals at the age of around 10 weeks from pipeline *I*. **b**,**c**, Mean sensitivity (**b**) and left-to-right preference (**c**) measured with Von Frey experiments for animals at the age of around 11 weeks from pipeline *I*. The estimates are averaged over the experimental days 1 and 2 (**b**,**c**), and over left and right foot (**b**). Boxplot boxes extend from the first to the third quartile, with a line at the median and whiskers at the fartherst data points within the 1.5 *×* inter-quartile range. Reported *p*-values are calculated using the Mann-Whitney-Wilcoxon test. Right, colored boxes: heterozygous animals; left, grey boxes: their WT littermates; *: *p <* 0.05.

**Supplementary Fig. 12.**
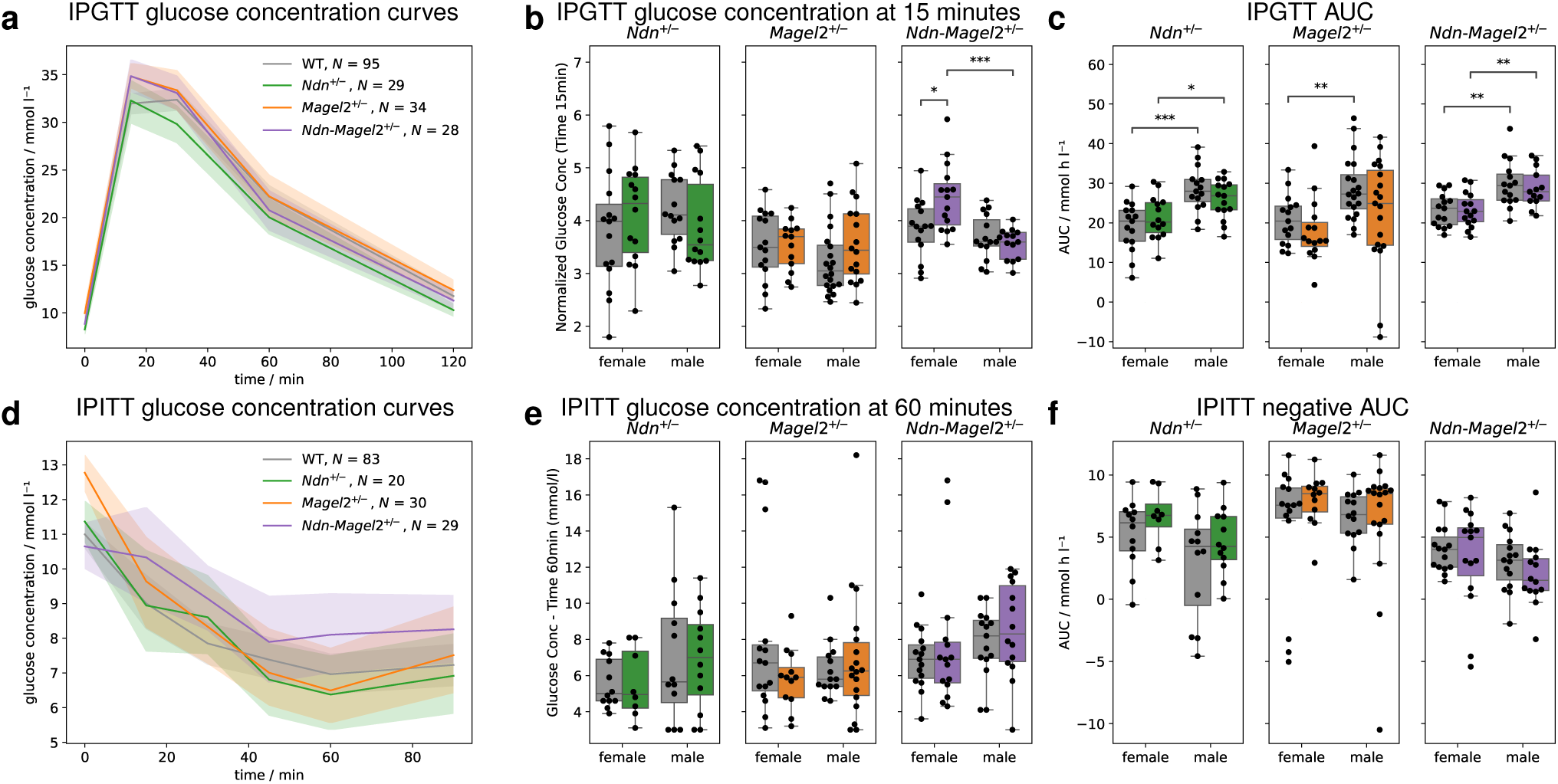
Intraperitoneal tolerance tests for glucose (IPGTT) and insulin (IPITT). **a**–**c**, IPGTT for over-night fasted animals at the age of around 18 weeks from pipeline *II* : glucose concentration curves (**a**), glucose concentration at 15 minutes (**b**), and area under the curve (AUC) of the glucose concentration curves (**c**). **d**–**f**, IPITT for animals at the age of around 21 weeks from pipeline *II* : glucose concentration curves (**d**), glucose concentration at 15 minutes (**e**), and negative AUC of the glucose concentration curves (**f**). The line and upper and lower bounds of the shaded area correspond to the mean *±* 95% bootstrap confidence interval. Boxplot boxes in **b**,**c**,**e**,**f** extend from the first to the third quartile, with a line at the median and whiskers at the fartherst data points within the 1.5 *×* inter-quartile range. Reported *p*-values are calculated using the Mann-Whitney-Wilcoxon test. Right, colored boxes: heterozygous animals; left, grey boxes: their WT littermates; *: *p <* 0.05; **: *p <* 0.01; ***: *p <* 0.001.

**Supplementary Fig. 13.**
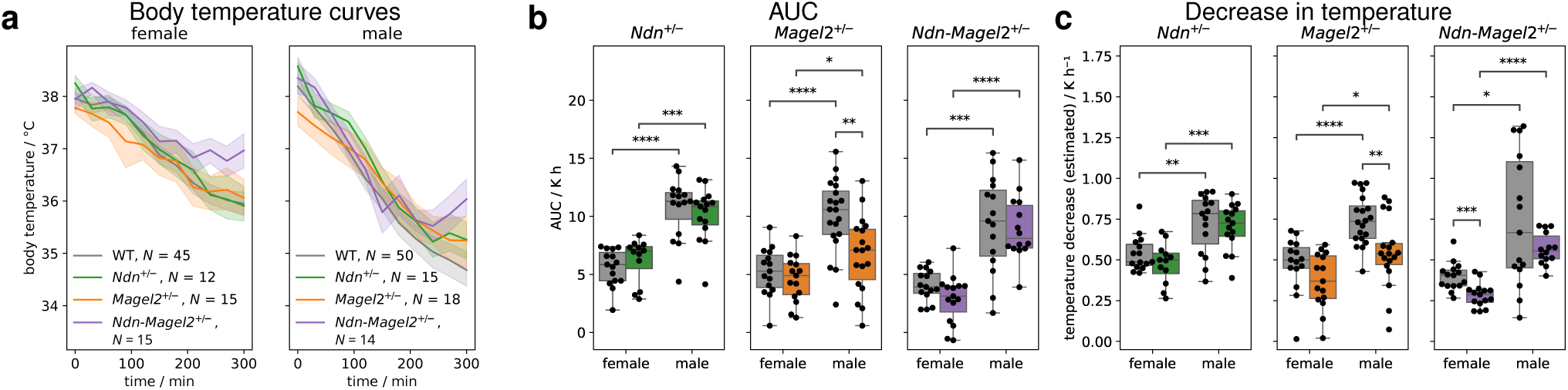
Temperature challenge. The experiments were performed on animals from pipeline *II* at the age of around 23 weeks. **a** Body temperature curves of females (left) and males (right) during 5 hours in fridge. **b** Negative area under the curve (AUC) of the body temperature curves. **c** Estimated decrease in temperature. Boxplot boxes in **b**,**c** extend from the first to the third quartile, with a line at the median and whiskers at the fartherst data points within the 1.5 *×* inter-quartile range. Reported *p*-values are calculated using the Mann-Whitney-Wilcoxon test. Right, colored boxes: heterozygous animals; left, grey boxes: their WT littermates; *: *p <* 0.05; **: *p <* 0.01; ***: *p <* 0.001; ****: *p <* 0.0001.

**Supplementary Fig. 14.**
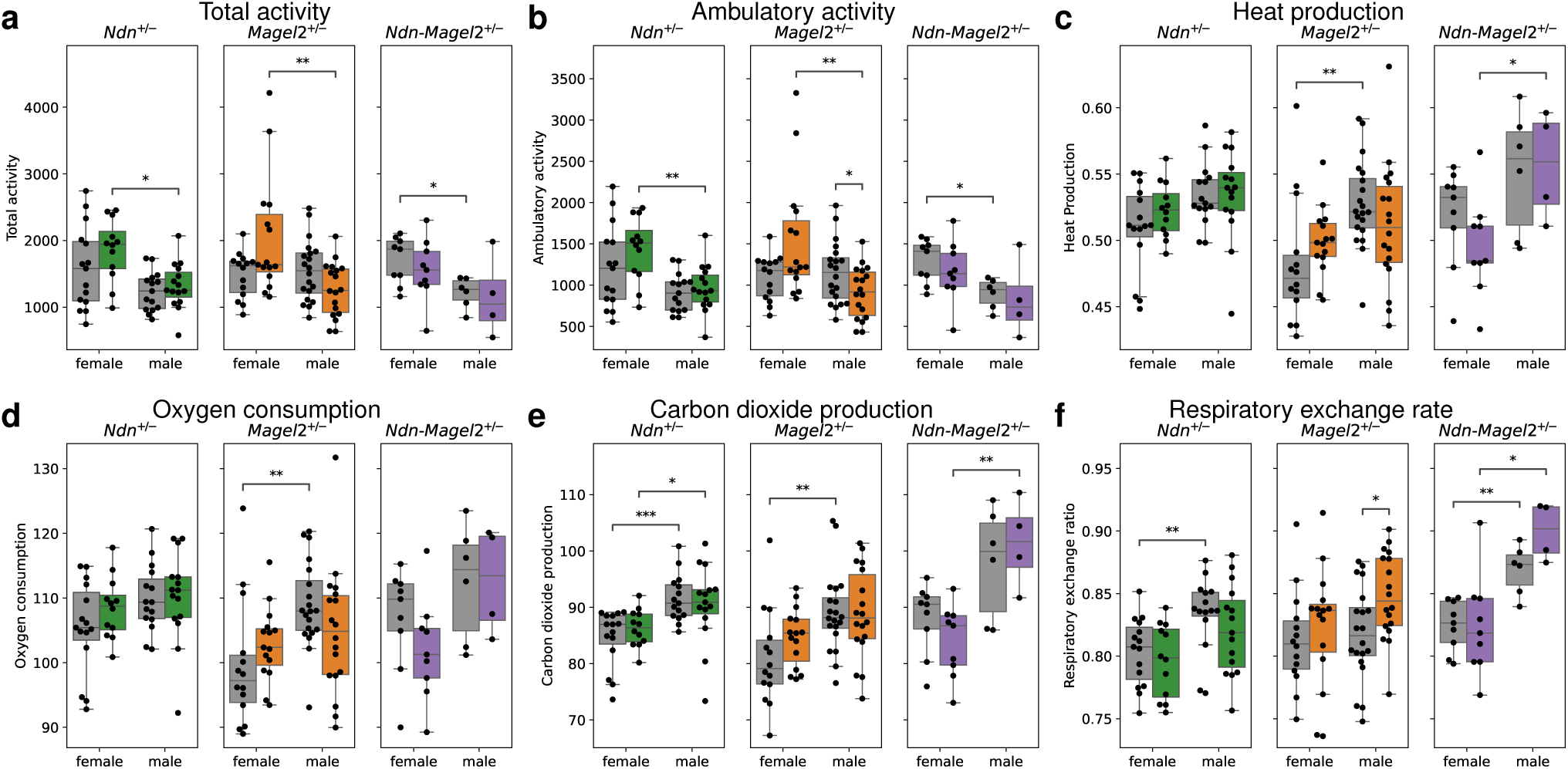
Calorimetry. The experiments were performed on animals from pipeline *II* at the age of around 26 weeks. **a**, Total activity. **b**, Ambulatory activity. **c**, Heat production. **d**, Oxygen consumption. **e**, Carbon dioxide production. **f**, Respiratory exchange rate. Boxplot boxes extend from the first to the third quartile, with a line at the median and whiskers at the fartherst data points within the 1.5 *×* inter-quartile range. Reported *p*-values are calculated using the Mann-Whitney-Wilcoxon test. Right, colored boxes: heterozygous animals; left, grey boxes: their WT littermates; *: *p <* 0.05; **: *p <* 0.01; ***: *p <* 0.001.

**Supplementary Fig. 15.**
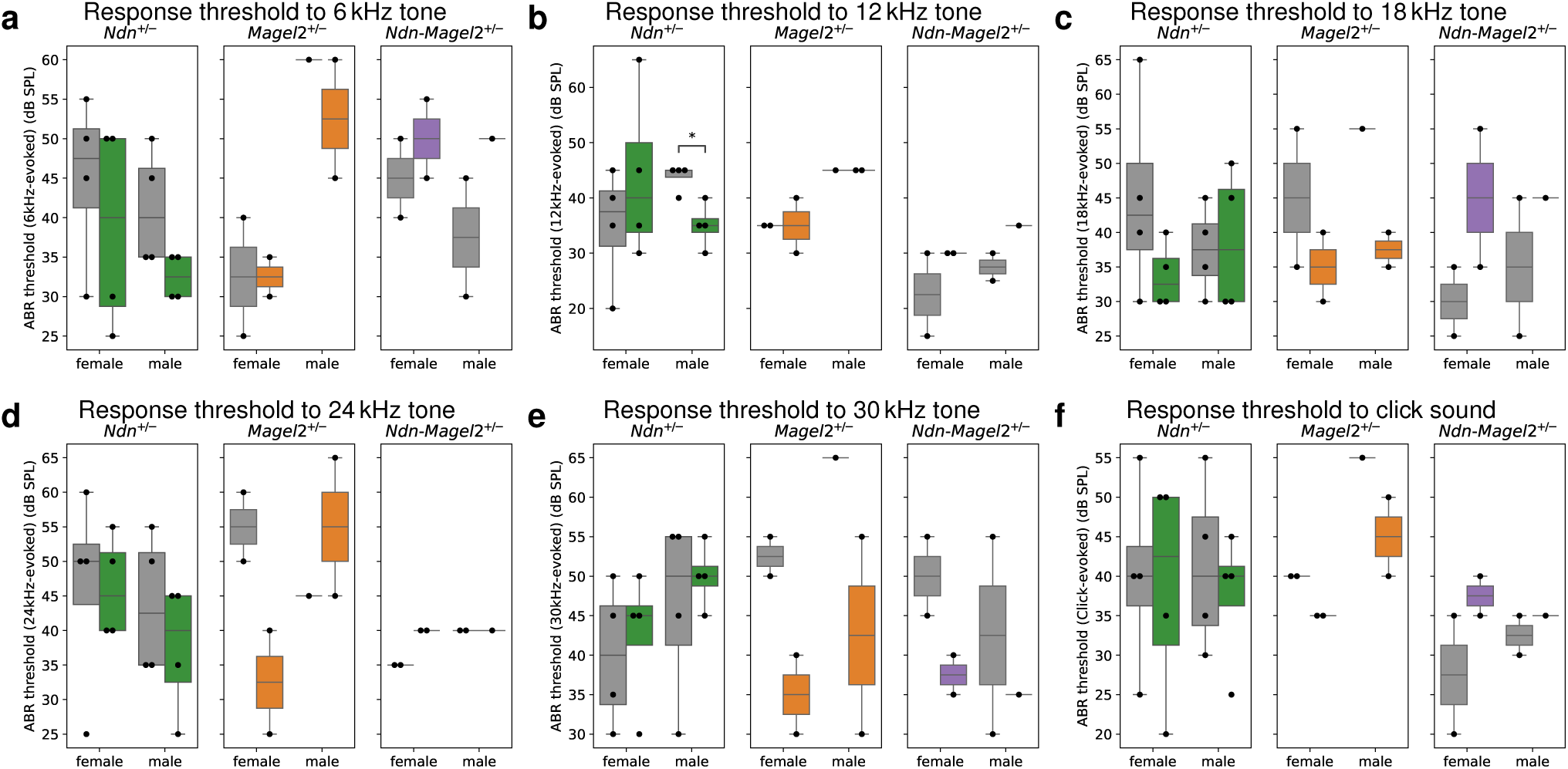
Auditory brain stem response (ABR). The experiments were performed on animals from pipeline *II* at the age of around 26 weeks. **a**–**f** Thresholds of EEG response to tones with frequencies of 6 kKz (**a**), 12 kKz (**b**), 18 kKz (**c**), 24 kKz (**d**), 30 kKz (**e**), and to click sound (**f**). Boxplot boxes extend from the first to the third quartile, with a line at the median and whiskers at the fartherst data points within the 1.5 *×* inter-quartile range. Reported *p*-values are calculated using the Mann-Whitney-Wilcoxon test. Right, colored boxes: heterozygous animals; left, grey boxes: their WT littermates; *: *p <* 0.05.

**Supplementary Fig. 16.**
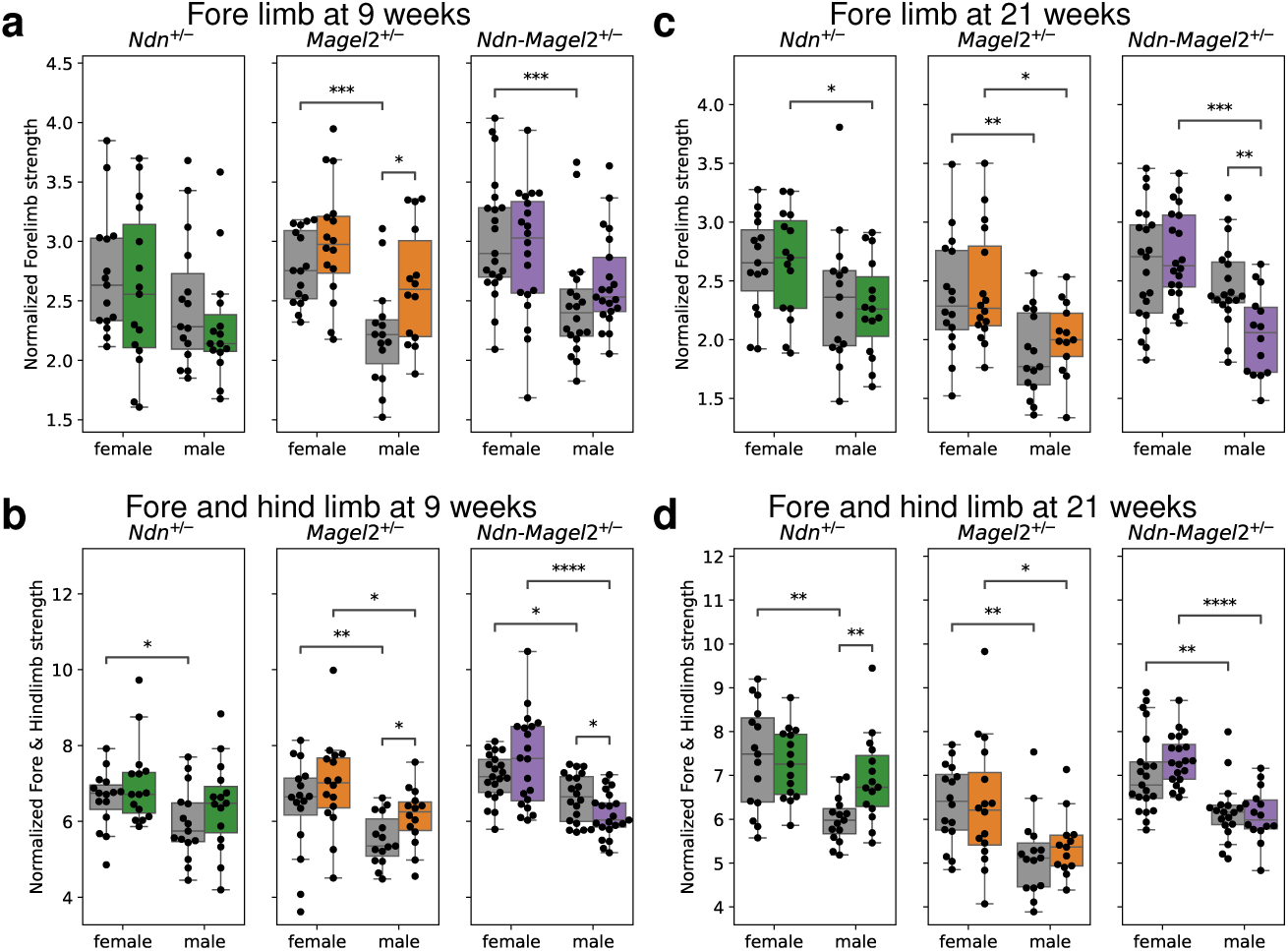
Grip strength. **a**,**b**, Normalized grip strength for animals from pipeline *III* at the age of around 9 weeks of fore limb only (**a**) and of fore and hind limb (**b**). **c**,**d**, Normalized grip strength for the same animals as in **a**,**b** at age of around 21 weeks of fore limb only (**c**) and of fore and hind limb (**d**). Boxplot boxes extend from the first to the third quartile, with a line at the median and whiskers at the fartherst data points within the 1.5 *×* inter-quartile range. Reported *p*-values are calculated using the Mann-Whitney-Wilcoxon test. Right, colored boxes: heterozygous animals; left, grey boxes: their WT littermates; *: *p <* 0.05; **: *p <* 0.01; ***: *p <* 0.001; ****: *p <* 0.0001.

**Supplementary Fig. 17.**
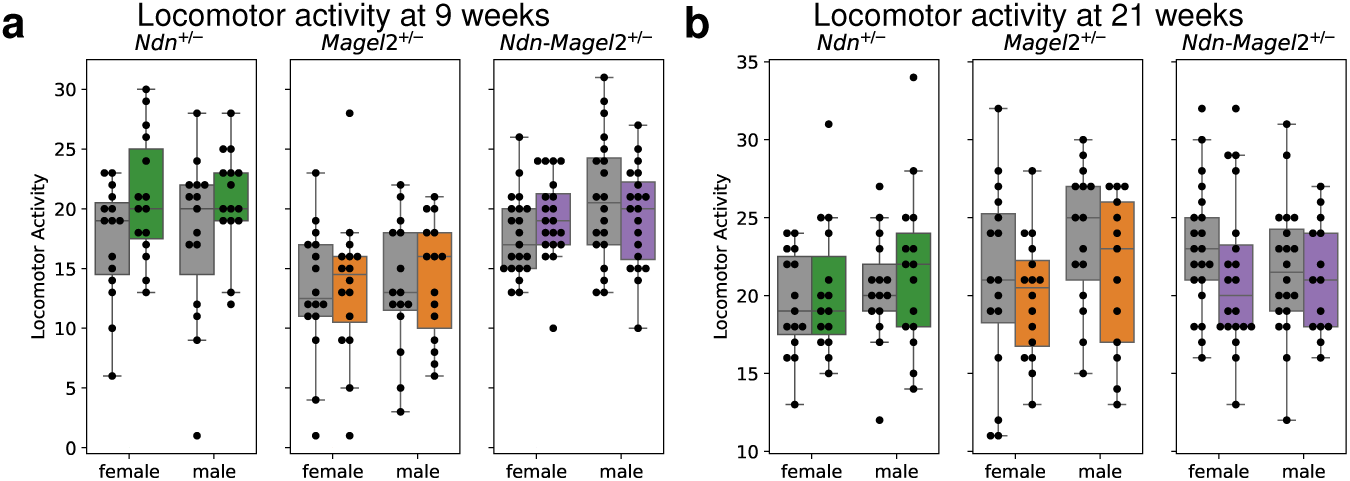
Locomotor activity from SHIRPA assessment. **a**, Locomotor activity for animals from pipeline *III* at the age of around 9 weeks. **a**, Locomotor activity for the same animals at the age of around 21 weeks. Boxplot boxes extend from the first to the third quartile, with a line at the median and whiskers at the fartherst data points within the 1.5 *×* inter-quartile range. Reported *p*-values are calculated using the Mann-Whitney-Wilcoxon test. Right, colored boxes: heterozygous animals; left, grey boxes: their WT littermates.

**Supplementary Fig. 18.**
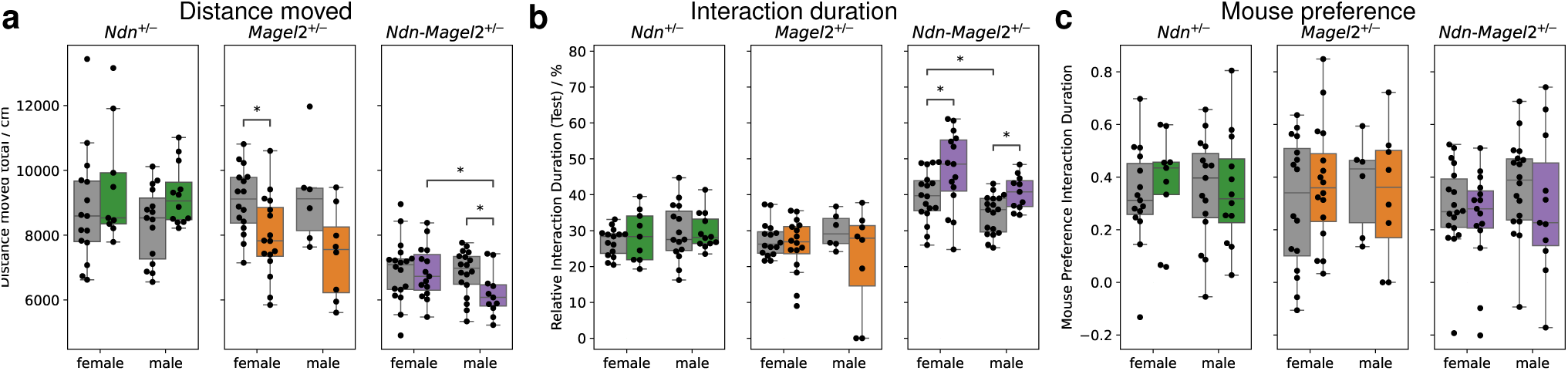
Three chamber. The experiments were performed on animals from pipeline *III* at the age of around 14 weeks. **a**, Total distance moved during habituation and test. **b**, Relative duration of interaction with the stranger mouse or the object during the test phase. **c**, Preference for interacting with the stranger mouse over the object during the test phase, where 1 means interaction only with the mouse, *−*1 interaction only with the object and 0 same interaction time with both. Boxplot boxes extend from the first to the third quartile, with a line at the median and whiskers at the fartherst data points within the 1.5 *×* inter-quartile range. Reported *p*-values are calculated using the Mann-Whitney-Wilcoxon test. Right, colored boxes: heterozygous animals; left, grey boxes: their WT littermates; *: *p <* 0.05.

**Supplementary Fig. 19.**
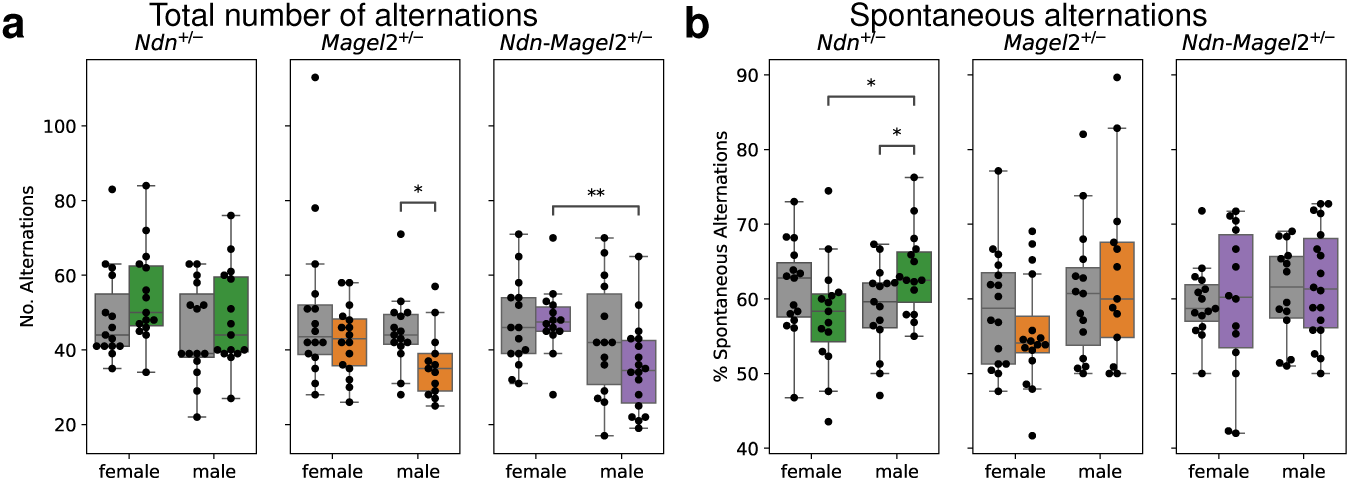
Spontaneous alternation. The experiments were performed on animals from pipeline *III* at the age of around 16 weeks. **a**, Total number of alternations between arms of the Y-maze. **b**, Percentage of spontaneous alternations. Boxplot boxes extend from the first to the third quartile, with a line at the median and whiskers at the fartherst data points within the 1.5 *×* inter-quartile range. Reported *p*-values are calculated using the Mann-Whitney-Wilcoxon test. Right, colored boxes: heterozygous animals; left, grey boxes: their WT littermates; *: *p <* 0.05; **: *p <* 0.01.

**Supplementary Fig. 20.**
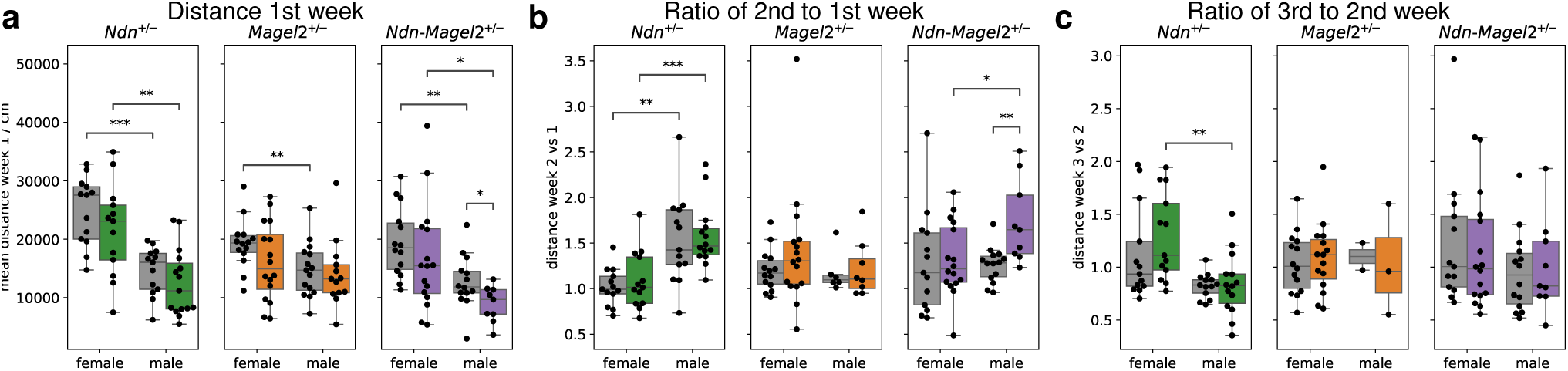
Home-cage motor functions (HCMF). The experiments were performed on animals from pipeline *III* at the age of around 23 weeks. Between the 2nd and the 3nd week the standard wheel was replaced by a wheel with irregular gaps. **a**, Mean distance ran on the wheel during the 1st week. **b**, Ratio of daily ran distance of 2nd to 1st week. **b**, Ratio of daily ran distance of 3rd to 2nd week. Boxplot boxes extend from the first to the third quartile, with a line at the median and whiskers at the fartherst data points within the 1.5 *×* inter-quartile range. Reported *p*-values are calculated using the Mann-Whitney-Wilcoxon test. Right, colored boxes: heterozygous animals; left, grey boxes: their WT littermates; *: *p <* 0.05; **: *p <* 0.01; ***: *p <* 0.001.

**Supplementary Fig. 21.**
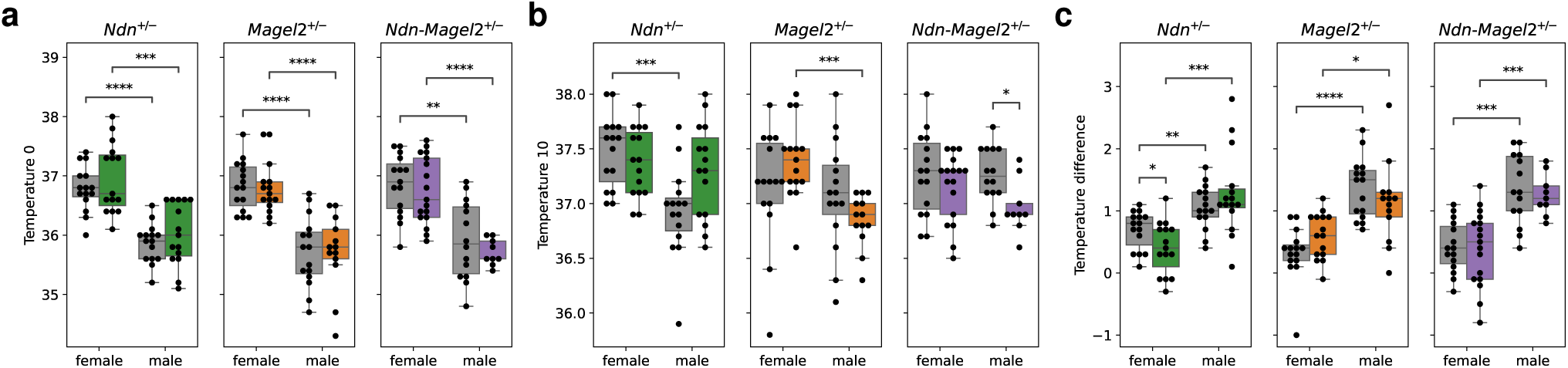
Stress induced hyperthermia. The experiments were performed on animals from pipeline *III* at the age of around 32 weeks. **a**, Body temperature at start of the experiment. **b**, Body temperature at the end of the experiments after 10 minutes. **c**, Temperature difference from 0 to 10 minutes. Boxplot boxes extend from the first to the third quartile, with a line at the median and whiskers at the fartherst data points within the 1.5 *×* inter-quartile range. Reported *p*-values are calculated using the Mann-Whitney-Wilcoxon test. Right, colored boxes: heterozygous animals; left, grey boxes: their WT littermates; *: *p <* 0.05; **: *p <* 0.01; ***: *p <* 0.001; ****: *p <* 0.0001.

**Supplementary Fig. 22.**
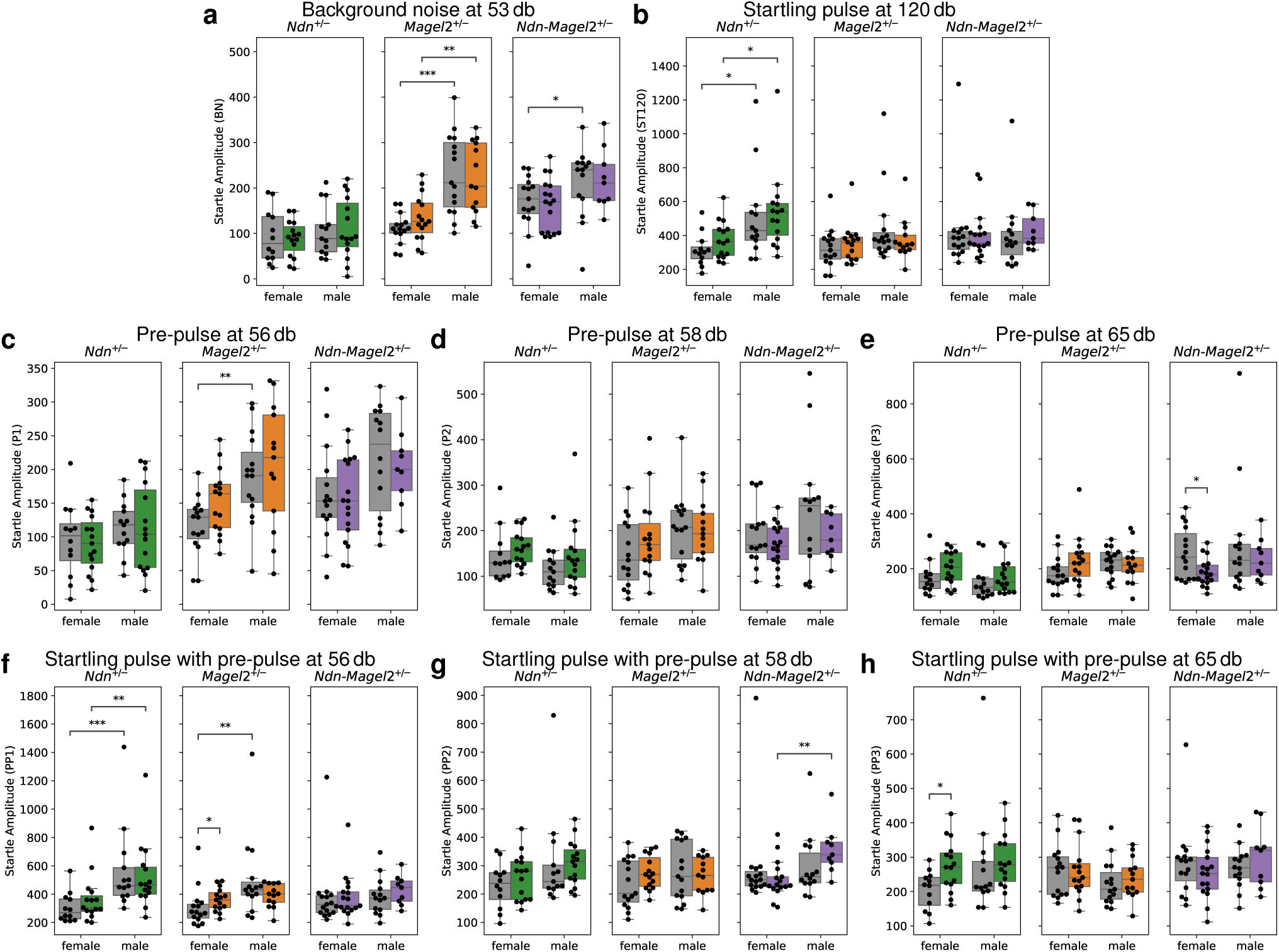
Acoustic startle. The experiments were performed on animals from pipeline *III* at the age of around 33 weeks. **a**–**h**, Startle amplitudes for background noise at 53 db (**a**) and startling pulse at 120 db (**b**), pre-pulses at 56 db (**c**), 58 db (**d**) and 65 db (**e**), and startling pulses preceded by a pre-pulse at 56 db (**f**), 58 db (**g**) and 65 db (**h**). Boxplot boxes extend from the first to the third quartile, with a line at the median and whiskers at the fartherst data points within the 1.5 *×* inter-quartile range. Reported *p*-values are calculated using the Mann-Whitney-Wilcoxon test. Right, colored boxes: heterozygous animals; left, grey boxes: their WT littermates; *: *p <* 0.05; **: *p <* 0.01; ***: *p <* 0.001.

**Supplementary Fig. 23.**
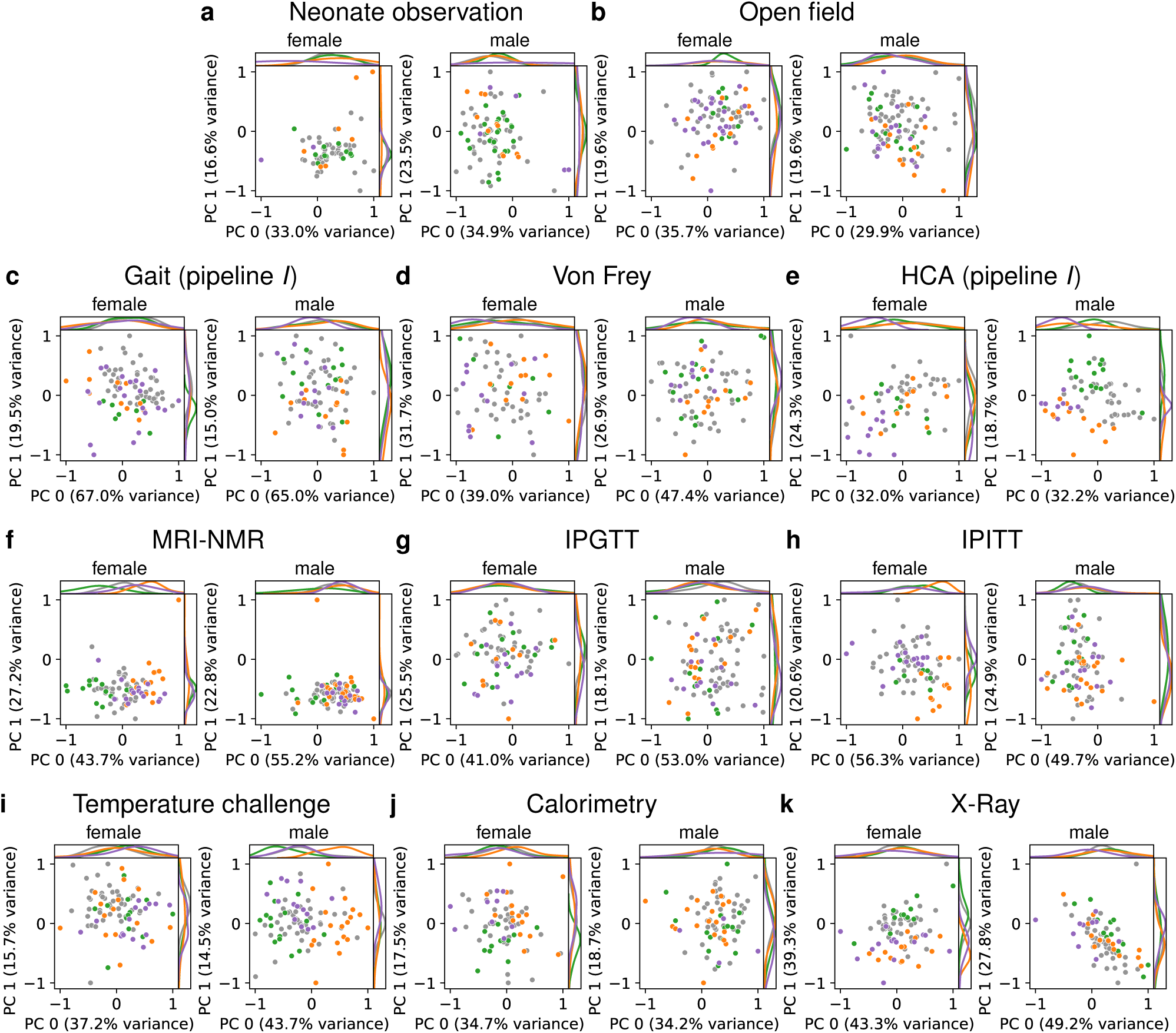
Single-experiment PCAs for experiments of pipelines *I* and *II*. The distributions are shown of the AE estimates of the single-experiment principal components, removing the common environment, for females (left) and males (right). Colors of points and estimated distribution lines are: green for *Ndn*^+^*^/−^*, orange for *Magel2*^+^*^/−^*, purple for *Ndn*-*Magel2*^+^*^/−^*, and gray for WT animals.**a**–**e**, Experiments of pipeline *I*. **f**–**k**, Experiments of pipeline *II*.

**Supplementary Fig. 24.**
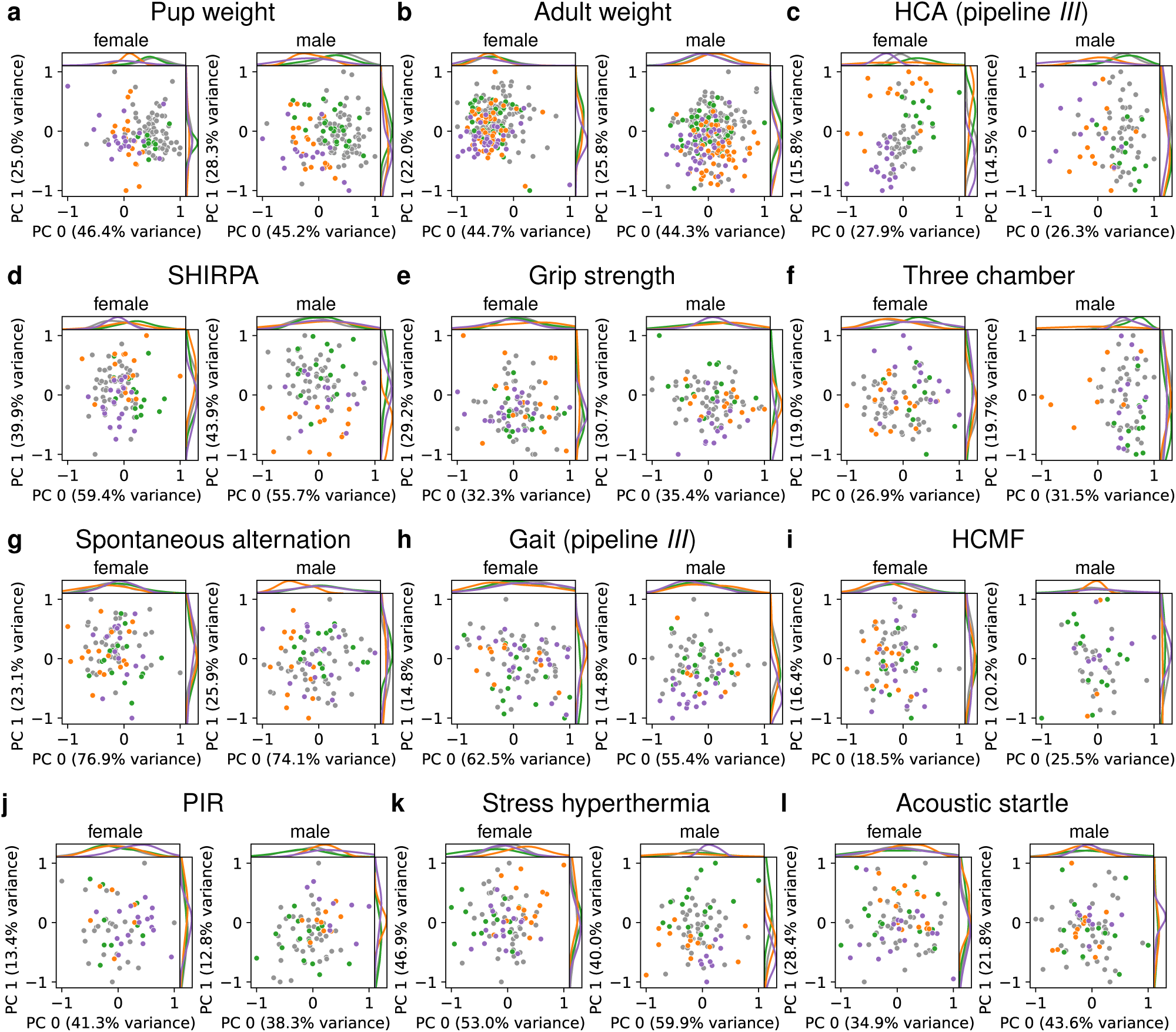
Single-experiment PCAs for body weight measurements and experiments of pipeline *III*. The distributions are shown of the AE estimates of the single-experiment principal components, removing the common environment, for females (left) and males (right). Colors of points and estimated distribution lines are: green for *Ndn*^+^*^/−^*, orange for *Magel2*^+^*^/−^*, purple for *Ndn*-*Magel2*^+^*^/−^*, and gray for WT animals.**a**, Pup weight. **b**, Adult weight. **c**–**l**, Experiments of pipeline *III*.

**Supplementary Fig. 25.**
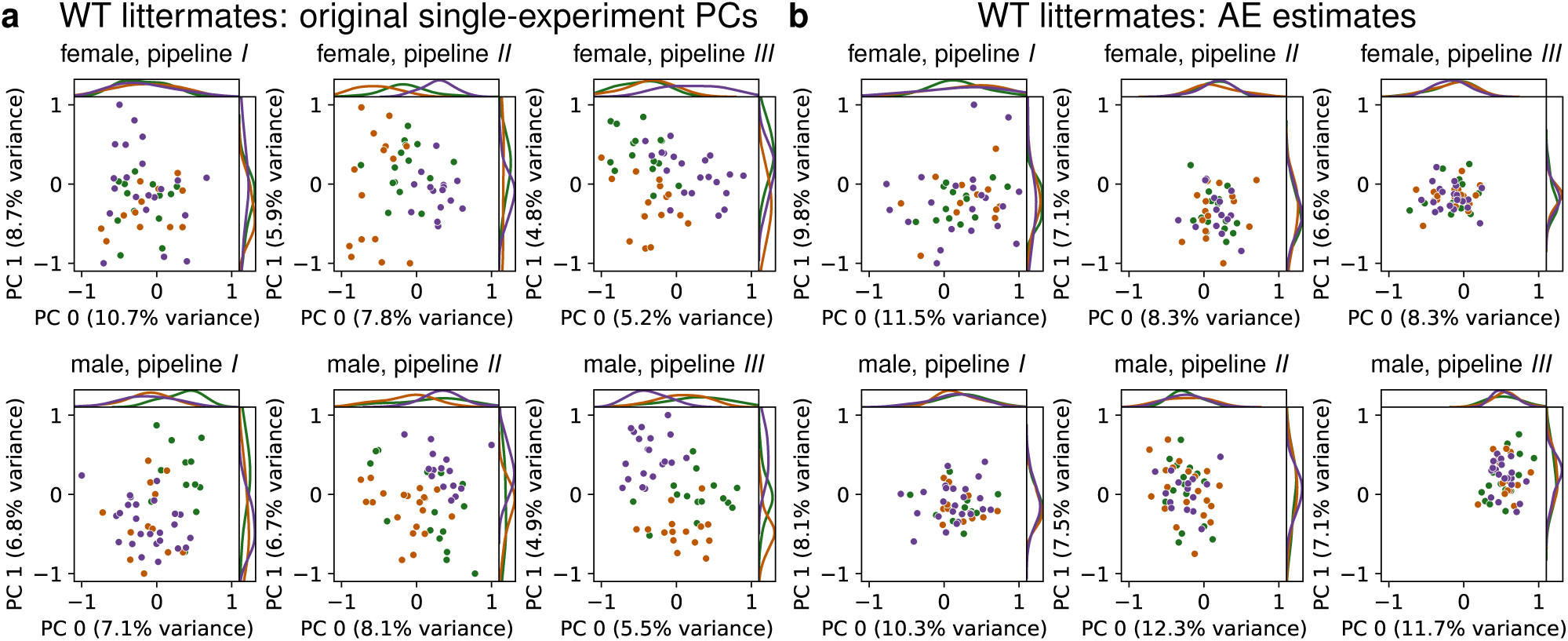
Principal component distributions for WT littermates. The final per-pipeline PCAs are trained on the single-experiment PCAs. The first two principal components (PC 0 and PC 1) for female (top) and male (bottom) WT littermates are shown. Colors of points and estimated distribution lines are: green for *Ndn*^+^*^/−^*, orange for *Magel2*^+^*^/−^*, purple for *Ndn*-*Magel2*^+^*^/−^*, and gray for WT animals.**a**, The original single-experiment PCs are used as input to the per-pipeline PCAs. **b**, The AE estimates (removing the common environment) of these single-experiment PCs are used as input to the per-pipeline PCAs, effectively removing the differences between WT animals of different batches.

**Supplementary Fig. 26.**
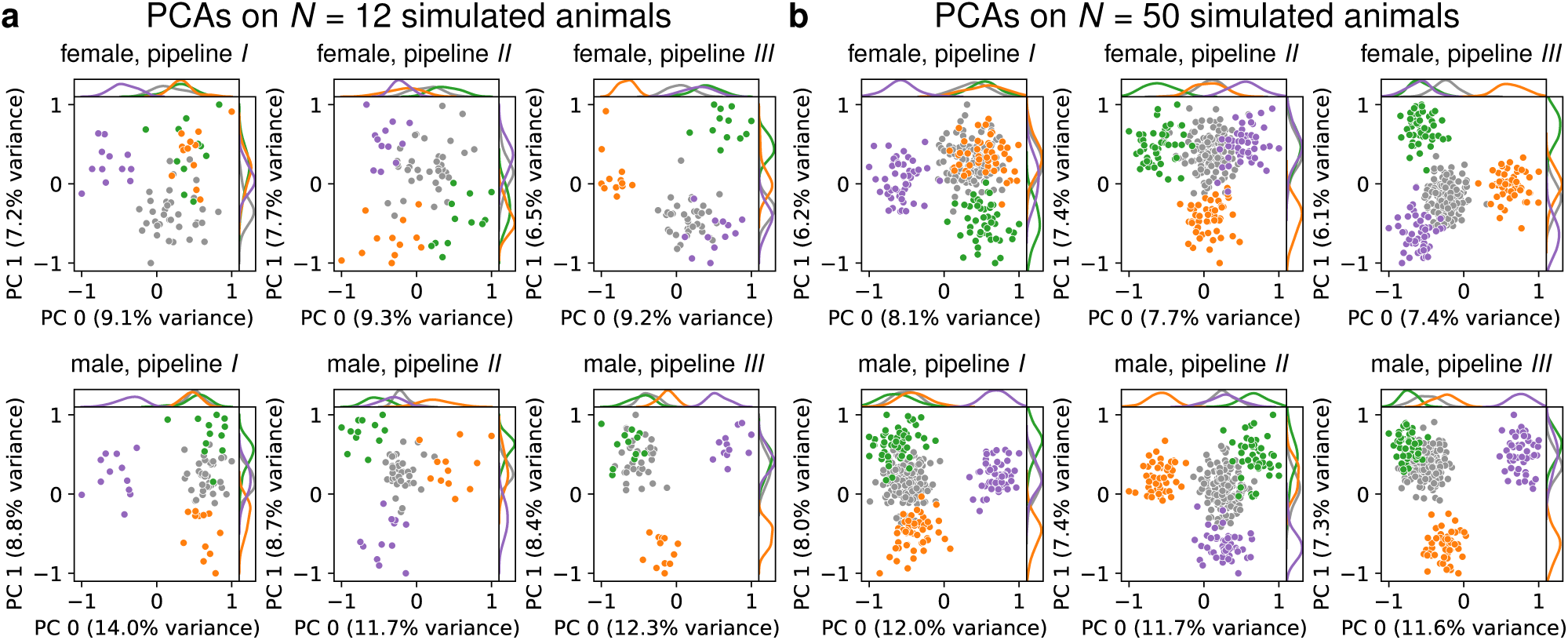
Single-pipeline PCAs on simulated animals. PCAs for the three pipelines for observations of females (top) and males (bottom) from pipeline *I* (left), pipeline *II* (middle) and pipeline *III* (right). **a**, *N* = 12 simulated animals per group (6 groups: *Ndn*^+^*^/−^*, *Magel2*^+^*^/−^* and *Ndn*-*Magel2*^+^*^/−^*, and their respective WT-littermate groups). **b**, *N* = 50 simulated animals per group. Colors of points and estimated distribution lines are: green for *Ndn*^+^*^/−^*, orange for *Magel2*^+^*^/−^*, purple for *Ndn*-*Magel2*^+^*^/−^*, and gray for WT animals.

