## Supplemental information for "The Preclinical Animal Network (PCAN): Integrative high-throughput phenotyping of standardized mouse models for Prader-Willi syndrome"

MRC | Harwell

### *Ndn* deletion: NDN-FLOX-EM1-B6

**Name of Mouse model or mutation:**
**NDN-FLOX-EM1-B6**

**Description:**
Floxed allele (ENSMUSE00000279946) generated using CRISPR/Cas9 genome editing.

**Type of mutation:**
Floxing of exon ENSMUSE00000279946

**Delivery method:**
Pronuclear injection into 1-cell stage embryo.

**Genetic Background:**
C57BL/6J

**Nuclease:**
Cas9 mRNA

**sgRNAs:**

| Protospacer sequence | PAM sequence |
|---|---|
| ATGAATAAGATGCAACATAT | TGG |
| GAGCAGGGCAAAAAGGTAGT | GGG |
| CCTTAATGCATACAGCAGGG | AGG |
| CTATCCTAAAACCTTTATAC | AGG |

**lssDNA donor sequence template**
**Ndn Flox lssDNA**
TCTGGCTTTGGAAGCTTCAACAATGGTCATTGTATAAAGGAGAGTTTGAGGAAATATCAGCTTTAGA
TGACCTCTCGATCTACCTTTTTATTTTTGGACAatccgggggtaccgcgtcgagGCGATCGCATAACTTCGTA
TAGCATACATTATACGAAGTTATAAAATATGAAAAATACACTGTCATAAAATGTGTATTAATAAATAT
GTTGATCATTTTCCACTAGAATCTTAACGGAAGCTTTAGATGAAAACAAAGTGACCTAATAGAAATG
GAGAGCCCAAATCTATAGTTATTAAAATAATTCAAAACCAACAAACTGAGCATCCAATGATGAATGA
ATGCATATATATATATATTTTATATATCCCTCATTTTCATGTGGGGCCTGGGGGATTTAAATTCTGGTTCT
CATGCTTGAACTGCAAAAGCTCTTAACCACTGAACCAAGTCTCATTTTAATTAAAATTTTAAACTCCCA
TCAATTATGCTATATGAATCTACTACTTCTAGAACATTCAAATCTCACATGGATTTATCTCCAGTGTCT
GTAACTACACTTTACATTCTTCCTTGTTTGATCTGCATTCAGATATGTAAGAATTTATCTCTATTAATAA
TTTCTCAATATTCCAATCTTAAACAACTCATCATCATCATAAGGTACAGCTTTCCAAAAGAAAAAAGA
AAAAAAAGTCATTTCTGTTTCTCTTATTCTTTTGTAGAAAAACCAAAATCAAGAATTAAGTCTTTCTCC
AGGACCTTCACATTTAATTTGATTTTGCACAAACTCAGTGTGTCCGAACTTAACTTCACCATCTCAGCT
ACATCTTTCTCCTTCAACTTGTTTCTTTCCCTACCATCATCTAGTTCTGTGCCATACAGGAGACCAGGA
AATCTTTTACATAAGCCTAGTGGTACCCTCCCTTAGACCCCAGTGGTTGGGCTTTGCTGCTTCGCTCCT
TTCCAGCCCTACCACCCTTCTGGCTTCCCAACACGCATGCGCAATATCGCATCAGCCCCGCCCGCCCG
CTGCTGCGGAAGGCGCAGTGCTCAGTAAAGCGCACTTCCTCTGCTGGTCTCCACCGAGGGAGTGCC
CGCTCCAAGAGCTCCAAGCCGCATCGGTCCTGCTCTGATCCGAAGGCGCAGACATGTCGGAACAAA

GTAAGGACCTGAGCGACCCTAACTTTGCAGCCGAGGTCCCCGACTGTGAGATGCAGGACAGCGATG
CCGTTCCGGTGGGGATCCCTCCTCCCGCTTCTCTGGCCGCTAACCTCGCAGGGCCACCGTGCGCTCCC
GAAGGCCCTATGGCAGCCCAACAGGCCTCGCCACCGCCCGAAGAACGGATAGAAGATGTTGACCCT
AAAATCCTGCAGCAGGCCGCAGAGGAGGGCCGCGCCCACCAGCCCCAGAGTCCAGCCCGGCCGAT
CCCAGCACCGCCAGCCCCTGCCCAGCTGGTGCAGAAGGCGCACGAGCTCATGTGGTACGTGTTGGT
GAAGGACCAGAAGAGGATGGTCCTCTGGTTTCCAGACATGGTGAAAGAGGTCATGGGCAGCTACA
AGAAATGGTGCAGAAGCATCCTCAGGCGCACCAGCGTCATCCTCGCCAGAGTGTTCGGGCTGCACC
TGAGGCTGACCAATCTCCACACCATGGAGTTTGCCCTGGTCAAAGCCCTCAGCCCAGAGGAGCTAGA
CAGGGTGGCGCTCAACAACCGTATGCCCATGACAGGCCTCCTGCTCATGATCCTGAGCCTCATCTAT
GTGAAGGGCCGCGGGGCCAGAGAGGGTGCGGTCTGGAATGTGCTGCGCATCCTGGGGCTGAGGC
CCTGGAAGAAGCACTCCACCTTCGGAGACGTGAGGAAGATAATCACCGAGGAGTTCGTCCAGCAGA
ATTACCTGAAGTACCAGCGTGTGCCCCACATCGAGCCTCCCGAGTACGAGTTCTTCTGGGGGTCCAG
AGCTAACCGTGAAATCACCAAGATGCAGATCATGGAGTTCCTGGCCAGAGTCTTCAAGAAAGATCCC
CAGGCGTGGCCTTCCCGATACAGGGAGGCTCTGGAGCAGGCCAGAGCTCTGCGGGAGGCTAATCTT
GCTGCCCAGGCCCCCCGCAGCAGTGTCTCTGAGGACTAAAAAGGTCCAGGGGCACACTGATAGTTT
CTGACCCATACTAGGGCTGTGTAAGGGTGGGGTTGAGTCATTAGAGTATCCCAAATCCACAGTGCA
GTATTTCATGTATAATTTTTAAGTTTTCCATACAGTGCTTTTGTACCTTGTAATGCTATTCATTTGTGTA
CTCGTGTAGTGTTTAAGATTGATGCATGTGTGATAAGTATTTGGTACTTTCACTTTTGTGCTTTCGTGC
ATTTTTGTACAAGAGATGTGCTGTGCTAAACTTGTGAAATACATTGAGGTGTTCTGTATCTTGTTCTTT
GTATGGGACTGATGATCTGTATCGACAAAGAAGGCCCTGGAGAGTTAGCAGGACTTAACAGCAACG
CAGACCTGAGCAAGAGAAAGGTCAAGGCCTTTCTCCATATGACTTCAACTGGCACAGGAAGCATCC
ATGTGGAATGGACTGATTTGAACTGGACTGTTCTCAGTGTAGGCACTTAGCACCCTTTACAAAACAT
GTATGCAACCCCACCATAAATAAACGTTAAAATGAGCATTAAAGATACTGTGAAATAATTTCTTGGG
GGGGGGGAGGAGGCATTTATTTTTGATGAAAGGGGGAGTGGGAATGGGTGGGGATGAGAAAGAT
TCTCCCTGTGTGCAGCATAAGCAAACACGGTGTCCTAACACAGTGGGAGACCACACGACCACTCAGA
AGAAACTTGCAGTGCTTTAATCAATTTGTCACATAAAAGGTATAACAACAACAACAAAAAAGCTCAT
TATCACATAACATGGGGATAATAACAGGTATATGGAGGGCCTGCCCCTTTTCTGTTGCATTTTTTAAA
ATAGTCTTTGATGGACACTTTAGCAGAATAAACCCTTATCATGCATTTTCTTCTTTATTTTGTCCTCCCA
CCAGCCATAAGAAGTAAGGAATAACTGTATCATCACAGAGCAAGAAAAAATAAAAGGGGTGTTGGT
AATGAACTATCCTAAAATAACTTCGTATAGCATACATTATACGAAGTTATCGCCGGCGggtctgagctcgc
catcagtTGCATTAAGGGTTTGATTTTTCTAATTAATTCCAAGAGTTTCACAGAAAATTAGCTTCTGAAT
GTACAAATTAGGAAACTCGTTTCAGAAAGAATTGATG

**Pronuclear Microinjection mixes:**
Microinjection buffer (MIB; 10 mM Tris–HCl, 0.1 mM EDTA, 100 mM NaCl, pH7.5) was prepared and filtered through a 2 nm filter and autoclaved. Cas9 mRNA, sgRNAs and ssODN were diluted and mixed in MIB to the working concentrations of 100 ng/µl, 50 ng/µl each and 50 ng/µl, respectively. Injected embryos were re-implanted in CD1 pseudo-pregnant females. Host females were allowed to litter and rear $F_0$ progeny.

**Sequence details**
**WT**
ATAACACGTCTAGGCTGGAGTTTAAAACAAGTTATTGTACATGTGCAAATATTCTCAAATCTGGAAA
ACTCTGAATCGCTTCTTGCCATAACTATTCTGGGTAAAAGATAGCTGTTATATCTAAACAAACATATT
TTAGAACCTAGGAATGCCAACATGATATCTTAAATCTGTCCGCTTTAAGTGATGAGAAGGGCTAAGG
AATTCCATTATTTCAGAACAAGAAAGATGATCAAATCTAAGGACTTCTGGCTTTGGAAGCTTCAACA
ATGGTCATTGTATAAAGGAGAGTTTGAGGAAATATCAGCTTTAGATGACCTCTCGATCTACCTTTTTA

TTTTTGGACACCAATATGTTGCATCTTATTCATGAGAGACTGTTAGGTATCGAAAAGAGCTGCATTTA
AAATATATTGGGAAAGATTTGGATGTGCTCCCAACATGGAATTAGTCATTTGTGAATGACATAATTT
AAGAAGTAGAAGATTTCTTATAAACGCAGAAGATGCAGTGAGCAGGGCAAAAAGGTAGTGGGAAA
ATATGAAAAATACACTGTCATAAAATGTGTATTAATAAATATGTTGATCATTTTCCACTAGAATCTTA
ACGGAAGCTTTAGATGAAAACAAAGTGACCTAATAGAAATGGAGAGCCCAAATCTATAGTTATTAA
AATAATTCAAAACCAACAAACTGAGCATCCAATGATGAATGAATGCATATATATATATTTTATATATC
CCTCATTTTCATGTGGGGCCTGGGGGATTTAAATTCTGGTTCTCATGCTTGAACTGCAAAAGCTCTTA
ACCACTGAACCAAGTCTCATTTTAATTAAAATTTTAAACTCCCATCAATTATGCTATATGAATCTACTA
CTTCTAGAACATTCAAATCTCACATGGATTTATCTCCAGTGTCTGTAACTACACTTTACATTCTTCCTTG
TTTGATCTGCATTCAGATATGTAAGAATTTATCTCTATTAATAATTTCTCAATATTCCAATCTTAAACA
ACTCATCATCATCATAAGGTACAGCTTTCCAAAAGAAAAAAGAAAAAAAAGTCATTTCTGTTTCTCTT
ATTCTTTTGTAGAAAAACCAAAATCAAGAATTAAGTCTTTCTCCAGGACCTTCACATTTAATTTGATTT
TGCACAAACTCAGTGTGTCCGAACTTAACTTCACCATCTCAGCTACATCTTTCTCCTTCAACTTGTTTCT
TTCCCTACCATCATCTAGTTCTGTGCCATACAGGAGACCAGGAAATCTTTTACATAAGCCTAGTGGTA
CCCTCCCTTAGACCCCAGTGGTTGGGCTTTGCTGCTTCGCTCCTTTCCAGCCCTACCACCCTTCTGGCT
TCCCAACACGCATGCGCAATATCGCATCAGCCCCGCCCGCCCGCTGCTGCGGAAGGCGCAGTGCTCA
GTAAAGCGCACTTCCTCTGCTGGTCTCCACCGAGGGAGTGCCCGCTCCAAGAGCTCCAAGCCGCATC
GGTCCTGCTCTGATCCGAAGGCGCAGACATGTCGGAACAAAGTAAGGACCTGAGCGACCCTAACTT
TGCAGCCGAGGTCCCCGACTGTGAGATGCAGGACAGCGATGCCGTTCCGGTGGGGATCCCTCCTCC
CGCTTCTCTGGCCGCTAACCTCGCAGGGCCACCGTGCGCTCCCGAAGGCCCTATGGCAGCCCAACAG
GCCTCGCCACCGCCCGAAGAACGGATAGAAGATGTTGACCCTAAAATCCTGCAGCAGGCCGCAGAG
GAGGGCCGCGCCCACCAGCCCCAGAGTCCAGCCCGGCCGATCCCAGCACCGCCAGCCCCTGCCCAG
CTGGTGCAGAAGGCGCACGAGCTCATGTGGTACGTGTTGGTGAAGGACCAGAAGAGGATGGTCCT
CTGGTTTCCAGACATGGTGAAAGAGGTCATGGGCAGCTACAAGAAATGGTGCAGAAGCATCCTCAG
GCGCACCAGCGTCATCCTCGCCAGAGTGTTCGGGCTGCACCTGAGGCTGACCAATCTCCACACCATG
GAGTTTGCCCTGGTCAAAGCCCTCAGCCCAGAGGAGCTAGACAGGGTGGCGCTCAACAACCGTATG
CCCATGACAGGCCTCCTGCTCATGATCCTGAGCCTCATCTATGTGAAGGGCCGCGGGGCCAGAGAG
GGTGCGGTCTGGAATGTGCTGCGCATCCTGGGGCTGAGGCCCTGGAAGAAGCACTCCACCTTCGGA
GACGTGAGGAAGATAATCACCGAGGAGTTCGTCCAGCAGAATTACCTGAAGTACCAGCGTGTGCCC
CACATCGAGCCTCCCGAGTACGAGTTCTTCTGGGGGTCCAGAGCTAACCGTGAAATCACCAAGATGC
AGATCATGGAGTTCCTGGCCAGAGTCTTCAAGAAAGATCCCCAGGCGTGGCCTTCCCGATACAGGG
AGGCTCTGGAGCAGGCCAGAGCTCTGCGGGAGGCTAATCTTGCTGCCCAGGCCCCCCGCAGCAGTG
TCTCTGAGGACTAAAAAGGTCCAGGGGCACACTGATAGTTTCTGACCCATACTAGGGCTGTGTAAG
GGTGGGGTTGAGTCATTAGAGTATCCCAAATCCACAGTGCAGTATTTCATGTATAATTTTTAAGTTTT
CCATACAGTGCTTTTGTACCTTGTAATGCTATTCATTTGTGTACTCGTGTAGTGTTTAAGATTGATGCA
TGTGTGATAAGTATTTGGTACTTTCACTTTTGTGCTTTCGTGCATTTTTGTACAAGAGATGTGCTGTGC
TAAACTTGTGAAATACATTGAGGTGTTCTGTATCTTGTTCTTTGTATGGGACTGATGATCTGTATCGA
CAAAGAAGGCCCTGGAGAGTTAGCAGGACTTAACAGCAACGCAGACCTGAGCAAGAGAAAGGTCA
AGGCCTTTCTCCATATGACTTCAACTGGCACAGGAAGCATCCATGTGGAATGGACTGATTTGAACTG
GACTGTTCTCAGTGTAGGCACTTAGCACCCTTTACAAAACATGTATGCAACCCCACCATAAATAAACG
TTAAAATGAGCATTAAAGATACTGTGAAATAATTTCTTGGGGGGGGGGAGGAGGCATTTATTTTTG
ATGAAAGGGGGAGTGGGAATGGGTGGGGATGAGAAAGATTCTCCCTGTGTGCAGCATAAGCAAAC
ACGGTGTCCTAACACAGTGGGAGACCACACGACCACTCAGAAGAAACTTGCAGTGCTTTAATCAATT
TGTCACATAAAAGGTATAACAACAACAACAAAAAAGCTCATTATCACATAACATGGGGATAATAACA
GGTATATGGAGGGCCTGCCCCTTTTCTGTTGCATTTTTTAAAATAGTCTTTGATGGACACTTTAGCAG
AATAAACCCTTATCATGCATTTTCTTCTTTATTTTGTCCTCCCACCAGCCATAAGAAGTAAGGAATAAC
TGTATCATCACAGAGCAAGAAAAAATAAAAGGGGTGTTGGTAATGAACTATCCTAAAACCTTTATAC

AGGAGTCTTGACACCAAGCTCCCCATGGCCACCACTCACACACGCACTCTGGCCTCCCTGCTGTATGC
ATTAAGGGTTTGATTTTTCTAATTAATTCCAAGAGTTTCACAGAAAATTAGCTTCTGAATGTACAAAT
TAGGAAACTCGTTTCAGAAAGAATTGATGACCTAATCAAGTCCTTATCTTGCTAAGGGGTAAAAACA
AGATCTGAAATTTCATCTTTCCAGAAGGATGAAGATGAGCGAAACTATTCTGACAGTCGTACCAGTA
AGAGCCATGGCACTAATGAAATAGAGGAGGGGTTGGGATTAAGGAGCTGCCCCCAGAAGCATAGT
CACATTTGCCACTCAGCAGTCAGTATGTTGCAGTGGGCAGTAGGTGAAGAGAGGGTGAAATGGGAT
GAGAATCACTGTAAACAGCAACAACTGGGGTAAAAATCTCCTTAGAAAGCTTTTGGTCTGGGGAGA
AAGAGAGACACGAGGCTTTAATGAGATTAGAAATCCTTTAGAATTTGTGGTAAAAGTCATTTAGTAA
AAGCTAAAAAGTAATTTCCCCCCTGGGAAAGGTGAATAAGGGGAGATACCTGTCACACCAT

**NDN-FLOX-EM1-B6**
ATAACACGTCTAGGCTGGAGTTTAAAACAAGTTATTGTACATGTGCAAATATTCTCAAATCTGGAAA
ACTCTGAATCGCTTCTTGCCATAACTATTCTGGGTAAAAGATAGCTGTTATATCTAAACAAACATATT
TTAGAACCTAGGAATGCCAACATGATATCTTAAATCTGTCCGCTTTAAGTGATGAGAAGGGCTAAGG
AATTCCATTATTTCAGAACAAGAAAGATGATCAAATCTAAGGACTTCTGGCTTTGGAAGCTTCAACA
ATGGTCATTGTATAAAGGAGAGTTTGAGGAAATATCAGCTTTAGATGACCTCTCGATCTACCTTTTTA
TTTTTGGACAatccgggggtaccgcgtcgagGCGATCGC*ATAACTTCGTATAGCATACATTATACGAAGTTA*
*T*AAAATATGAAAAATACACTGTCATAAAATGTGTATTAATAAATATGTTGATCATTTTCCACTAGAAT
CTTAACGGAAGCTTTAGATGAAAACAAAGTGACCTAATAGAAATGGAGAGCCCAAATCTATAGTTAT
TAAAATAATTCAAAACCAACAAACTGAGCATCCAATGATGAATGAATGCATATATATATATTTTATAT
ATCCCTCATTTTCATGTGGGGCCTGGGGGATTTAAATTCTGGTTCTCATGCTTGAACTGCAAAAGCTC
TTAACCACTGAACCAAGTCTCATTTTAATTAAAATTTTAAACTCCCATCAATTATGCTATATGAATCTA
CTACTTCTAGAACATTCAAATCTCACATGGATTTATCTCCAGTGTCTGTAACTACACTTTACATTCTTCC
TTGTTTGATCTGCATTCAGATATGTAAGAATTTATCTCTATTAATAATTTCTCAATATTCCAATCTTAAA
CAACTCATCATCATCATAAGGTACAGCTTTCCAAAAGAAAAAAGAAAAAAAAGTCATTTCTGTTTCTC
TTATTCTTTTGTAGAAAAACCAAAATCAAGAATTAAGTCTTTCTCCAGGACCTTCACATTTAATTTGAT
TTTGCACAAACTCAGTGTGTCCGAACTTAACTTCACCATCTCAGCTACATCTTTCTCCTTCAACTTGTTT
CTTTCCCTACCATCATCTAGTTCTGTGCCATACAGGAGACCAGGAAATCTTTTACATAAGCCTAGTGG
TACCCTCCCTTAGACCCCAGTGGTTGGGCTTTGCTGCTTCGCTCCTTTCCAGCCCTACCACCCTTCTGG
CTTCCCAACACGCATGCGCAATATCGCATCAGCCCCGCCCGCCCGCTGCTGCGGAAGGCGCAGTGCT
CAGTAAAGCGCACTTCCTCTGCTGGTCTCCACCGAGGGAGTGCCCGCTCCAAGAGCTCCAAGCCGCA
TCGGTCCTGCTCTGATCCGAAGGCGCAGACATGTCGGAACAAAGTAAGGACCTGAGCGACCCTAAC
TTTGCAGCCGAGGTCCCCGACTGTGAGATGCAGGACAGCGATGCCGTTCCGGTGGGGATCCCTCCT
CCCGCTTCTCTGGCCGCTAACCTCGCAGGGCCACCGTGCGCTCCCGAAGGCCCTATGGCAGCCCAAC
AGGCCTCGCCACCGCCCGAAGAACGGATAGAAGATGTTGACCCTAAAATCCTGCAGCAGGCCGCAG
AGGAGGGCCGCGCCCACCAGCCCCAGAGTCCAGCCCGGCCGATCCCAGCACCGCCAGCCCCTGCCC
AGCTGGTGCAGAAGGCGCACGAGCTCATGTGGTACGTGTTGGTGAAGGACCAGAAGAGGATGGTC
CTCTGGTTTCCAGACATGGTGAAAGAGGTCATGGGCAGCTACAAGAAATGGTGCAGAAGCATCCTC
AGGCGCACCAGCGTCATCCTCGCCAGAGTGTTCGGGCTGCACCTGAGGCTGACCAATCTCCACACCA
TGGAGTTTGCCCTGGTCAAAGCCCTCAGCCCAGAGGAGCTAGACAGGGTGGCGCTCAACAACCGTA
TGCCCATGACAGGCCTCCTGCTCATGATCCTGAGCCTCATCTATGTGAAGGGCCGCGGGGCCAGAG
AGGGTGCGGTCTGGAATGTGCTGCGCATCCTGGGGCTGAGGCCCTGGAAGAAGCACTCCACCTTCG
GAGACGTGAGGAAGATAATCACCGAGGAGTTCGTCCAGCAGAATTACCTGAAGTACCAGCGTGTGC
CCCACATCGAGCCTCCCGAGTACGAGTTCTTCTGGGGGTCCAGAGCTAACCGTGAAATCACCAAGAT
GCAGATCATGGAGTTCCTGGCCAGAGTCTTCAAGAAAGATCCCCAGGCGTGGCCTTCCCGATACAG
GGAGGCTCTGGAGCAGGCCAGAGCTCTGCGGGAGGCTAATCTTGCTGCCCAGGCCCCCCGCAGCA
GTGTCTCTGAGGACTAAAAAGGTCCAGGGGCACACTGATAGTTTCTGACCCATACTAGGGCTGTGTA

AGGGTGGGGTTGAGTCATTAGAGTATCCCAAATCCACAGTGCAGTATTTCATGTATAATTTTTAAGTT
TTCCATACAGTGCTTTTGTACCTTGTAATGCTATTCATTTGTGTACTCGTGTAGTGTTTAAGATTGATG
CATGTGTGATAAGTATTTGGTACTTTCACTTTTGTGCTTTCGTGCATTTTTGTACAAGAGATGTGCTGT
GCTAAACTTGTGAAATACATTGAGGTGTTCTGTATCTTGTTCTTTGTATGGGACTGATGATCTGTATC
GACAAAGAAGGCCCTGGAGAGTTAGCAGGACTTAACAGCAACGCAGACCTGAGCAAGAGAAAGGT
CAAGGCCTTTCTCCATATGACTTCAACTGGCACAGGAAGCATCCATGTGGAATGGACTGATTTGAAC
TGGACTGTTCTCAGTGTAGGCACTTAGCACCCTTTACAAAACATGTATGCAACCCCACCATAAATAAA
CGTTAAAATGAGCATTAAAGATACTGTGAAATAATTTCTTGGGGGGGGGGAGGAGGCATTTATTTTT
GATGAAAGGGGGAGTGGGAATGGGTGGGGATGAGAAAGATTCTCCCTGTGTGCAGCATAAGCAAA
CACGGTGTCCTAACACAGTGGGAGACCACACGACCACTCAGAAGAAACTTGCAGTGCTTTAATCAAT
TTGTCACATAAAAGGTATAACAACAACAACAAAAAAGCTCATTATCACATAACATGGGGATAATAAC
AGGTATATGGAGGGCCTGCCCCTTTTCTGTTGCATTTTTTAAAATAGTCTTTGATGGACACTTTAGCA
GAATAAACCCTTATCATGCATTTTCTTCTTTATTTTGTCCTCCCACCAGCCATAAGAAGTAAGGAATAA
CTGTATCATCACAGAGCAAGAAAAAATAAAAGGGGTGTTGGTAATGAACTATCCTAAA*ATAACTTCG*
*TATAGCATACATTATACGAAGTTAT*CGCCGGCGggtctgagctcgccatcagtTGCATTAAGGGTTTGATTT
TTCTAATTAATTCCAAGAGTTTCACAGAAAATTAGCTTCTGAATGTACAAATTAGGAAACTCGTTTCA
GAAAGAATTGATGACCTAATCAAGTCCTTATCTTGCTAAGGGGTAAAAACAAGATCTGAAATTTCAT
CTTTCCAGAAGGATGAAGATGAGCGAAACTATTCTGACAGTCGTACCAGTAAGAGCCATGGCACTA
ATGAAATAGAGGAGGGGTTGGGATTAAGGAGCTGCCCCCAGAAGCATAGTCACATTTGCCACTCAG
CAGTCAGTATGTTGCAGTGGGCAGTAGGTGAAGAGAGGGTGAAATGGGATGAGAATCACTGTAAA
CAGCAACAACTGGGGTAAAAATCTCCTTAGAAAGCTTTTGGTCTGGGGAGAAAGAGAGACACGAG
GCTTTAATGAGATTAGAAATCCTTTAGAATTTGTGGTAAAAGTCATTTAGTAAAAGCTAAAAAGTAA
TTTCCCCCCTGGGAAAGGTGAATAAGGGGAGATACCTGTCACACCAT

**Primers used to QC the allele: All amplicons were amplified and sequence verified in both directions.**
*Please note, the size of the allele warranted sequencing of multiple overlapping PCR products to get complete coverage.

| | |
|---|---|
| Geno_Ndn_F3 primer (5’-3’) | ATAACACGTCTAGGCTGGAGTT |
| Geno_Ndn_R3 primer (5’-3’) | AAGTGTAGTTACAGACACTGGAG |
| Annealing Temperature (°C) | 59 |
| Elongation time (min) | 1 |
| WT product size (bp) | 932 |
| Mutant product size (bp) | 808 |
| Sequence with primers (if different to amplification primers) | |
| Notes | |

| | |
|---|---|
| Geno_Ndn_F4 primer (5’-3’) | CCACACGACCACTCAGAAGAA |
| Geno_Ndn_R4 primer (5’-3’) | ATGGTGTGACAGGTATCTCCC |
| Annealing Temperature (°C) | 59 |
| Elongation time (min) | 1 |
| WT product size (bp) | 909 |
| Mutant product size (bp) | 896 |
| Sequence with primers (if different to amplification primers) | |
| Notes | |

| | |
|---|---|
| Geno_Ndn_F1 primer (5’-3’) | CGCTTTAAGTGATGAGAAGGGC |
| Geno_Ndn_R2 primer (5’-3’) | TGGTGTGGAGATTGGTCAGC |
| Annealing Temperature (°C) | 59 |
| Elongation time (min) | 3.5 |
| WT product size (bp) | 1845 |
| Mutant product size (bp) | 1721 |
| Sequence with primers (if different to amplification primers) | |
| Notes | |

| | |
|---|---|
| Geno_Ndn_F2 primer (5’-3’) | CGTGTTGGTGAAGGACCAGA |
| Geno_Ndn_R1 primer (5’-3’) | ACTGACTGCTGAGTGGCAAA |
| Annealing Temperature (°C) | 59 |
| Elongation time (min) | 2.5 |
| WT product size (bp) | 1997 |
| Mutant product size (bp) | 1984 |
| Sequence with primers (if different to amplification primers) | Also sequenced with:<br>Geno_Ndn_F5 primer<br>(CACTTTTGTGCTTTCGTGCATT)<br>Geno_Ndn_R5 primer<br>(TTCACAAGTTTAGCACAGCACA) |
| Notes | |

| | |
|---|---|
| Geno_Ndn_F2 primer (5’-3’) | CGTGTTGGTGAAGGACCAGA |
| Geno_Ndn_R4 primer (5’-3’) | ATGGTGTGACAGGTATCTCCC |
| Annealing Temperature (°C) | 59 |
| Elongation time (min) | 2.5 |
| WT product size (bp) | 2233 |
| Mutant product size (bp) | 2220 |
| Sequence with primers (if different to amplification primers) | Also sequenced with:<br>Geno_Ndn_F5 primer<br>(CACTTTTGTGCTTTCGTGCATT)<br>Geno_Ndn_R5 primer<br>(TTCACAAGTTTAGCACAGCACA)<br>Geno_Ndn_R1 primer<br>(ACTGACTGCTGAGTGGCAAA) |
| Notes | |

| | |
|---|---|
| Geno_Ndn_F2 primer (5’-3’) | CGTGTTGGTGAAGGACCAGA |
| Geno_Ndn_R1 primer (5’-3’) | ACTGACTGCTGAGTGGCAAA |
| Annealing Temperature (°C) | 59 |
| Elongation time (min) | 2.5 |
| WT product size (bp) | 1997 |
| Mutant product size (bp) | 1984 |
| Sequence with primers (if different to amplification primers) | Also sequenced with:<br>LoxPR primer<br>(ACTGATGGCGAGCTCAGACC)<br>Geno_Ndn_F4 primer<br>(CCACACGACCACTCAGAAGAA) |
| Notes | |

| | |
|---|---|
| Geno_Ndn_F6 primer (5'-3') | ACGGAAGCTTTAGATGAAAACAAAG |
| Geno_Ndn_R6 primer (5'-3') | TCTTGTAGCTGCCCATGACC |
| Annealing Temperature (°C) | 61 |
| Elongation time (min) | 1.5 |
| WT product size (bp) | 1328 |
| Mutant product size (bp) | 1328 |
| Sequence with primers (if different to amplification primers) | Also sequenced with:<br>Geno_Ndn_R7 primer<br>(GGAAAGCTGTACCTTATGATGATGATGAG) |
| Notes | |

| | |
|---|---|
| LoxPF primer (5'-3') | atccgggggtaccgcgtcgag |
| Geno_Ndn_R6 primer (5'-3') | TCTTGTAGCTGCCCATGACC |
| Annealing Temperature (°C) | 63 |
| Elongation time (min) | 1.5 |
| WT product size (bp) | NA |
| Mutant product size (bp) | 1462 |
| Sequence with primers (if different to amplification primers) | Also sequenced with:<br>Geno_Ndn_R7 primer<br>(GGAAAGCTGTACCTTATGATGATGATGAG) |
| Notes | |

Copy counting of the donor sequence was carried out by ddPCR at the F1 stage to confirm donor oligos were inserted once on target into the genome. The following Taqman assay was used to copy count the donor sequence compared against a VIC-labelled reference assay for Dot1l:

| | |
|---|---|
| Assay name | Ndn-CR-LOA |
| Forward Primer (5'-3') | GCACCTGAGGCTGACCAAT |
| Reverse Primer (5'-3') | TGGGCTGAGGGCTTTGAC |
| Probe (5'-3') | TCCACACCATGGAGTTTGCCCTG |
| Label | FAM-BHQ1 |

The ddPCR assay recognises the critical region of the Ndn gene. Therefore, WT controls and F1 animals with evidence of the Floxed exon will call at 2 copies and heterozygous deletion mutants are expected to call at 1 copy.

| | |
|---|---|
| Assay name | Ndn-FLOX- 5'-MUT1 |
| Forward Primer (5'-3') | ATCAGCTTTAGATGACCTCTCGA |
| Reverse Primer (5'-3') | TAAAGCTTCCGTTAAGATTCTAGTGG |
| Probe (5'-3') | TCGAGGCGATCGCATAACTTCG |
| Label | FAM-BHQ1 |

The ddPCR assay recognises the 5' LoxP site specific to the Ndn FLOX modification. Therefore, WT controls are expected to call at 0 copies and F1 animals with evidence of the Floxed exon will call at ~1 copy.

| | |
|---|---|
| Assay name | Ndn-FLOX-3’-MUT1 |
| Forward Primer (5’-3’) | GGGTGTTGGTAATGAACTATCCTAAA |
| Reverse Primer (5’-3’) | GAAGCTAATTTTCTGTGAAACTCTTGGA |
| Probe (5’-3’) | AAGTTATCGCCGGCGGGTCTGA |
| Label | FAM-BHQ1 |

The ddPCR assay recognises the 3' LoxP site specific to the Ndn FLOX modification. Therefore, WT controls are expected to call at 0 copies and F1 animals with evidence of the Floxed exon will call at ~1 copy.

| | |
|---|---|
| Reference Assay Name | Dot1l |
| Forward primer (5’-3’) | GCCCCAGCACGACCATT |
| Reverse primer (5’-3’) | TAGTTGGCATCCTTATGCTTCATC |
| Probe (5’-3’) | CCCAACAGGCCTGGATTCTCAATGC |
| Label | VIC |

VIC-labelled reference assay for Dot1l gene.

MRC Harwell

### *Magel2* deletion: MAGEL2-FLOX-EM1-B6

**Name of Mouse model or mutation:**
**MAGEL2-FLOX-EM1-B6**

**Description:**
Floxed allele made by CRISPR/Cas9 gene editing.

**Type of mutation:**
Floxing of exon ENSMUSE00000497291. Note that Magel2 is a single exon gene.

**Delivery method:**
Pronuclear injection into 1-cell stage embryo.

**Genetic Background:**
C57BL/6J

**Nuclease:**
Cas9 mRNA

**sgRNAs:**

| Protospacer sequence | PAM sequence |
|---|---|
| GGGAAAATCAACTTCTAAAG | TGG |
| GTAAGACGCGGCAAGCTAAG | AGG |

**lssDNA donor sequence template**

**Magel2 Flox 5' lssDNA**
TTCCAGAAAACTTTTCTTATACTCTCCATTTTTCTAGAACTAATTTCCAGCACATTGCTTCATTTTTT
AATTATTGTTATTTTCAGGATGTCTTTCACACTCTATTGTCTCATCTCACatccgggggtaccgcgtcgag
GCGATCGCATAACTTCGTATAGCATACATTATACGAAGTTATTGAATTTCTGTTAATTTACTTCCAA
GGTTCCTTTTCTTTCCTCTCTAATTGGCTCGTCTTTTCTCTACCCGATTCTACGATGTAGTCAGTAT
TCTTCTAGATGAGTTTAGATTTTCCCC

**Magel2 Flox 3' lssDNA**
GATAGTAAAACCTGTGGATGCTGACAGCTTCCCAGAAGAGTACTCCATTCATCCTAAAAGTGATG
CAGTGATGGTGGGTGGTGAAGCCAGAGATCTCCTTAAAGTGAAAGTAAAACTTCATAACTTCGTA
TAGCATACATTATACGAAGTTATCGCCGGCGggtctgagctcgccatcagtGAGTCCCCTCTGATTTAAT
TTTTACAATAACTGTTCAAGTTAGACATTGTATCTAGCTTGTAGGTGAGGCTATAAGACACAAACA
GGTTAATTAAAATACATATATGGTGGCATGCTGG

**Pronuclear Microinjection mixes:**
Microinjection buffer (MIB; 10 mM Tris–HCl, 0.1 mM EDTA, 100 mM NaCl, pH7.5) was prepared and filtered through a 2 nm filter and autoclaved. Cas9 mRNA, sgRNAs and ssODNs were diluted and mixed in MIB to the working concentrations of 100 ng/µl, 50 ng/µl each and 50 ng/µl, respectively. Injected embryos were re-implanted in CD1 pseudo-pregnant females. Host females were allowed to litter and rear $F_0$ progeny.

**Sequence details**
**MAGEL2 WT**

AAACGGTCAGTCATCCTGGGGCATATTTCTTCTTTATGGCCCACCAGTTAAAAACGAACAACAACAA
CAAAACTATTTAGGAGCATGTGCTGAAGGTATGTTAAGAGTTGGGTACAGCCTTCAACATTCTCTTCT
GCCCCCTAGTGGTTTCTTTCCTCACCTTGGACGACTGCCCTAGGCTGCAAACCACTCCTTGTAAAGGA
ATCATAATGCTAGTTACCGTCTATAAACCACTAAACTTGACCTAAACTGAGATTGCCACTTAATTTTG
GGGAGAAAGAATACCAATATGCAGTAATTTTGATGGGTACCTCTTTATTTGTTAACATGGTATATGTC
CATTTTTTAGCTGATTTTCCAGAAAACTTTTCTTATACTCTCCATTTTTCTAGAACTAATTTCCAGCACA
TTGCTTCATTTTTTAATTATTGTTATTTTCAGGATGTCTTTCACACTCTATTGTCTCATCTCACCCACTTT
AGAAGTTGATTTTCCCGCGTTTAAACTTCTGCCTCCGTGAATTTCTGTTAATTTACTTCCAAGGTTCCT
TTTCTTTCCTCTCTAATTGGCTCGTCTTTTCTCTACCCGATTCTACGATGTAGTCAGTATTCTTCTAGAT
GAGTTTAGATTTTCCCCCCTCTGAACACACTTTTAAACAATTGTATCCACTACTACTACTATGTTTTGA
TAAAGAATGGGCAAACTTGATTCTCGGATCCTCGTAATGAATACAGTTATAATAGACCACCTATGTTA
TCCCTGGGTTGACTGACTCATGTCCTAGCAAATATTCATCAACTCCCCTTACCCTGCATGTGCTTCCTG
CCCTTCAGTTACTTGACAGAATAAACTACTCCAGGAAATTCACACACCCGAGACAGATACCTGAATA
CAGTCATTTTTTTCTACACCTTCGTTAAATTGTAATAGCTGTCTCGAATAGCTCCCCAGTCCCTCTTAC
GTCAATGTACTTCGTTTCTGAGCTCAGACCACACCACTACCCCCACCTTTGGACATGTTCCATGTTGCT
AGTGGTTTATTCCCACTTCATAATGCAGCACAGTGATTTTTTTTGAACCGTAAATCTGTCCCTATCACT
TGCCTATATAAACTCTTGCTTAAAGAGTATGAACTAGCTTAGTTGAGGATCTATACCGCAGGAGAAG
GGATATTGAGTCAGTCGGGACTTTGGTAGACGCCCCTCTGAACAATCCACTTGTGGTGGTCGGGGG
CGGGGAAGGGGCGGGATTGAGCCCCCTCCCCCCTCCTTCTTACTGCGGAGAGCTGCAGAGCCAGCC
GACACTCTGTGTGAGCTGCGCTGCGAGCTGTGAGCTGAGAGCTGTGAGCTGCGTGGCGGAGCAAG
CCAGGCAGTAGCACTTGGCTGAAAGCTGCGGTGCTAGCCAGGCAGCGCTCGCTGAGAGTTGCGGT
GCCAGCCAGGTAGTGCTTGCTGAGAGCTGCTGAGAGAGCCCAGCTGCGCTCGCTGAGAGCTGCGG
AGAGAACCGCGGAACACGCCAGTCTGCACCGGCTTCCTATTCAGCCTTCAAGCTGGAGGGGGATCC
TGGACAGCCACGTCGGCATGCTCAGCACTGGAAGAAAAAGCGCAGCAACCGGAGACGGGCCACCA
CCAGTCAAGCAAGGGAC**ATGTCGCAGCTAAGTACGAATCTGGGGGATTCCAGCCCTCCGGAGTCC**
**CCAGTGCCTGCAGTCCATAGCCGCCCTACGGTTCTGATGCGGGCTCCGCCTGCTTCCTCCCGGGCTC**
**CTCCTGTCCCCTGGGATCCACCTCCAGTGGACTTGCAGGCTCCCATGGCTGCTTGGCAGGCTCCTCA**
**ACCTGCCTGGGAGGCTCCGGAGGGCCAGCTGCCTGCCCCAGTGGCTCAGCTGGCCCAGCCTCCTG**
**GTCTAGGGGCCCCAATGGTCCAGGCTCCACCGCTCGGAGGGGGGATGGCCAAGCCTCCAACTCCT**
**GGAGTCTTGATGGTCCATCAGCCCCCTCCGGGAGCCCCCATGGCCCAGTCTTCAACTCCGGGAGTC**
**CTGATGCTACATCCTTCTGTCACGGGGGCCCCTTTGGCTCATCCTCCTCCCCCAGGAACCCCGATGA**
**CACACCCTCCCGGGACCTCGATGGCGCACCCTCCTCCTCCTCCTCCTCCCCCTCCTCCTCCTCCTCC**
**CTGGGACCCCAATGACCCACCCACCTCCTCCTGGGACTCCGATGGGTCACCATCCTCCTCCTGGGAA**
**CCCTATGACCCATCCTCCACCGGGGAACCCGATGGTGCATCCTCTCACTCATGGAGCCCCGATGGTC**
**CATGGGGGACCACATGGAACTCCAATGCCGCATGTTCCCATTACGGGGACACCGATAGCCCAGCA**
**GCCAACTCCAGGAGTCCTGATGGCCCAGCAGCTGACACCGGGAGTCCTGATGGTCCAGCCGCCTG**
**CTCCGGGAGCTCCGATGGTCCAGCCACCTCCACAGGCTGCCTTGATGACCCAGCCTGCACCTTCGAT**
**TACTCCGATGGCCAAGCCTCCAGGTCCTGGTGTCGTGATGATCCATCCTCCAGGTGCCAGAGGTCC**
**AATCATTCAGACTCCAGTATCAGGAGCACCAATGGCTCAGACAGTGCTGCCCCCTGGGCAGCCTCT**
**GGCCACTTGGGCCCCACAGGGTCAGCCTTTGATCCTACAAATCCAGTCTCAAGTCATAAGGGCTCC**
**TCCACAGGTTCCCTCTGTACCACAAGCCCCCCAGGTACAGCTGGCCACACCCCCAGGCTGGCAAGC**
**CACCACACCCAACTGGCAGGTGACCCCCCAGGGGTGGCCAGCAACGCCTTTGACATGGCAGGCTA**
**CACAGGTGACCTGGCAGGCCCCCACAATAGCCTGGCAGGCCACTCAACCTGGGCGCCAAGGGCAC**
**TCAACCATTCGTACTGGTCACACACCCATTCGACCTGGACCAGCTCCATTGCTTCGCCAGATACCTC**

**CTATGATCCGTCAGATCCAACCTGTGATGAGGCAAGCCCCACCACTGATCCGACAGGTCCCTATCA**
**GACCAGCTCCACATGGCATAGCAAGCCAGCCTCAGCTGTGGCAGGTCCTGCCACCCCCACCTCCAC**
**TGCGGCAGGCTCCACAGGCTCGTCTACTGCTCCCGAGGGTACCAGGAACAGGCCAGGTGTCTACA**
**GTACCACCAGTTGCTCAGATACATTTGGTGCCACAGTCAGGCCCACAGGTGCCCCAGACAGTACTG**
**CCAGCCCAACTGTCTATCCCAATTCCTGTACCCCAGGCTGCTGCTCAGTCTGCTCCCCGGACTGTGC**
**ATTGCCCACCCATCATCTGGCAGGCCCCCAAAGGCCAGGCCCCGGTGCCACAGGAGCTCCCAGTGC**
**CACAGGAGCTCCCGGTGCCACAGGAGCTCCCGGTGCCACAAGAGGTCCCGGTTCCACAGGAGATC**
**CCGGTGCCACAGGAGATCCCGGTGCCACAGGAGCTCCCGGTTCCACAGGAGCTCCCGGTGCCACA**
**GGAGCTCCCAGTGCCACAAGAGCTCCCGGTGCCACAGGAGCTCCCAGTGCCACAGGAGCTCCCGG**
**TGCCACAGGAGCTCCCGGTGCCACAGGAGCTCCCGGTGCCATTGGAGTTCCAGGAGGTACAGCAG**
**GCCCAGGCAGTGGGCTGGCGGGCACCCAAGGTACCTCCTCACTTCTGGCAGCCTGTGTCTGCCCAG**
**GAGGCCCAGGAGCAGGCCACTCAGATCGCCCATGTGGAGCAGCAGCAGCCCTTTCAGGGAGCTCC**
**AGCCTCCTCCAAGGCACTGCAAACTCAGCTGCCGACCCACCAGGCCCAAGCCTCTGGCTTGCAGGC**
**AGAACTGCCTTCAGTGCAGCTGCAGCCTTCTTGGCAAGGCCCACTGCCCATGTTGCAGGCCCAGCC**
**TGGAGCCTCTGCCACACTGGCAAACTTTCCCCGGGGCTCCACTCGATCACGTATGGCTCCATCAGG**
**AGAACCTGGCCCCTCTTCTCTAGAACCTCGGGGCCCTCCTAGGGAACGTAGGGCACCTGCAAGGG**
**ACAAAAAGGGTCCTCCAAAAGAGCGCATGTTCATTGGTGCCACTTTCTGTGCTCCAAGGGGGGCA**
**TCAGCATCCAGGGCATACGTGCCAACTGCCTGGAAAAACTTGCCTGCCACATCAGAGACCTTTCCT**
**GCCACCTCAAGGGTCTTTCCATCTACCTCTCATTTCCAGCCTGCCTCTTCTAATGCCTTTAGAGGTCC**
**ATCTGCCGCCTCAGAGAGCCCAAAGTCACTGCCATTTGCTCTGCAGGATCCTTATGCCTGCGTAGA**
**GGCCCTGCCTGCAGTTCCCTGGGTTCCGTATCCAGATGGAAATGCCTCATCAGCATGTAAGTCAGT**
**GCCTGCCATCTTGATGGTGGCAGCAGCTGCCCCCCAGGCAAGTGCCACTGCTGCAGAGGCCTCTAA**
**GTCTTCAGAGCCGCCAAGACGCCCCGGCAAAGCCACCAGGAAGAAGAAGCATCTGGAACCCAAA**
**GAAGACAACTGTGGCCACAGGCTCTCCTCACGTGACTGGCGGGGGCCCCGAACCTGGGGCAATCC**
**CAGTCATTCTGACTGGGAGATTCAGAGGGCTATGCAGCTCCTGGGGGACCGGGAATCCCTCTACA**
**CTCCGCAGGGCCTGAATGACTGGGGGTGCCCCAACACTTCTAGGATGCCAAGGAGCTTGGAGGGC**
**CCTAGCACTTCACGGGACCAGGAATTCTGTGGTGACTCGGGTGGGTCTCAGACATGGATGGCTTCT**
**GAGGTCCCAAGCGTCTCTCGGGGATCCAGTGCTGCTCAGGAGGACCCTGATAGGGAGAGTCAGCC**
**CTTATCTCCCTTAGATGAGAGAGCAAACGCTTTGGTGCAGTTTCTCTTGGTCAAAGACCAAGCCAA**
**GGTGCCTGTCCAGCTCTCGGAGATGGTAAATGTTGTCATCCGAGAATACAAAGACGACAGCTTAG**
**ACATCATCAACCGTGCCAACACTAAGCTGGAGTGCACCTTTGGTTGTCAACTGAAGGAAGTTGACA**
**CCAAAACCCACACTTACATCATCGTCAACAAGATGGCGTACCCTCAGTGTAATTTGCTGGCATCCTA**
**TTTAGAGAGGCCAAAGTTCAGCCTCCTGATGGTGGTCTTGAGCCTCATCTTTATGAAAGGCTACTG**
**TATCAGGGAGAATCTGCTCTTTAGTTTTCTGTTCCAGCTAGGGCTGGATGTCCAGGAGACAAGTGG**
**TCTCTTCAGAATTACAAAGAAGCTCATCACCAGTGTGTTTGTGAGACACAGGTACCTAGAGTACAG**
**GCAAATCCCGTTCACTGAGCCTGCAGAATACGAGCTTCTCTGGGGGCCCCGGGCATTCCTCGAAAC**
**CAACAGGGTGCACATCTTGAGATTTTTGGCCGCACTCTACGAGAACCAGCCCCAGATCTGGTCATG**
**CCAGTACCTTGACTCACTGGCAGAGTTAGAATACAAGGACGCAAATGCCGCCGCCGAAGAGTCCC**
**ATGACAGCGATGATGATGCCCACGACCCCACCAGCAGTCCCCATCCTCACTAA**TAGACGTTACTAA
GATGATTTTTATACTTGTGGGTCCAAAGCCAAAGGCAAAAGACAATGGGGGGTGGGGGACATTGTG
CTTTCTGGTGTTTTTTGGTATTGTTCTCTGTGTATAAATTTTACAGCCTGTTTTTATATTTGCCAAAGCT
TTTGTACAGTTTTGTGAAATGTTGATTTAAACTGGCTGCTGTATTGTTTTGTATTTTGGTGTCTGTATT
TTTTTAAATGAGATCTGTGATTCCCTGTTTGGTGTACTAATTGTAATGTTACAGTATCAGAGTTGTGCT
GCCTTGTGTGTGAAAAGAAATGACAGTTTATTTGTCTCGTTTTGTTTATGTAAAATCAGAATGTTTTT
GGATGTTAATATGACTCATCGGTAAAATACTGTCTGGAGAAAAGTAAAGATCTATACTATATAAATA
TACTTTTAGCATAACATAACGTGTCTGTCTTTATTCTAATTTTGAACCAGGGAGGTGGGAGCCACACT
TTAACATGCTTTACGCACAAACGCCAATATTACTATCAGTGATAGTAAAACCTGTGGATGCTGACAG

CTTCCCAGAAGAGTACTCCATTCATCCTAAAAGTGATGCAGTGATGGTGGGTGGTGAAGCCAGAGA
TCTCCTTAAAGTGAAAGTAAAACTTCCCAAGTAAGACGCGGCAAGCTAAGAGGGGGTGGGAGTCCC
CTCTGATTTAATTTTTACAATAACTGTTCAAGTTAGACATTGTATCTAGCTTGTAGGTGAGGCTATAA
GACACAAACAGGTTAATTAAAATACATATATGGTGGCATGCTGGTAAAATTACAATTCCAAGCAGGA
AATCTCAATGGTGTTTGGAAAAGGAGGGCACTTTCCCACAGGACCATAAAAATGCAGGTTTAGTAG
CCATACCAAACCACCAAAGAGATCCTTCTGTGCTGCATGTAGAACATGTGGAACACATGTGGAACAC
ATCAACACACTAAGGTGATACCAGATACCTGTGCCCTCTAATGTATCAGGATCCTACCAGGAAAAAG
AAATCAGGTTTAAGGTGTTAAGCATTCTGCAGATTACAGGTGGCCATTGCAAAGGGAGGGTGAGCC
GCTGCAGGGCACTTTCTTGGGAAACTTCTGAAAGAAAGTGGCTAAATACCTACAACCAGCCTTTAGT
TACAGGAACCATGGCAAAGGGCTGACCAGCCTGGCTGTTCTATTCAATGAATTCATGTGAATATGAA
TGTAATTATACATGAAAAGACTGGCTGAAGGTCACAGCTTTTAGACAGGGACATGAGGGTTCTGCTT
TTGCCACCTCTTGCTCTGTCTGAATCAAGAAACCCAACCCCATGTCTTCCCATTCTTCCTGAAAAGCAG
AGGGCAG

**MAGEL2-FLOX-EM1-B6**

AAACGGTCAGTCATCCTGGGGCATATTTCTTCTTTATGGCCCACCAGTTAAAAACGAACAACAACAA
CAAAACTATTTAGGAGCATGTGCTGAAGGTATGTTAAGAGTTGGGTACAGCCTTCAACATTCTCTTCT
GCCCCCTAGTGGTTTCTTTCCTCACCTTGGACGACTGCCCTAGGCTGCAAACCACTCCTTGTAAAGGA
ATCATAATGCTAGTTACCGTCTATAAACCACTAAACTTGACCTAAACTGAGATTGCCACTTAATTTTG
GGGAGAAAGAATACCAATATGCAGTAATTTTGATGGGTACCTCTTTATTTGTTAACATGGTATATGTC
CATTTTTTAGCTGATTTTCCAGAAAACTTTTCTTATACTCTCCATTTTTCTAGAACTAATTTCCAGCACA
TTGCTTCATTTTTTAATTATTGTTATTTTCAGGATGTCTTTCACACTCTATTGTCTCATCTCAC*atccgggg*
*gtaccgcgtcgag*GCGATCGCATAACTTCGTATAGCATACATTATACGAAGTTATTGAATTTCTGTTAATT
TACTTCCAAGGTTCCTTTTCTTTCCTCTCTAATTGGCTCGTCTTTTCTCTACCCGATTCTACGATGTAGT
CAGTATTCTTCTAGATGAGTTTAGATTTTCCCCCCTCTGAACACACTTTTAAACAATTGTATCCACTAC
TACTACTATGTTTTGATAAAGAATGGGCAAACTTGATTCTCGGATCCTCGTAATGAATACAGTTATAA
TAGACCACCTATGTTATCCCTGGGTTGACTGACTCATGTCCTAGCAAATATTCATCAACTCCCCTTACC
CTGCATGTGCTTCCTGCCCTTCAGTTACTTGACAGAATAAACTACTCCAGGAAATTCACACACCCGAG
ACAGATACCTGAATACAGTCATTTTTTTCTACACCTTCGTTAAATTGTAATAGCTGTCTCGAATAGCTC
CCCAGTCCCTCTTACGTCAATGTACTTCGTTTCTGAGCTCAGACCACACCACTACCCCCACCTTTGGAC
ATGTTCCATGTTGCTAGTGGTTTATTCCCACTTCATAATGCAGCACAGTGATTTTTTTTGAACCGTAAA
TCTGTCCCTATCACTTGCCTATATAAACTCTTGCTTAAAGAGTATGAACTAGCTTAGTTGAGGATCTAT
ACCGCAGGAGAAGGGATATTGAGTCAGTCGGGACTTTGGTAGACGCCCCTCTGAACAATCCACTTG
TGGTGGTCGGGGGCGGGGAAGGGGCGGGATTGAGCCCCCTCCCCCCTCCTTCTTACTGCGGAGAGC
TGCAGAGCCAGCCGACactctgtgtgagctgcgCTGCGAGCTGTGAGCTGAGAGCTGTGAGCTGCGTGGC
GGAGCAAGCCAGGCAGTAGCACTTGGCTGAAAGCTGCGGTGCTAGCCAGGCAGCGCTCGCTGAGA
GTTGCGGTGCCAGCCAGGTAGTGCTTGCTGAGAGCTGCTGAGAGAGCCCAGCTGCGCTCGCTGAGA
GCTGCGGAGAGAACCGCGGAACACGCCAGTCTGCACCGGCTTCCTATTCAGCCTTCAAGCTGGAGG
GGGATCCTGGACAGCCACGTCGGCATGCTCAGCACTGGAAGAAAAAGCGCAGCAACCGGAGACGG
GCCACCACCAGTCAAGCAAGGGAC**atgtcgcagctaagtacgaATCTGGGGGATTCCAGCCCTCCGGAGT**
**CCCCAGTGCCTGCAGTCCATAGCCGCCCTACGGTTCTGATGCGGGCTCCGCCTGCTTCCTCCCGGGC**
**TCCTCCTGTCCCCTGGGATCCACCTCCAGTGGACTTGCAGGCTCCCATGGCTGCTTGGCAGGCTCCT**
**CAACCTGCCTGGGAGGCTCCGGAGGGCCAGCTGCCTGCCCCAGTGGCTCAGCTGGCCCAGCCTCCT**
**GGTCTAGGGGCCCCAATGGTCCAGGCTCCACCGCTCGGAGGGGGGATGGCCAAGCCTCCAACTCC**
**TGGAGTCTTGATGGTCCATCAGCCCCCTCCGGGAGCCCCCATGGCCCAGTCTTCAACTCCGGGAGT**
**CCTGATGCTACATCCTTCTGTCACGGGGGCCCCTTTGGCTCATCCTCCTCCCCCAGGAACCCCGATG**

**ACACACCCTCCCGGGACCTCGATGGCGCACCCTCCTCCTCCTCCTCCCCCTCCTCCTCCTCCT
CCTGGGACCCCAATGACCCACCCACCTCCTCCTGGGACTCCGATGGGTCACCATCCTCCTCCTGGGA
ACCCTATGACCCATCCTCCACCGGGGAACCCGATGGTGCATCCTCTCACTCATGGAGCCCCGATGG
TCCATGGGGGACCACATGGAACTCCAATGCCGCATGTTCCCATTACGGGGACACCGATAGCCCAG
CAGCCAACTCCAGGAGTCCTGATGGCCCAGCAGCTGACACCGGGAGTCCTGATGGTCCAGCCGCC
TGCTCCGGGAGCTCCGATGGTCCAGCCACCTCCACAGGCTGCCTTGATGACCCAGCCTGCACCTTC
GATTACTCCGATGGCCAAGCCTCCAGGTCCTGGTGTCGTGATGATCCATCCTCCAGGTGCCAGAGG
TCCAATCATTCAGACTCCAGTATCAGGAGCACCAATGGCTCAGACAGTGCTGCCCCCTGGGCAGCC
TCTGGCCACTTGGGCCCCACAGGGTCAGCCTTTGATCCTACAAATCCAGTCTCAAGTCATAAGGGC
TCCTCCACAGGTTCCCTCTGTACCACAAGCCCCCCAGGTACAGCTGGCCACACCCCCAGGCTGGCAA
GCCACCACACCCAACTGGCAGGTGACCCCCCAGGGGTGGCCAGCAACGCCTTTGACATGGCAGGC
TACACAGGTGACCTGGCAGGCCCCCACAATAGCCTGGCAGGCCACTCAACCTGGGCGCCAAGGGC
ACTCAACCATTCGTACTGGTCACACACCCATTCGACCTGGACCAGCTCCATTGCTTCGCCAGATACC
TCCTATGATCCGTCAGATCCAACCTGTGATGAGGCAAGCCCCACCACTGATCCGACAGGTCCCTAT
CAGACCAGCTCCACATGGCATAGCAAGCCAGCCTCAGCTGTGGCAGGTCCTGCCACCCCCACCTCC
ACTGCGGCAGGCTCCACAGGCTCGTCTACTGCTCCCGAGGGTACCAGGAACAGGCCAGGTGTCTA
CAGTACCACCAGTTGCTCAGATACATTTGGTGCCACAGTCAGGCCCACAGGTGCCCCAGACAGTAC
TGCCAGCCCAACTGTCTATCCCAATTCCTGTACCCCAGGCTGCTGCTCAGTCTGCTCCCCGGACTGT
GCATTGCCCACCCATCATCTGGCAGGCCCCCAAAGGCCAGGCCCCGGTGCCACAGGAGCTCCCAGT
GCCACAGGAGCTCCCGGTGCCACAGGAGCTCCCGGTGCCACAAGAGGTCCCGGTTCCACAGGAGA
TCCCGGTGCCACAGGAGATCCCGGTGCCACAGGAGCTCCCGGTTCCACAGGAGCTCCCGGTGCCA
CAGGAGCTCCCAGTGCCACAAGAGCTCCCGGTGCCACAGGAGCTCCCAGTGCCACAGGAGCTCCC
GGTGCCACAGGAGCTCCCGGTGCCACAGGAGCTCCCGGTGCCATTGGAGTTCCAGGAGGTACAGC
AGGCCCAGGCAGTGGGCTGGCGGGCACCCAAGGTACCTCCTCACTTCTGGCAGCCTGTGTCTGCCC
AGGAGGCCCAGGAGCAGGCCACTCAGATCGCCCATGTGGAGCAGCAGCAGCCCTTTCAGGGAGC
TCCAGCCTCCTCCAAGGCACTGCAAACTCAGCTGCCGACCCACCAGGCCCAAGCCTCTGGCTTGCA
GGCAGAACTGCCTTCAGTGCAGCTGCAGCCTTCTTGGCAAGGCCCACTGCCCATGTTGCAGGCCCA
GCCTGGAGCCTCTGCCACACTGGCAAACTTTCCCCGGGGCTCCACTCGATCACGTATGGCTCCATCA
GGAGAACCTGGCCCCTCTTCTCTAGAACCTCGGGGCCCTCCTAGGGAACGTAGGGCACCTGCAAG
GGACAAAAAGGGTCCTCCAAAAGAGCGCATGTTCATTGGTGCCACTTTCTGTGCTCCAAGGGGGG
CATCAGCATCCAGGGCATACGTGCCAACTGCCTGGAAAAACTTGCCTGCCACATCAGAGACCTTTC
CTGCCACCTCAAGGGTCTTTCCATCTACCTCTCATTTCCAGCCTGCCTCTTCTAATGCCTTTAGAGGT
CCATCTGCCGCCTCAGAGAGCCCAAAGTCACTGCCATTTGCTCTGCAGGATCCTTATGCCTGCGTAG
AGGCCCTGCCTGCAGTTCCCTGGGTTCCGTATCCAGATGGAAATGCCTCATCAGCATGTAAGTCAG
TGCCTGCCATCTTGATGGTGGCAGCAGCTGCCCCCCAGGCAAGTGCCACTGCTGCAGAGGCCTCTA
AGTCTTCAGAGCCGCCAAGACGCCCCGGCAAAGCCACCAGGAAGAAGAAGCATCTGGAACCCAA
AGAAGACAACTGTGGCCACAGGCTCTCCTCACGTGACTGGCGGGGGCCCCGAACCTGGGGCAATC
CCAGTCATTCTGACTGGGAGATTCAGAGGGCTATGCAGCTCCTGGGGGACCGGGAATCCCTCTAC
ACTCCGCAGGGCCTGAATGACTGGGGGTGCCCCAACACTTCTAGGATGCCAAGGAGCTTGGAGGG
CCCTAGCACTTCACGGGACCAGGAATTCTGTGGTGACTCGGGTGGGTCTCAGACATGGATGGCTTC
TGAGGTCCCAAGCGTCTCTCGGGGATCCAGTGCTGCTCAGGAGGACCCTGATAGGGAGAGTCAGC
CCTTATCTCCCTTAGATGAGAGAGCAAACGCTTTGGTGCAGTTTCTCTTGGTCAAAGACCAAGCCA
AGGTGCCTGTCCAGCTCTCGGAGATGGTAAATGTTGTCATCCGAGAATACAAAGACGACAGCTTA
GACATCATCAACCGTGCCAACACTAAGCTGGAGTGCACCTTTGGTTGTCAACTGAAGGAAGTTGAC
ACCAAAACCCACACTTACATCATCGTCAACAAGATGGCGTACCCTCAGTGTAATTTGCTGGCATCCT
ATTTAGAGAGGCCAAAGTTCAGCCTCCTGATGGTGGTCTTGAGCCTCATCTTTATGAAAGGCTACT
GTATCAGGGAGAATCTGCTCTTTAGTTTTCTGTTCCAGCTAGGGCTGGATGTCCAGGAGACAAGTG**

**GTCTCTTCAGAATTACAAAGAAGCTCATCACCAGTGTGTTTGTGAGACACAGGTACCTAGAGTACA GGCAAATCCCGTTCACTGAGCCTGCAGAATACGAGCTTCTCTGGGGGCCCCGGGCATTCCTCGAAA CCAACAGGGTGCACATCTTGAGATTTTTGGCCGCACTCTACGAGAACCAGCCCCAGATCTGGTCAT GCCAGTACCTTGACTCACTGGCAGAGTTAGAATACAAGGACGCAAATGCCGCCGCCGAAGAGTCC CATGACAGCGATGATGATGCCCACGACCCCACCAGCAGTCCCCATCCTCACTAA**TAGACGTTACTAA GATGATTTTTATACTTGTGGGTCCAAAGCCAAAGGCAAAAGACAATGGGGGGTGGGGACATTGTG CTTTCTGGTGTTTTTTGGTATTGTTCTCTGTGTATAAATTTTACAGCCTGTTTTTATATTTGCCAAAGCT TTTGTACAGTTTTGTGAAATGTTGATTTAAACTGGCTGCTGTATTGTTTTGTATTTTGGTGTCTGTATT TTTTTAAATGAGATCTGTGATTCCCTGTTTGGTGTACTAATTGTAATGTTACAGTATCAGAGTTGTGCT GCCTTGTGTGTGAAAAGAAATGACAGTTTATTTGTCTCGTTTTGTTTATGTAAAATCAGAATGTTTTT GGATGTTAATATGACTCATCGGTAAAATACTGTCTGGAGAAAAGTAAAGATCTATACTATATAAATA TACTTTTAGCATAACATAACGTGTCTGTCTTTATTCTAATTTTGAACCAGGGAGGTGGGAGCCACACT TTAACATGCTTTACGCACAAACGCCAATATTACTATCAGTGATAGTAAAACCTGTGGATGCTGACAG CTTCCCAGAAGAGTACTCCATTCATCCTAAAAGTGATGCAGTGATGGTGGGTGGTGAAGCCAGAGA TCTCCTTAAAGTGAAAGTAAAACTTCATAACTTCGTATAGCATACATTATACGAAGTTAT*CGCCGGCG ggtctgagctcgccatcagt*GAGTCCCCTCTGATTTAATTTTTACAATAACTGTTCAAGTTAGACATTGTATC TAGCTTGTAGGTGAGGCTATAAGACACAAACAGGTTAATTAAAATACATATATGGTGGCATGCTGGT AAAATTACAATTCCAAGCAGGAAATCTCAATGGTGTTTGGAAAAGGAGGGCACTTTCCCACAGGAC CATAAAAATGCAGGTTTAGTAGCCATACCAAACCACCAAAGAGATCCTTCTGTGCTGCATGTAGAAC ATGTGGAACACATGTGGAACACATCAACACACTAAGGTGATACCAGATACCTGTGCCCTCTAATGTA TCAGGATCCTACCAGGAAAAAGAAATCAGGTTTAAGGTGTTAAGCATTCTGCAGATTACAGGTGGC CATTGCAAAGGGAGGGTGAGCCGCTGCAGGGCACTTTCTTGGGAAACTTCTGAAAGAAAGTGGCTA AATACCTACAACCAGCCTTTAGTTACAGGAACCATGGCAAAGGGCTGACCAGCCTGGCTGTTCTATT CAATGAATTCATGTGAATATGAATGTAATTATACATGAAAAGACTGGCTGAAGGTCACAGCTTTTAG ACAGGGACATGAGGGTTCTGCTTTTGCCACCTCTTGCTCTGTCTGAATCAAGAAACCCAACCCCATGT CTTCCCATTCTTCCTGAAAAGCAGAGGGCAG

LoxP sites are in red and underlined and genotyping handles (restriction enzyme site plus primer unique to each LoxP site) are in red and italics. Floxed exon highlighted in bold.

##### QC strategy employed at Harwell to check the edited allele:

Genomic DNA was extracted from ear clip biopsies and amplified in a PCR reaction using the following conditions/primer sequences:

| | |
|---|---|
| Geno_Magel2_F1 (5’-3’) | AAACGGTCAGTCATCCTGGG |
| Geno_Magel2_R1 (5’-3’) | CAGAGGGGCGTCTACCAAAG |
| Taq Polymerase used | Roche Expand Long Range DNTPack |
| Annealing Temperature (°C) | 63 |
| Elongation time (min) | 1.5 |
| WT product size (bp) | 1206 |
| Mutant product size (bp) | 1225 |
| Notes | This targets the 5’ LoxP insertion site. |

| | |
|---|---|
| Geno_Magel2_F2 (5'-3') | GAGTTGTGCTGCCTTGTGTG |
| Geno_Magel2_R2 (5'-3') | CTGCCCTCTGCTTTTCAGGA |
| Taq Polymerase used | Roche Expand Long Range DNTPack |
| Annealing Temperature (°C) | 63 |
| Elongation time (min) | 1.5 |
| WT product size (bp) | 1090 |
| Mutant product size (bp) | 1119 |
| Notes | This targets the 3' LoxP insertion site. |

All amplicons were sent for Sanger sequencing to check for integration of the donor oligo sequence at the target site. F1 sequences should be heterozygous unless on sex chromosome.

**Additional integrations of the donor sequence**
Copy counting of the donor sequence was carried out by ddPCR at the F1 stage to confirm donor oligos were inserted once on target into the genome. The following Taqman assay was used to copy count the donor sequence compared against a VIC-labelled reference assay for Dot1l:

| | |
|---|---|
| Assay name | Magel2-CR-LOA-WT1 |
| Forward Primer (5'-3') | GGTCCTGGTGTCGTGATGA |
| Reverse Primer (5'-3') | GCCATTGGTGCTCCTGATAC |
| Probe (5'-3') | TCCATCCTCCAGGTGCCAGAGG |
| Label | FAM |

This ddPCR assay is universal; both the WT and mutant alleles are recognised by this assay. Therefore, WT controls are expected to call at 2 copies and a single integration for a correct mutation is expected to call at 2 copies for F1 (HET) animals.

| | |
|---|---|
| Assay name | Magel2-FLOX-5'-MUT1 |
| Forward Primer (5'-3') | CAGGATGTCTTTCACACTCTATTGTC |
| Reverse Primer (5'-3') | GGGTAGAGAAAAGACGAGCCAATTAG |
| Probe (5'-3') | TCGAGGCGATCGCATAACTTCG |
| Label | FAM |

This ddPCR assay is specific to the donor used to create the engineered mutation and only mutant alleles are expected to be recognised by this assay. Therefore, WT controls are expected to call at 0 copies and a single integration for a correct mutation is expected to call at 1 copy for F1 (HET) animals.

| | |
|---|---|
| Assay name | Magel2-FLOX-3'-MUT1 |
| Forward Primer (5'-3') | TGGGTGGTGAAGCCAGAGAT |
| Reverse Primer (5'-3') | GCCTCACCTACAAGCTAGATACAATG |
| Probe (5'-3') | AAGTTATCGCCGGCGGGTCTGA |
| Label | FAM |

This ddPCR assay is specific to the donor used to create the engineered mutation and only mutant alleles are expected to be recognised by this assay. Therefore, WT controls are expected to call at 0 copies and a single integration for a correct mutation is expected to call at 1 copy for F1 (HET) animals.

| | |
|---|---|
| Reference Assay Name | Dot1l |
| Forward primer (5'-3') | GCCCCAGCACGACCATT |
| Reverse primer (5'-3') | TAGTTGGCATCCTTATGCTTCATC |
| Probe (5'-3') | CCCAACAGGCCTGGATTCTCAATGC |
| Label | VIC |

VIC-labelled reference assay for Dot1l gene.

No additional donor integrations were detected in the animals taken forward to establish the colony.

MRC Harwell

### Large deletion: PCAN-DEL3Mbp-EM4-B6

**Name of Mouse model or mutation:**
**PCAN-DEL3Mbp-EM4-B6**

**Description:**
**PCAN-DEL3Mbp-EM4-B6:** CRISPR/Cas9 induced mutation. Deletion of 3 mega base pairs from 7:59336000-59337600 (between Ube3a and snoRNAs) and 7:62423586-62427187 (between Mkm3 and Peg12).

**Type of mutation:**
**PCAN-DEL3Mbp-EM4-B6:** Deletion of 3,089,372 nt from Chromosome 7.

**Delivery method:**
Cytoplasmic injection into 1-cell stage embryo.

**Genetic Background:**
C57BL/6J

**Nuclease:**
Cas9 mRNA

**sgRNAs:**

| Protospacer sequence | PAM sequence |
|---|---|
| TAAGGCTAATGTCTGCCCAT | TGG |
| GAGACGACTTGCTTGTATAA | AGG |

**Pronuclear Microinjection mixes:**
Microinjection buffer (MIB; 10 mM Tris–HCl, 0.1 mM EDTA, 100 mM NaCl, pH7.5) was prepared and filtered through a 2 nm filter and autoclaved. Cas9 mRNA and sgRNAs were diluted and mixed in MIB to the working concentrations of 50 ng/µl and 6.5 ng/µl each, respectively. Injected embryos were re-implanted in CD1 pseudo-pregnant females. Host females were allowed to litter and rear $F_0$ progeny.

**Sequence details**
**PCAN-DEL3Mbp-EM4-B6:**

TCTCTAAACATGATGCTAATGCTGATATTTTATTCATCAGAAATAACAAAAGCAAAAATTAAAGAAATCAATTT
GCAAAAGAATAAATAGCTATTATAATAAACTAATACAGTTGTATTTAGGTATAATTTGTATCTGTTTTCCCATG
GAACTTTCTCAATTTTACTGATAGTTACGTTCATCTTTATTACCAGTTGATCATGTTTTATTTCTATATATTTCTGT
AACCAAGTATTCACGCAATACTTGGATGTAACCAATATCAAACTGTTCACAGTAATTAAAATATCAAACTTCTC
CTTAATGTGAAACCTTAAGGCTAATGTCTG **[3 Mbp (3,089,372 bp)  deletion]**
TAAAGGCACAGTCTGGACATATGAAACACTAGGGGTATAAAATTTCATATCTGAACA **[10 nt deletion]**
TTTTTAGATTTCAAGTCAATGTATTTGTGAATTTTTTCCTGTTAATAGATTTTTTTGCCTCACAGAAAAATTTACA
CTACATAATCTAACCTCCCACAAATATGGTTTTTATACTTTTGTTTTGAAGTCTTCATGCATTCTCTGAGAATTCA
ACTCAATGCCTTTGGATTATATTCATCTCCTTTCCCAGCTTCTCCC

**Primers/ assays used to QC the allele:**

| | |
|---|---|
| Geno_PCAN_LD_F1 primer (5'-3') | TCTCTAAACATGATGCTAATGCTG |
| Geno_PCAN_LD_R1 primer (5'-3') | GGGAGAAGCTGGGAAAGGAG |
| Annealing Temperature (°C) | 58°C |
| Elongation time (min) | 1 min |
| WT product size (bp) | Too large to PCR – 3 Mb |
| Mutant product size (bp) | ~600 nt |
| Sequence with primers (if different to amplification primers) | |
| Notes | This is a very large deletion encompassing many genes. We performed copy counting assays for those genes to determine if there had been a deletion. |

**Assays used to copy count for the large deletion:**
Copy counting of the deleted region was carried out by ddPCR at the F1 stage to confirm there had been a deletion. The following assay was used to copy count the donor sequence compared against a VIC-labelled reference assay for Dot1l:

| | |
|---|---|
| Assay Name | Snrpn_UPL21 |
| Forward primer | tgtgtggaaatgtggagtgtc |
| Reverse primer | ataattctgtaccacggtgtatgc |
| Probe | UPL21 |
| Annealing temperature | 58C |
| Notes | The ddPCR assays detailed above sit within the 30Mb region which we are trying to delete. Therefore WT controls are expected to call at 2 copies and a correct deletion is expected to call at 1 copy for F1 (HET) animals. |

| | |
|---|---|
| Assay Name | Snurf-201_UPL5 |
| Forward primer | gtcatttctgcggtgttgg |
| Reverse primer | ctcagttctccagggcctta |
| Probe | UPL5 |
| Annealing temperature | 58C |
| Notes | The ddPCR assays detailed above sit within the 30Mb region which we are trying to delete. Therefore WT controls are expected to call at 2 copies and a correct deletion is expected to call at 1 copy for F1 (HET) animals. |

| | |
|---|---|
| Assay Name | Ipw-CR-LOA-WT1 |
| Forward primer | TTGAAGGCTATCCCCATAGTTTC |
| Reverse primer | CCCCACCAGACAGTATGTGTATC |
| Probe | TCTGTTAAGAGAAGGATATACATGC |
| Annealing temperature | 58C |
| Notes | The ddPCR assays detailed above sit within the 30Mb region which we are trying to delete. Therefore WT controls are expected to call at 2 copies and a correct deletion is expected to call at 1 copy for F1 (HET) animals. |

| | |
|---|---|
| Assay Name | Magel2-CR-LOA-WT1 |
| Forward primer | GGTCCTGGTGTCGTGATGA |
| Reverse primer | GCCATTGGTGCTCCTGATAC |
| Probe | TCCATCCTCCAGGTGCCAGAGG |
| Annealing temperature | 58C |
| Notes | The ddPCR assays detailed above sit within the 30Mb region which we are trying to delete. Therefore WT controls are expected to call at 2 copies and a correct deletion is expected to call at 1 copy for F1 (HET) animals. |

| | |
|---|---|
| Assay Name | Ndn-CR-LOA-WT1 |
| Forward primer | GCACCTGAGGCTGACCAAT |
| Reverse primer | TGGGCTGAGGGCTTTGAC |
| Probe | TCCACACCATGGAGTTTGCCCTG |
| Annealing temperature | 58C |
| Notes | The ddPCR assays detailed above sit within the 30Mb region which we are trying to delete. Therefore WT controls are expected to call at 2 copies and a correct deletion is expected to call at 1 copy for F1 (HET) animals. |

| | |
|---|---|
| Assay Name | PCAN3'_UPL39 |
| Forward primer | TGTTGCCATGGTACAGACTCA |
| Reverse primer | TTTCCCTCTCTTGTGCAGTG |
| Probe | UPL39 |
| Annealing temperature | 58C |
| Notes | The ddPCR assays detailed above sit within the 30Mb region which we are trying to delete. Therefore WT controls are expected to call at 2 copies and a correct deletion is expected to call at 1 copy for F1 (HET) animals. |

| | |
|---|---|
| Assay Name | PCAN3'_UPL12 |
| Forward primer | GGCAGACTTGCACCTCTCTAC |
| Reverse primer | CCATTTGAAAATAGAAGAGTGATCG |
| Probe | UPL12 |
| Annealing temperature | 58C |
| Notes | The ddPCR assays detailed above sit within the 30Mb region which we are trying to delete. Therefore WT controls are expected to call at 2 copies and a correct deletion is expected to call at 1 copy for F1 (HET) animals. |

| | |
|---|---|
| Reference Assay Name | Dot1l |
| Forward primer | GCCCCAGCACGACCATT |
| Reverse primer | TAGTTGGCATCCTTATGCTTCATC |
| Probe | CCCAACAGGCCTGGATTCTCAATGC |
| Annealing temperature (°C) | 58 |
| Notes | Vic labelled reference assay specific to Dot1l on chromosome 10. |
